# AI-guided pipeline for protein-protein interaction drug discovery identifies a SARS-CoV-2 inhibitor

**DOI:** 10.1101/2023.06.14.544560

**Authors:** Philipp Trepte, Christopher Secker, Simona Kostova, Sibusiso B. Maseko, Soon Gang Choi, Jeremy Blavier, Igor Minia, Eduardo Silva Ramos, Patricia Cassonnet, Sabrina Golusik, Martina Zenkner, Stephanie Beetz, Mara J. Liebich, Nadine Scharek, Anja Schütz, Marcel Sperling, Michael Lisurek, Yang Wang, Kerstin Spirohn, Tong Hao, Michael A. Calderwood, David E. Hill, Markus Landthaler, Julien Olivet, Jean-Claude Twizere, Marc Vidal, Erich E. Wanker

**Affiliations:** Proteomics and Molecular Mechanisms of Neurodegenerative Diseases, Max Delbrück Center for Molecular Medicine in the Helmholtz Association, 13125, Berlin, Germany; Brain Development and Disease, Institute of Molecular Biotechnology of the Austrian Academy of Sciences, 1030, Vienna, Austria; Zuse Institute Berlin, Berlin, Germany; Laboratory of Viral Interactomes, Interdisciplinary Cluster for Applied Genoproteomics (GIGA)-Molecular Biology of Diseases, University of Liège, 4000, Liège, Belgium; Center for Cancer Systems Biology (CCSB), Dana-Farber Cancer Institute, Boston, MA, 02215, USA; Department of Genetics, Blavatnik Institute, Harvard Medical School, Boston, MA, 02115, USA; Department of Cancer Biology, Dana-Farber Cancer Institute, Boston, MA, 02215, USA; RNA Biology and Posttranscriptional Regulation, Max Delbrück Center for Molecular Medicine in the Helmholtz Association, Berlin Institute for Medical Systems Biology, 13125, Berlin, Germany; Département de Virologie, Unité de Génétique Moléculaire des Virus à ARN (GMVR), Institut Pasteur, Centre National de la Recherche Scientifique (CNRS), Université de Paris, Paris, France; Protein Production & Characterization, Max Delbrück Center for Molecular Medicine in the Helmholtz Association, 13125, Berlin, Germany; Multifunctional Colloids and Coating, Fraunhofer Institute for Applied Polymer Research (IAP), 14476, Potsdam-Golm, Germany; Structural Chemistry and Computational Biophysics, Leibniz-Institut für Molekulare Pharmakologie (FMP), 13125, Berlin, Germany; Institute of Biology, Humboldt-Universität zu Berlin, 13125, Berlin, Germany; Structural Biology Unit, Laboratory of Virology and Chemotherapy, Rega Institute for Medical Research, Department of Microbiology, Immunology and Transplantation, Katholieke Universiteit Leuven, 3000, Leuven, Belgium; TERRA Teaching and Research Center, Gembloux Agro-Bio Tech, University of Liège, 5030, Gembloux, Belgium; Laboratory of Algal Synthetic and Systems Biology, Division of Science and Math, New York University Abu Dhabi, Abu Dhabi, United Arab Emirates

**Author notes:** These authors contributed equally: Philipp Trepte, Christopher Secker. These authors jointly supervised this work (Julien Olivet, Jean-Claude Twizere, Marc Vidal, Erich E. Wanker).

**Keywords:** protein-protein interactions, machine learning, AlphaFold, VirtualFlow, SARS-CoV-2

## Abstract

Protein-protein interactions (PPIs) offer great opportunities to expand the druggable proteome and therapeutically tackle various diseases, but remain challenging targets for drug discovery. Here, we provide a comprehensive pipeline that combines experimental and computational tools to identify and validate PPI targets and perform early-stage drug discovery. We have developed a machine learning approach that prioritizes interactions by analyzing quantitative data from binary PPI assays and AlphaFold-Multimer predictions. Using the quantitative assay LuTHy together with our machine learning algorithm, we identified high-confidence interactions among SARS-CoV-2 proteins for which we predicted three-dimensional structures using AlphaFold Multimer. We employed VirtualFlow to target the contact interface of the NSP10-NSP16 SARS-CoV-2 methyltransferase complex by ultra-large virtual drug screening. Thereby, we identified a compound that binds to NSP10 and inhibits its interaction with NSP16, while also disrupting the methyltransferase activity of the complex, and SARS-CoV-2 replication. Overall, this pipeline will help to prioritize PPI targets to accelerate the discovery of early-stage drug candidates targeting protein complexes and pathways.

## INTRODUCTION

Enzymes, ion channels and receptors are among the most favored proteins for target-based drug discovery (Santos et al, 2017). However, the number of newly approved drugs per billion dollars invested per year has decreased in the last 60 years (Scannell et al, 2012; Ringel et al, 2020), and the currently approved small molecules target less than 700 proteins altogether or approximately 3% of the human protein-coding genome (Harding et al, 2018). Proteins are part of signaling pathways and multisubunit complexes (Vidal et al, 2011), thus their macromolecular interactions such as protein-DNA and protein-protein interactions (PPIs) are key targets to expand the druggable proteome (Makley & Gestwicki, 2013; Lu et al, 2020). Consequently, characterizing molecular complex interactions and direct contacts between constitutive protein subunits is essential to identify new classes of targets for drug discovery and development.

Affinity purification coupled to mass spectrometry (AP-MS) techniques are highly efficient in identifying the composition of protein complexes at proteome-scale (Huttlin et al, 2021; Bludau & Aebersold, 2020), while binary PPI assays such as yeast two-hybrid (Y2H) can provide high-quality information about directly interacting protein subunits (Luck et al, 2020). Structural biology technologies, and in particular cryo-electron microscopy (cryo-EM), can capture near atomic resolution pictures of complexes purified from native sources (Costa et al, 2017; Callaway, 2020). Also, they can provide information on the precise assembly of subunits and the organization of their interaction interfaces. However, out of the ∼7,000 protein complexes that have been found in the human proteome (Drew et al, 2021), only ∼4% of them currently have an experimentally resolved structure in the literature, which calls for complementary approaches to rapidly model subunit-subunit interactions.

Computational predictions are on the rise to help address this challenge. On the one hand, predictions of 3D protein structures based on artificial intelligence (AI) strategies such as those available in AlphaFold and RoseTTAFold (Jumper et al, 2021; Baek et al, 2021) can be exploited to model protein assemblies and interaction interfaces with much improved accuracies than previous computational tools (Evans et al, 2022; Gao et al, 2022). On the other hand, platforms like VirtualFlow can be used to screen billions of molecules *in silico* against the predicted target in a time- and cost-effective manner (Gorgulla et al, 2020).

Here, we combine experimental binary PPI mapping with *in silico* structure prediction and virtual screening for PPI-based drug discovery. We first used reference sets of PPIs and quantitative interaction data from seven binary PPI assays to establish an unbiased machine-learning PPI scoring approach. We then applied this strategy to map and prioritize interactions between SARS-CoV-2 proteins and used AlphaFold to determine the corresponding 3D protein complex structures. Finally, we targeted the contact interface of the NSP10-NSP16 complex in an ultra large virtual drug screening with VirtualFlow and identified a small molecule PPI inhibitor that disrupts the NSP16-linked methyltransferase activity and SARS-CoV-2 replication. Our findings show that combining high-quality quantitative binary interaction data, AI-based scoring systems, and computational modeling can help prioritizing PPI targets for the development of novel therapeutics.

## RESULTS

### Scoring binary interaction assays using fixed cutoffs results in variable recovery rates

We previously demonstrated that combining multiple complementary interaction assays and/or versions thereof significantly increases PPI recovery while maintaining high specificity (Venkatesan et al, 2009; Choi et al, 2019; Trepte et al, 2018). LuTHy, a bioluminescence-based technology (Trepte et al, 2018) combines two readouts in one go. First, a bioluminescence resonance energy transfer (BRET)-based readout is used to quantify interactions in living cells (LuTHy-BRET; **Figure EV1A**); then, cells are lysed and luminescence is used to quantify interactions after protein co-precipitation (LuTHy-LuC; **Figure EV1A**). Since LuTHy plasmids allow expression of each protein as N- or C-terminal fusions, and as donor (NanoLuc tag or NL) or acceptor (mCitrine tag or mCit) proteins, eight tagging configurations can be assessed for every protein pair of interest (**Figure EV1B**). Thus, when all eight configurations are tested, LuTHy-BRET and LuTHy-LuC assays generate a total of 16 data points for every tested X-Y pair.

To determine the accuracy of the LuTHy assay and compare it to other binary interaction assays (Choi et al, 2019; Yao et al, 2020), we tested an established positive reference set (PRS), hsPRS-v2, which contains 60 well-characterized human PPIs (Venkatesan et al, 2009; Choi et al, 2019). To control for specificity, a random reference set (RRS), hsRRS-v2, made of 78 pairs of human proteins not known to interact (Choi et al, 2019), was also tested (**Source Data Figure 1**). To coherently score quantitative PPI data among different readouts and assays, we initially tried two different approaches: i) we applied a receiver operating characteristic (ROC) analysis to determine cutoffs at maximal specificity (i.e. under conditions where none of the random protein pairs from hsRRS-v2 are scored positive in any of the tested configurations, (**Figure EV1C**), and ii) we determined cutoffs based on the distribution of the data at the mean (**Figure EV1D**) or median plus one standard deviation (**Figure EV1E**). To analyze the reproducibility of such fixed cutoffs, we combined the positive (hsPRS-v2) and negative (hsRRS-v2) reference sets and randomly split them twenty times into a training and a test set (**Figure EV2A**). Next, for each of these paired training and test sets, we used the LuTHy data of the training set to determine a cutoff under conditions of maximal specificity and applied it to the test set to determine the recovery rates of the two LuTHy readouts. Interestingly, depending on the subset of reference interactions, we obtained highly variable recovery rates in the test set with LuTHy-BRET or LuTHy-LuC detecting on average 27±15% or 47±10% of hsPRS-v2 interactions, while recovering on average 2±2% or 2±3% of hsRRS-v2 protein pairs, respectively (**Figure EV2B,C**). This shows that fixed cutoffs can result in highly variable recovery rates, highlighting the need for more robust and coherent approaches to score quantitative PPI data.

**Figure 1.**
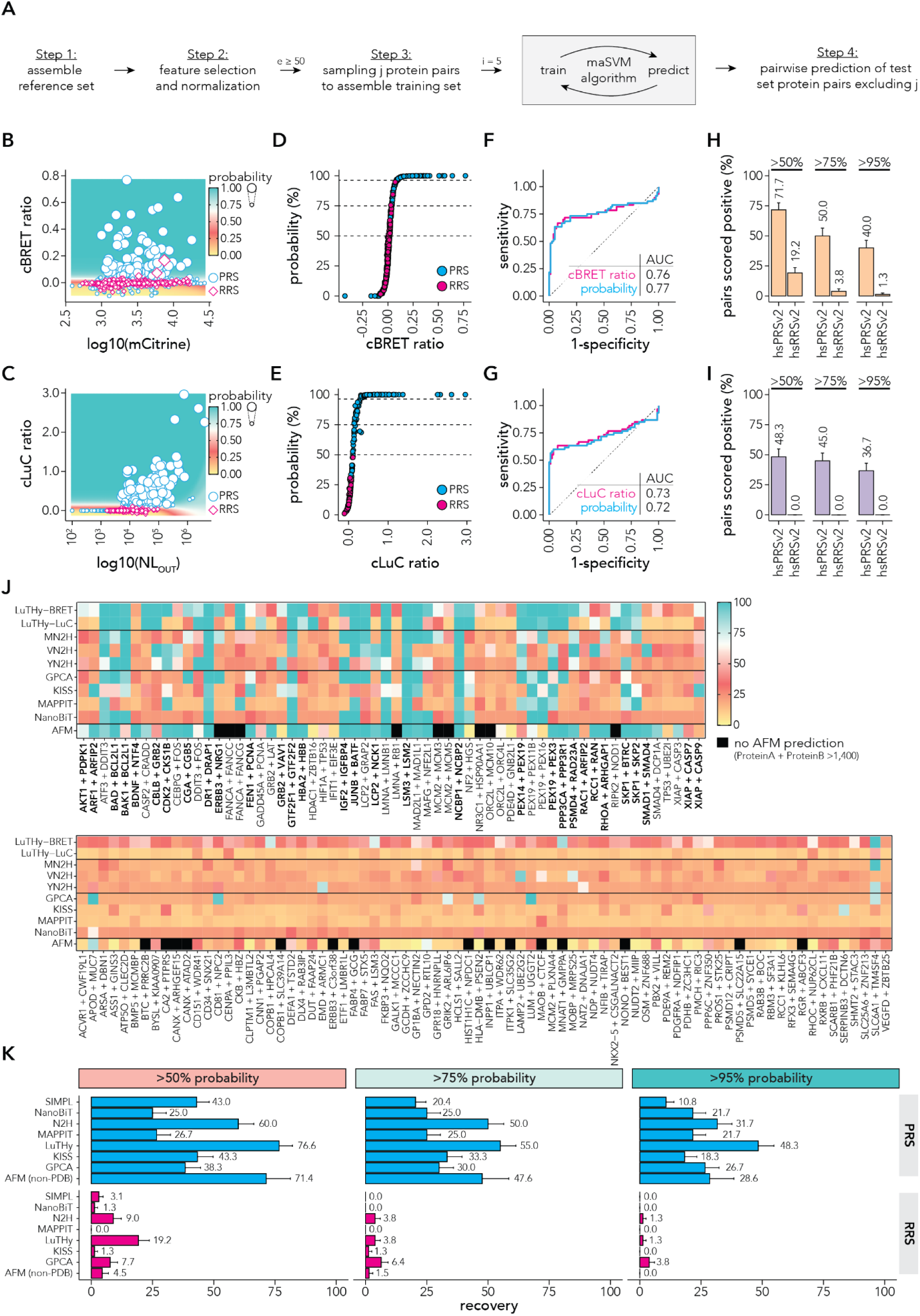
Developing a maSVM algorithm to classify protein pairs from hsPRS-v2 and hsRRS-v2 using the LuTHy assay. (**A**) Schematic overview of the maSVM learning algorithm. Step 1: assembly of reference set; Step 2: feature selection and RobustScaler normalization for reference and test set; Step 3: assembly of ‘e’ training sets (ensembles) by weighted sampling j protein pairs from the reference set to train ‘e≥50’ maSVM algorithms, where the training classifier labels are reclassified in ‘i=5’ iterations; Step 4: prediction of test set protein pairs excluding training set using the paired maSVM model. If classifier probabilities were not predicted for all test set protein pairs in ‘e=50’ ensembles, the maSVM algorithm was repeated from Step 3 assembling additional ‘e=10’ training sets excluding protein pairs from the training set that lack classifier probabilities. (**B,C**) Scatter plot showing (**B**) in-cell mCitrine expression (x-axis) against cBRET ratios (y-axis limited to ‘> −0.1’) or (**C**) luminescence after co-precipitation (NL_OUT_) (x-axis) against cLuC ratios (y-axis) for all hsPRS-v2 (blue) and hsRRS-v2 (magenta) protein pairs from all eight tagging configurations. Average classifier probabilities from the 50 maSVM models are displayed as the size of the data points and as a colored grid in the background. (**D,E**) Scatter plot showing (**D**) cBRET ratios (x-axis) or (**E**) cLuC ratios (x-axis) against classifier probability (y-axis) for all hsPRS-v2 (blue) and hsRRS-v2 (magenta) protein pairs from all eight tagging configurations. (**F,G**) Receiver characteristic analysis comparing sensitivity and specificity between (**F**) cBRET ratios or (**G**) cLuC ratios and classifier probabilities. The calculated areas under the curve are displayed. (**H,I**) Bar plots showing the fraction of hsPRS-v2 and hsRRS-v2 protein pairs that scored above classifier probabilities of 50%, 75% or 95% by (**H**) LuTHy-BRET or (**I**) LuTHy-LuC. Only the highest classifier probability per tested tagging configuration is considered. (**J**) Heatmaps showing the highest classifier probabilities for the hsPRS-v2 (top) and hsRRS-v2 (bottom) protein pairs per tested tagging configuration. hsPRS-v2 interactions supported by structures or homologous structures are highlighted in bold. LuTHy and AFM data from this study; all other from Choi et al (Choi et al, 2019). (**K**) Bar plots showing the fraction of hsPRS-v2 and hsRRS-v2 protein pairs that scored above classifier probabilities of 50%, 75% or 95% for 10 binary PPI assay versions. Only the highest classifier probability per tested tagging configuration is considered. For AFM, the fraction of hsPRS-AF with no experimental structure or homologous structures (non-PDB) is shown (see Figure EV4D for recovery rate of all hsPRS-AF interactions). LuTHy and AFM data from this study; SIMPL from Yao et al (Yao et al, 2020); all other from Choi et al (Choi et al, 2019). Note that the SIMPL assay was benchmarked by Yao et al against 88 positive proteins pairs derived from the hsPRS-v1 (Venkatesan et al, 2009) and as a random reference set against “88 protein pairs with baits and preys selected from the PRS but used in combinations determined computationally to have low probability of interaction” (Yao et al, 2020).

### Establishing a machine learning algorithm to classify binary interactions

To provide a universal and unbiased approach to score quantitative PPI data from various assays, we aimed to implement a support vector machine (SVM) learning algorithm, which is commonly used for binary classifications (Chang & Lin, 2011). For datasets that contain a high degree of mislabeling, it has been shown that a multi-adaptive sampling approach can be used to iteratively update the labeling class (positive or negative) of the training data and thereby improve the performance of the SVM learning algorithm (Yang et al, 2017). We rationalized that this approach could be applied to analyze quantitative PPI datasets, in which frequently, a positive reference set interaction is labeled positive, but will not score positive in a certain assay or tagging configuration and should therefore be relabeled as negative (for this specific assay or tagging orientation) during the training of the SVM.

To test if such a multi-adaptive SVM learning (maSVM) algorithm could be used to classify quantitative PPI data, we first applied it to the LuTHy assay. We used the reference sets that we had randomly split into 20 paired training and test sets (**Figure EV2A**), which consisted of 96 training and 42 test interactions. Since each interaction was tested in eight tagging configurations, the training set consisted of quantitative interaction data from 768 pairwise combinations (i.e. 96 interactions x 8 configurations). For each of the 20 training sets, we randomly sampled 90 of the 768 protein pairs, and used for the LuTHy-BRET the cBRET ratios and acceptor fluorescence (mCit) and for the LuTHy-LuC the cLuC and luminescence after co-precipitation (NL_OUT_) as training features for the maSVM algorithms (see methods for details). During training of each maSVM model, the classifier label of each reference interaction was iteratively reclassified five times. In total, the sampling and training was repeated 50 times, resulting in 50 maSVM models that were each used to predict the classification probabilities of all protein pair combinations of the test set (**Figure 1A and Figure EV2D-G**). Since we had split the reference set 20 times (**Figure EV2A)**, we generated maSVM models for each training set, which were then used to predict the average classification probabilities of the 20 test sets in a paired fashion (**Figure EV2H,I**). Finally, we calculated for each paired training and test set the recovery rates at >50%, >75% and >95% interaction probabilities (**Figure EV2J,K**). On average, 29±8% of hsPRS-v2 PPIs and 1±2% of hsRRS-v2 pairs were recovered with >95% interaction probability by LuTHy-BRET in the test sets. Thus, similar recovery rates were obtained by this approach as compared to scoring with fixed cutoffs (hsPRS-v2: 27±15%; hsRRS-v2: 2±2%, **Figure EV2B**) but with a significantly lower variance (F-test, p = 0.009). Indeed, for LuTHy-LuC, only 30±7% of hsPRS-v2 and 0±0% of hsRRS-v2 pairs were recovered with >95% interaction probability from the test sets, which was lower than when applying fixed cutoffs (hsPRS-v2: 47±10%; hsRRS-v2: 2±3%; F-test, p = 0.586, **Figure EV2B**). However, for LuTHy-LuC at >50% probability a similar recovery rate of 44±10% was obtained compared to the fixed cutoffs, suggesting that some protein pairs above the fixed cutoffs show very similar interaction scores as random protein pairs and have thus relatively low probability to be detected as true positive interactions with the LuTHy-LuC. Together, these results suggest that scoring PPIs using a maSVM algorithm can provide two advantages over fixed cutoffs: 1) lower variability of recovered interactions; and 2) the probability of a tested protein pair to be a true positive interaction in the corresponding assay is provided.

To evaluate whether this approach would affect sensitivity or specificity, we next applied the maSVM algorithm to the LuTHy data of all hsPRS-v2 and hsRRS-v2 protein pairs. As before, every training resulted in 50 independent models for each assay (**Figure 1B,C**), which were used to predict the classification of the 138 reference set pairs (60 hsPRS-v2 + 78 hsRRS-v2; 1,104 protein pair configurations), making sure that the protein pair configurations used in the training sets were absent from the paired test sets (**Figure 1D,E**). We then performed ROC analyses to compare the sensitivity and specificity when scoring interactions using cBRET or cLuC ratios or when using maSVM model predicted probabilities (**Figure 1F,G**). Importantly, for both assays, the maSVM model-predicted probabilities showed comparable accuracies to the cBRET and cLuC ratios. Finally, we calculated recovery rates for LuTHy-BRET and LuTHy-LuC protein pairs with >50%, >75% and >95% interaction probability, and observed for LuTHy-BRET that specificity increases with increasing probability, while sensitivity decreases. For LuTHy-LuC, only the sensitivity decreases with increasing probability, as none of the hsRRS-v2 pairs scored positive even at 50% probability (**Figure 1H,I**).

Next, we evaluated whether the maSVM algorithm would be broadly applicable to score quantitative interaction data from various assays. Therefore, we applied it to published benchmarking data (Choi et al, 2019) from six quantitative binary PPI assays: GPCA (Cassonnet et al, 2011), KISS (Lievens et al, 2014), MAPPIT (Eyckerman et al, 2001), NanoBiT (Dixon et al, 2015), N2H (Choi et al, 2019) and SIMPL (Yao et al, 2020) (**Figure EV2L**). For each assay, we obtained probability classifiers for all tagging configurations of hsPRS-v2 interactions and hsRRS-v2 protein pairs (**Figure 1J**), and calculated recovery rates with >50%, >75% and >95% interaction probabilities (**Figure 1K**). For all assays, we observed that specificity increases with increasing probabilities while sensitivity decreases. Moreover, this allowed us to distinguish for each assay, protein pairs to be “unlikely” (>50%), “likely” (>75%), or “very likely” (>95%) detected as true-positive interactions. Overall, this analysis suggested that the maSVM learning algorithm should be universally applicable to reproducibly and robustly classify quantitative PPI results with comparable sensitivity and specificity to traditional approaches, while adding additional information on interaction probabilities and improving comparisons between assays.

### Benchmarking AlphaFold against established reference sets of protein pairs

With the emergence of highly accurate protein structure prediction algorithms, we asked how AlphaFold-Multimer (AFM) performs compared to binary PPI assays, when benchmarked against the hsPRS-v2 and hsRRS-v2 protein pairs (**Figure EV3A**). To this end, we used Google Colaboratory hosted ColabFold that provides accelerated protein complex prediction with the limitation that only protein complexes with less than 1,400 amino acids could be predicted (Mirdita et al, 2022). This resulted in the downsizing of the reference sets to 51 positive (hsPRS-AF) and 67 random (hsRRS-AF) reference pairs for which we predicted five models each (**Source Data Figure EV3-EV4**).

Using PDBePISA (Krissinel & Henrick, 2007), we obtained the interaction interface areas (iA) and the solvation free energies (ΔG) for each model that contained a measurable interface (521 out of 590, see methods for detail, **Figure EV3B**). Since it had been shown that the predicted alignment errors (PAE) can be used to rank and assess the confidence of a predicted PPI (Mirdita et al, 2022), we extracted the PAEs from the AFM structures, which we filtered for amino acids with a predicted local distance difference test (pLDDT) of over 50 to exclude disordered regions (Tunyasuvunakool et al, 2021). Next, we used kmeans clustering to group the respective regions into eight inter-chain clusters (**Figure EV3C-D**). However, if fewer than 10 amino acids in the inter-chain region had a pLDDT scores >50, the PAE scores were discarded to ensure sufficient observations for kmeans subgrouping. From the eight clusters, we extracted the inter-subunit amino acid cluster with the lowest average PAE, for which we postulated that it would most likely be the region involved in the interaction, potentially forming the interaction interface (**Figure EV3E**).

To evaluate which of the obtained measures (i.e. PAE, iA, ΔG) can best distinguish between positive and random reference pairs, we performed ROC analyses for each of the five AFM models (**Figure EV3F**). We found that all three measures are suitable to identify true-positive interactions, but that the average PAE of the inter-subunit cluster and the iA better distinguish between true- and false-positive interactions compared to the ΔG values (**Figure EV5F)**. We then applied the maSVM algorithm to train 50 distinct models on the PAE and iA of the reference sets (**Figure EV4A-C**) that we used to classify 51% of the hsPRS-AF pairs but none of the hsRRS-AF pairs as true positive with >95% interaction probability (**Figure EV4C**). Since, AlphaFold’s neural networks were trained on PDB structures, it shows especially high recovery rates for hsPRS-v2 interactions with a known structure including homologous structures (**Supplementary Table 1**) (Meyer et al, 2018), while interactions without an experimentally solved structures were still recovered at similar sensitivities to the results from the binary interaction assays (**Figure 1K, Figure EV4D**). Overall, the LuTHy assay showed the highest overall recovery rate, but it has to be noted that it was the only assay that was tested in all eight tagging configurations.

In summary, our analysis confirms that AFM is a powerful computational tool capable of distinguishing between well-established positive PPIs and random protein pairs with similar accuracy to commonly used binary interaction assays. This suggests that some complex structures of experimentally determined interactions can be predicted using AFM.

### Classifying binary interactions within multiprotein complexes

To further generalize the overall applicability of the maSVM algorithm to more complex PPI datasets, we screened proteins that are part of well-characterized multiprotein complexes where the same protein is tested against a number of interacting and non-interacting proteins using two quantitative assays: LuTHy and mN2H. To this end, we selected three human complexes based on the following criteria: 1) they consist of at least four subunits; 2) at least one 3D structure is available in PDB (Berman et al, 2000); and 3) at least 80% of cloned open reading frames (ORFs) encoding the reported subunits are available in the human ORFeome 8.1 collection (Yang et al, 2011). This resulted in a list of 24 distinct protein complexes (**Supplementary Table 2**), among which three structurally diverse candidates with well-characterized biological functions were selected: 1) the LAMTOR complex, also termed “Ragulator” complex (Araujo et al, 2017), which regulates MAP kinases and mTOR activity and consists of seven subunits (LAMTOR1, LAMTOR2, LAMTOR3, LAMTOR4, LAMTOR5, RRAGA and RRAGC); 2) the MIS12 complex that connects the kinetochore to microtubules (Petrovic et al, 2016), and is made of five subunits (CENPC1, DSN1, MIS12, NSL1 and PMF1); and 3) the BRISC complex, a large deubiquitinating machinery (Rabl et al, 2019) consisting of five proteins (ABRAXAS2, BABAM1, BABAM2, BRCC3 and SHMT2) (**Figure 2A**).

**Figure 2.**
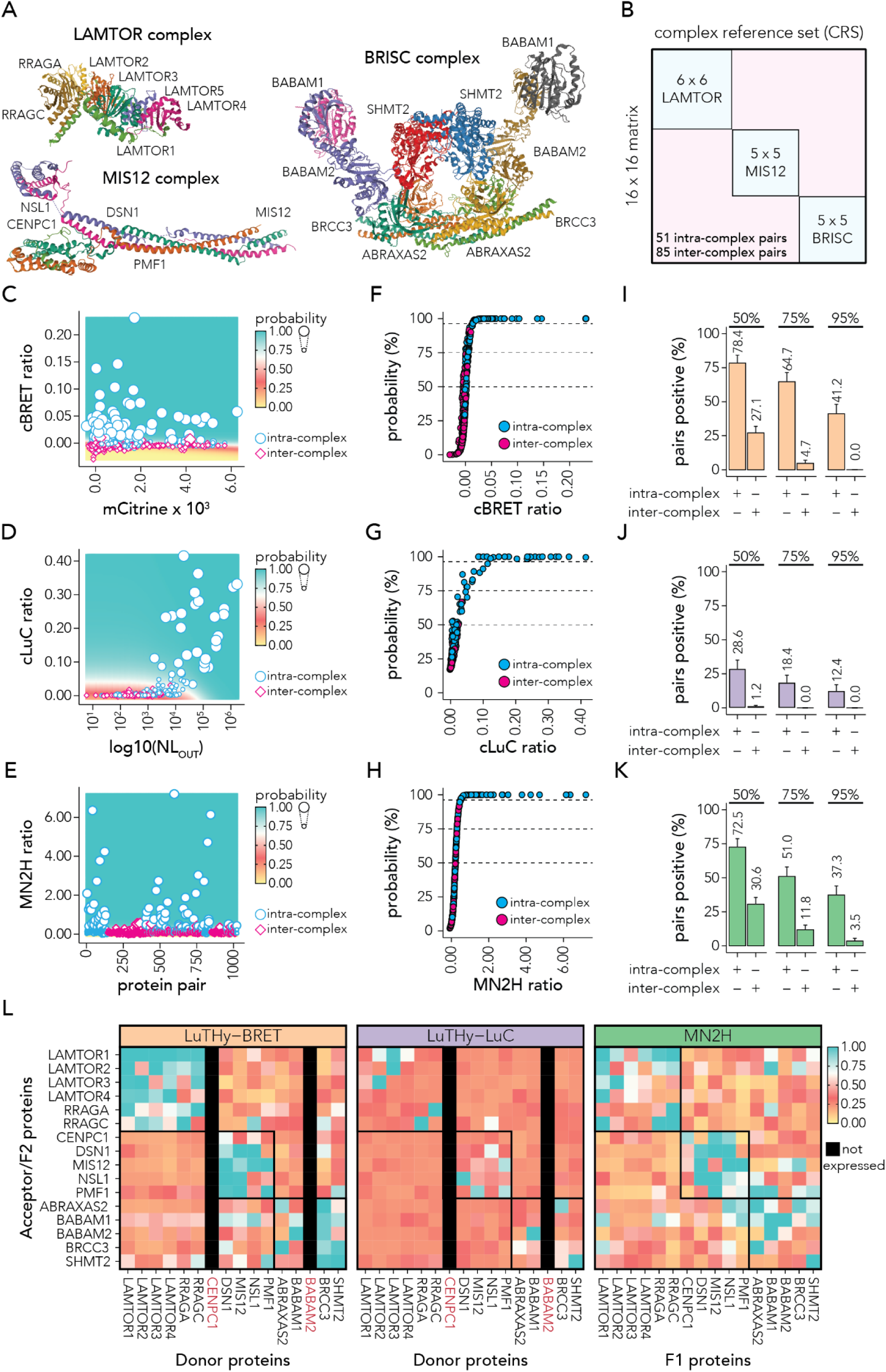
Validating the maSVM algorithm by mapping interactions within multiprotein complexes using the LuTHy and mN2H assays. (**A**) Structures of the protein complexes analyzed in this study: LAMTOR (PDB: 6EHR), MIS12 (PDB: 5LSK), and BRISC (PDB: 6H3C). (**B**) Binary interaction approach to systematically map PPIs within distinct complexes. Every protein subunit from each complex was screened against every other one (all-by-all, 16×16 matrix). (**C-E**) Scatter plot showing (**C**) in-cell mCitrine expression (x-axis) against cBRET ratios (y-axis), (**D**) luminescence after co-precipitation (NL_OUT_) (x-axis) against cLuC ratios (y-axis) or (**E**) the number of protein pairs (x-axis) against the mN2H ratios (y-axis) for all intra-complex (blue) and inter-complex (magenta) protein pairs from all eight tagging configurations. Average classifier probabilities from the 50 maSVM models are displayed as the size of the data points and as a colored grid in the background. (**F-H**) Scatter plot showing on the x-axis the (**F**) cBRET ratios, (**G**) cLuC ratios or (**H**) mN2H ratios against classifier probabilities (y-axis) for all intra-complex (blue) and inter-complex (magenta) protein pairs from all eight tagging configurations. (**I-K**) Bar plots showing the fraction of intra-complex and inter-complex protein pairs that scored above the classifier probabilities of 50%, 75% or 95% by (**I**) LuTHy-BRET, (**J**) LuTHy-LuC and (**K**) mN2H. Only the highest classifier probability per tested tagging configuration is considered. (**L**) Heatmaps showing the classifier probabilities for the Donor/F1 protein pairs (x-axis) against the Acceptor/F2 protein pairs (y-axis) for LuTHy-BRET (orange, left), LuTHy-LuC (purple, middle) and mN2H (green, right). Only the highest classifier probability per tested tagging configuration is shown. Not expressed constructs are filled black and protein names colored in red.

To map interactions between subunits of the LAMTOR, MIS12 and BRISC complexes, out of 17 ORFs encoding the selected target proteins, 16 were sequence verified and cloned into both LuTHy and N2H expression plasmids, whereas the ORF for LAMTOR5 was not available in the human ORFeome 8.1 collection. The resulting search space of 136 unique pairwise combinations, corresponding to a total of 16 subunits for the three complexes, was systematically assessed with LuTHy and mN2H (**Figure 2B, Source Data Figure 2**). Since the different complexes are involved in distinct biological functions, we rationalized that true binary PPIs should only be found between the respective subunits of a given complex (i.e. intra-complex pairs), but not between subunits belonging to different complexes (i.e. inter-complex pairs). Therefore, we treated all inter-complex pairs as negative controls, similar to protein pairs from a random reference set (e.g. hsRRS-v2). We observed that each individual LuTHy and mN2H fusion construct showed a broad distribution of interaction scores for intra- and inter-complex pairs, with a high variability between individual constructs (**Figure EV5A-H**). To compensate for different background signals between constructs in the downstream analysis, we median-normalized outputs from all constructs and performed a robust scaler normalization (Pedregosa et al, 2011) for constructs with a higher interquartile range (IQR) than the IQR of the entire dataset (see methods for details; **Figure EV5A-H**).

We then trained the maSVM algorithm with the normalized LuTHy-BRET, LuTHy-LuC and mN2H data, iteratively sampling from the intra-complex pairs as positive reference interactions and the inter-complex protein pairs as negative reference interactions. Each training resulted in 50 independent models for each assay (**Figure 2C-E**), which were each used to predict the classification of all 136 multiprotein complex pairs (1,024 tagging configurations, cloning for 80 configurations failed), making sure that the protein pair configurations used in the training sets were absent from the paired test sets (**Figure 2F-H**). We calculated recovery rates for LuTHy-BRET, LuTHy-LuC and mN2H for protein pairs with >50%, >75% and >95% interaction probabilities. At >95% interaction probability, we recovered 41.2%, 12.4% and 37.3% of intra-complex interactions by the three different assay versions, and no inter-complex protein pairs were recovered by the LuThy assay and only 3.5% by mN2H (**Figure 2I-L**). Thereby, the fraction of detected intra-complex interactions in the multi-protein complex set is similar to the fraction of recovered hsPRS-v2 interactions for the LuTHy-BRET and mN2H, whereas significantly fewer interactions were recovered by LuTHy-LuC.

Overall, these results demonstrate that in addition to be applicable to classify distinct unrelated heterodimeric interactions like hsPRS-v2 PPIs, the maSVM algorithm can also be used to systematically identify binary PPIs within distinct multiprotein complexes, thanks to outlier-insensitive normalization methods, such as robust-scaler normalization.

### Identifying high-confidence PPI targets for SARS-CoV-2

To apply the maSVM algorithm to an independent test-case scenario, we experimentally assessed all possible pairwise combinations between SARS-CoV-2 proteins using LuTHy (**Figure 3A)**. Since SARS-CoV-2 interactions potentially include homo- and heterodimer, as well as subunit-subunit interactions of larger multiprotein complexes, we trained the maSVM algorithm to generate models representing both the hsPRS-v2/hsRRS-v2 and the multiprotein complex protein pairs (**Figure 3B**). As described above, we median-normalized all constructs of the training set and the SARS-CoV-2 test set and performed a robust scaler normalization for constructs with a higher IQR than the IQR of the entire dataset (see methods; **Figure EV5,6**). Next, we used the normalized training set to train 50 maSVM models for LuTHy-BRET and LuTHy-LuC, in order to predict the classification probabilities of the 350 SARS-CoV-2 protein pairs in the test set (2,548 configurations, **Figure 3C-F**). In total, 26, 56 and 143 protein pairs were classified by the algorithm to interact with >95%, >75% or >50% probability, respectively (**Figure 3G,H, Figure EV7A, Supplementary Table 3**). Among the high-confidence PPIs (>95% interaction probability), we could confirm the structurally resolved interactions between NSP8 and NSP12 (PDB: 6YYT, 7EIZ), NSP10 and NSP16 (PDB: 6WVN, 6W4H) (Rosas-Lemus et al, 2020), NSP10 and NSP14 (PDB: 7DIY, 7EIZ) (Lin et al, 2021; Yan et al, 2021); and the homodimerization of NSP8 (PDB: 7EIZ) (Yan et al, 2021), ORF3a (PDB: 6XDC) (Kern et al, 2021), the nucleocapsid protein (N) (PDB: 6VYO), and the well-established homodimerization of the spike protein (S) (PDB: 6VYB for example) (Walls et al, 2020). We also detected the NSP7 homodimerization, which was previously described in two independent studies (Yin et al, 2020; Wilamowski et al, 2021). Additionally, we confirmed the interactions of NSP3 and N (Jiang et al, 2021), the homodimerization of the envelope protein E (Mandala et al, 2020; Li et al, 2021), and its interaction with the membrane glycoprotein M (Savitt et al, 2021; Yuan et al, 2022). Overall, we did not detect 69 previously reported interactions (Orchard et al, 2014), of which 16 were recently found to interact by Y2H (Kim et al, 2022) (**Supplementary Table 4**). High confidence interactions detected with LuTHy (>95% probability) but not previously reported (Orchard et al, 2014; Kim et al, 2022; Perfetto et al, 2020; Toro et al, 2021), include the heterodimerization of the envelope protein E with NSP6 and ORF7a, as well as between NSP12 and NSP16, NSP2 and NSP3, NSP6 and ORF7a, ORF3b and ORF8, ORF3a and ORF7a and the NSP4 homodimer (**Figure EV7A, Supplementary Table 5**).

**Figure 3.**
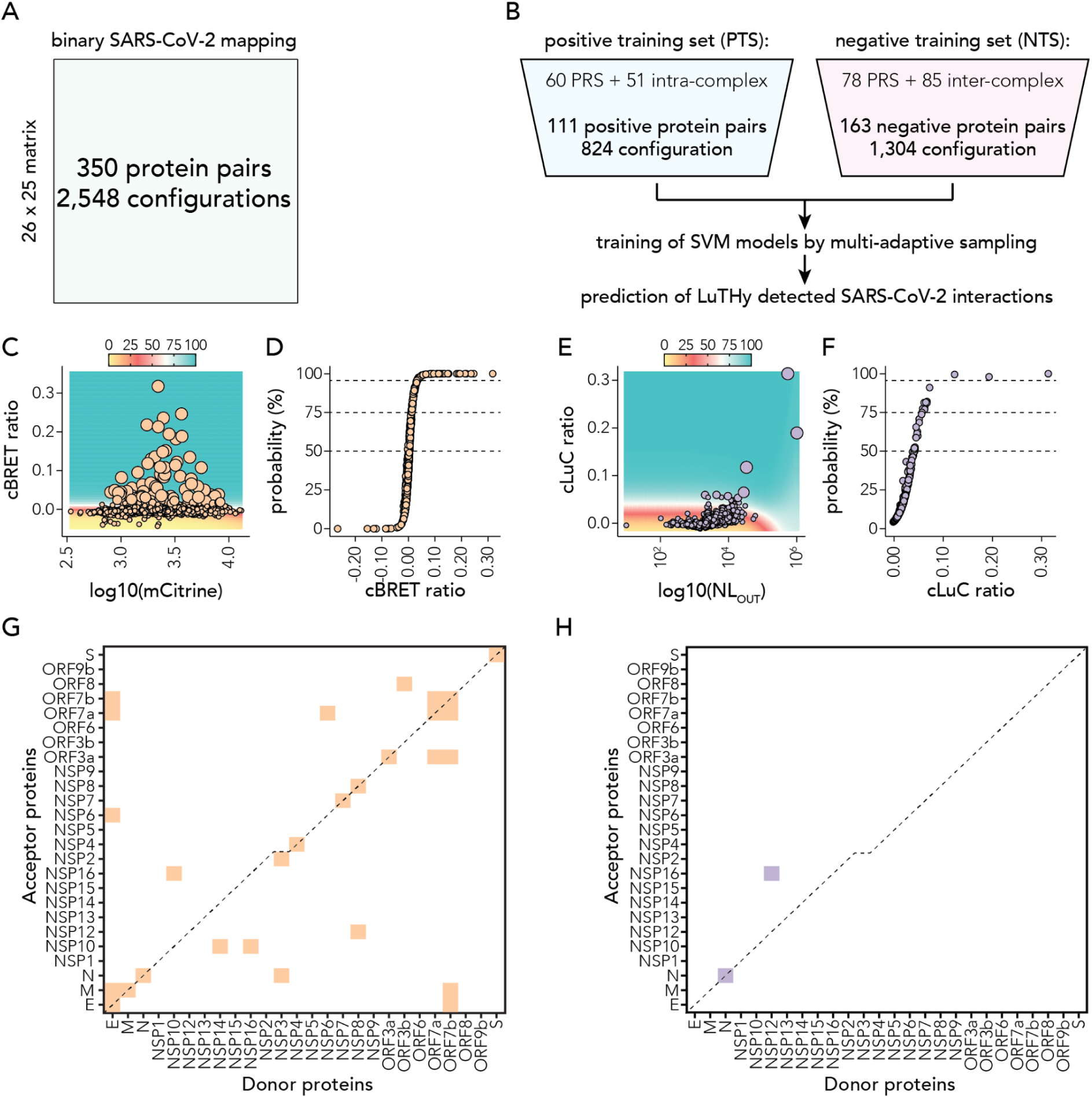
Mapping binary interactions between SARS-CoV-2 proteins. (**A**) Search space between SARS-CoV-2 proteins tested by LuTHy. (**B**) Strategy to classify screened SARS-CoV-2 protein pairs using the maSVM learning algorithm. The positive training set (PTS) was assembled from the hsPRS-v2 and the intra-complex protein pairs of the multiprotein complex set. The negative training set (NTS) was assembled from the hsRRS-v2 and the inter-complex protein pairs of the multiprotein complex set. The trained 50 models were used to predict the classifier probabilities of all LuTHy-BRET and LuTHy-LuC tested SARS-CoV-2 protein pairs. (**C**) Scatter plot showing in-cell mCitrine expression (x-axis) against cBRET ratios (y-axis) for SARS-CoV-2 (orange) protein pairs from all eight tagging configurations. Average classifier probability from the 50 maSVM models is displayed as the size of the data points and as a colored grid in the background. (**D**) Scatter plot showing cBRET ratios (x-axis) against classifier probability (y-axis) for all SARS-CoV-2 (orange) protein pairs from all eight tagging configurations for LuTHy-BRET. (**E**) Scatter plot showing luminescence after co-precipitation (NL_OUT_) (x-axis) against cLuC ratios (y-axis) for SARS-CoV-2 (orange) protein pairs from all eight tagging configurations. Average classifier probability from the 50 maSVM models is displayed as the size of the data points and as a colored grid in the background. (**F**) Scatter plot showing cLuC ratios (x-axis) against classifier probability (y-axis) for all SARS-CoV-2 (orange) protein pairs from all eight tagging configurations for LuTHy-LuC. The number of protein pairs with classifier probabilities of >50%, >75% and >95% are indicated. (**G,H**) Binary heatmaps showing SARS-CoV-2 protein pairs with >95% classifier probability detected with (**G**) LuTHy-BRET and (**H**) LuTHy-LuC. Only the highest classifier probability per tested tagging configuration is shown.

### Predicting the structure of SARS-CoV-2 PPI complexes using AlphaFold-Multimer

To predict dimeric structures of the LuTHy-identified interactions, we used AFM to obtain structures for 23 out of the 26 LuTHy-positive interactions (<1,400 amino acids) and employed PDBePISA to determine the iA and ΔG between each protein pair. Similar to before, we used kmeans clustering to identify the region with the lowest average inter-subunit PAE, which we suggest is most likely participating in the interaction (**Figure EV7B**). We then used the 50 models derived from the maSVM algorithm that were trained on the PAE and iA of the predicted structures of hsPRS-AF and hsRRS-AF pairs, to predict the classification probabilities for the 23 AFM SARS-CoV-2 structures (**Figure EV7C,D**). We thereby identified 15 AFM-predicted structures with a classification probability of over 75% and eight of over 95%. (**Figure EV7E,F**). To strengthen the confidence in the AFM predicted structures, we used the LuTHy-BRET assay to perform donor saturation experiments to determine the in-cell binding strength (BRET_50_) between the interacting proteins, which should correlate to the ΔG of the AFM predicted interface region (**Figure EV8**). As expected, we observed a significant correlation between the BRET_50_ and the ΔG (**Figure EV7G**) similar to published results, where the BRET_50_ was shown to be directly correlated to the binding affinity, K_D_ (Trepte et al, 2018). Furthermore, six out of the eight AFM predicted complex structures with a probability of >95% were also experimentally resolved, such as for example the hetero-dimerization between NSP10 and NSP16, which supports our classification approach (**Figure EV7H**) and demonstrates that AFM can be used to predict structures for binary interactions identified by quantitative binary PPI assays.

### Targeting the NSP10-NSP16 interaction interface by virtual drug screening

After having identified high-confidence SARS-CoV-2 PPIs by LuTHy and predicted their structures using AFM, we next wanted to target one interaction interface for a virtual drug screening. To maximize success rate and prioritize between the predicted structures we applied the following criteria: i) inhibition of one complex member was previously shown to affect viral replication or function, and ii) the 3D complex structure was both experimentally solved and AFM predicted. Based on this rationale, we selected the interaction between NSP10 and the NSP16 RNA methyltransferase (MTase). Importantly, it was reported that inhibiting this interaction completely abrogates the MTase activity of NSP16 (Chen et al, 2011), which is required to ensure normal viral replication (Daffis et al, 2010).

Overall, the five AFM predicted complex structures showed very low predicted aligned errors (**Figure 4A**) and a high overlap to the published 3D structure (Rosas-Lemus et al, 2020) (**Figure 4B**). We used PDBePISA to determine the interaction hot spots (Clackson & Wells, 1995), i.e. the interface residues that contribute most to the binding, and identified lysine 93 of NSP10 having the lowest ΔG (**Figure 4C,D**). We then performed site directed mutagenesis and introduced a charge change at lysine 93 by substituting it with a glutamic acid (Lys93Glu), which resulted in a strong reduction in interaction as measured by LuTHy-BRET (**Figure 4E**). We did not observe an effect on expression levels of either wild-type or Lys93Glu NL-NSP10 nor on mCit-NSP16, suggesting that the overall stabilities of the proteins were not affected by the point mutation (**Figure EV7I,J**). This confirmed that Lys93 is a critical hot spot residue in the NSP10-NSP16 interface, which is consistent with published results (Hamre & Jafri, 2022; Lugari et al, 2010).

**Figure 4.**
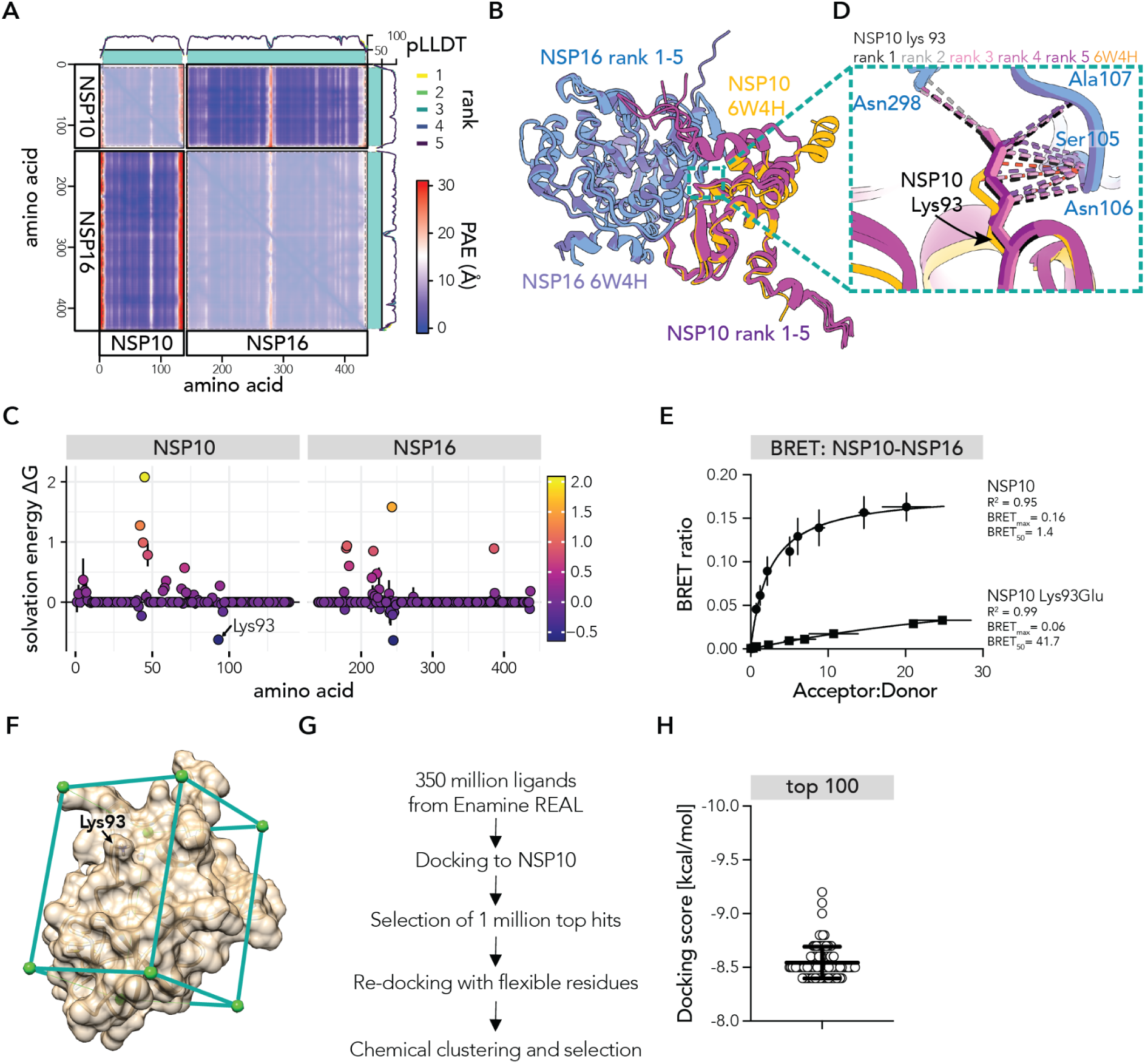
Predicting the NSP10-NSP16 PPI complex with AFM to target the interaction interface by ultra-large virtual drug screening. (**A**) Heatmap showing the predicted alignment error (PAE) of the AlphaFold-Multimer predicted NSP10-NSP16 complex for the rank 1 model. The intra-molecular PAEs are shown with 50% opacity. The predicted local distance difference test (pLDDT) for all five predicted models (rank 1-5) are shown as line graphs on top and on the right of the heatmap. The area with pLDDT scores >50 is highlighted in teal. (**B**) The five models of the AlphaFold-Multimer predicted NSP10-NSP16 complex and the published crystal structure (PDB: 6W4H) are shown. Structures were overlaid using the ‘matchmaker’ tool of ChimeraX. (**C**) Scatter plot showing for each amino acid (x-axis) the solvation free energy (ΔG, y-axis, fill color) upon formation of the interface, in kcal/M, as determined by PDBePISA. Dots represent the average ΔG for the five predicted models and error bars correspond to the standard deviation. Lysine 93 of NSP10 is indicated. (**D**) Zoom-in into the NSP10-NSP16 complex showing the contacts of NSP10s Lysine 93. (**E**) LuTHy-BRET donor saturation assay, where constant amounts of NSP10-NL wt or K93E are co-expressed with increasing amounts of mCitrine-NSP16. Non-linear regression was fitted through the data using the ‘One-Site – Total’ equation of GraphPad Prism. (**F**) Docking box on the NSP10 structure (PDB: 6W4H) used for the ultra-large virtual screen. (**G**) Schematic overview of the workflow of the virtual docking screen using VirtualFlow. (**H**) Docking scores of the top 100 molecules identified that target NSP10.

Based on these results, we decided to target the NSP10-NSP16 interface with small molecules using VirtualFlow, a highly automated and versatile open-source platform for ultra high-throughput virtual drug screening (Gorgulla et al, 2020). We placed the docking box on NSP10 within its interaction interface with NSP16 (**Figure 4F**) and screened ∼350 million compounds from the Enamine REAL library (**Figure 4G**) using Quick Vina 2 (Alhossary et al, 2015). Among the top 100 virtual screening hits, we obtained comparable docking scores as previously described for similarly groove-shaped target regions (Gorgulla et al, 2021), which suggested high quality results (**Figure 4H**). The top ∼10 million (0.03%) hits were re-docked using AutoDock Vina (Trott & Olson, 2010) and Smina Vinardo (Quiroga & Villarreal, 2016), allowing 12 amino acid residues at the binding interface to be flexible. Finally, we selected compounds among the top 10,000 virtual hits that were re-docked with the two different approaches, and subjected ∼2,000 molecules to chemical clustering and filtering (see methods for details). A total of 20 representative molecules were selected, among which 15 were successfully synthesized and used for follow-up studies.

### Inhibiting the NSP10-NSP16 interaction reduces SARS-CoV-2 replication

To prioritize between the 15 selected compounds, we tested their abilities to inhibit the MTase activity of the NSP10-NSP16 complex *in vitro* (Bouvet et al, 2010; Decroly et al, 2011). We incubated the purified NSP10-NSP16 complex with a Cap-0 RNA (m7G, N7-methyl guanosine) and monitored the methylation on the initiating nucleotide, which would generate a Cap-1 structure (**Figure 5A**). Among the 15 selected compounds, three showed a significant reduction in MTase activity compared to the DMSO control (p < 0.05), of which compound 459 had the strongest effect (**Figure 5B**) with about 50% enzyme inhibition and was thus selected for further studies.

**Figure 5.**
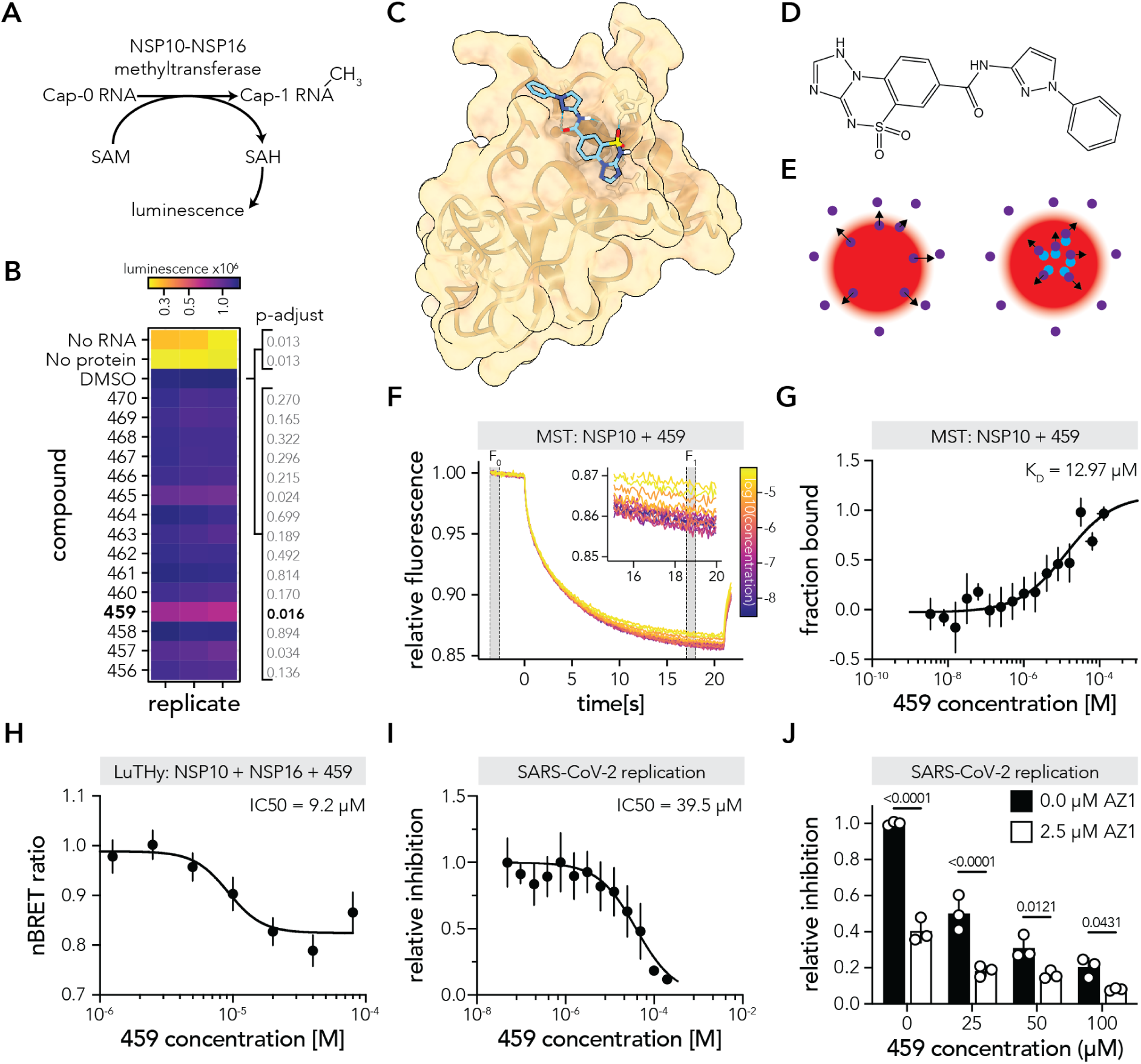
Compound 459 inhibits the NSP10-NSP16 interaction and reduces SARS-CoV-2 replication. (**A**) Schematic overview of the NSP10-NSP16 methyltransferase (MTase) assay. (**B**) Heatmap showing the result of the MTase activity of the NSP10-NSP16 complex in the absence or presence of 100 µM of the top 15 compounds. Statistical significance was calculated with a kruskal-wallis test (p-value = 9.7e-5, chi-squared = 47.656, df = 17, n = 3), followed by a post-hoc Dunn test and adjusted p-values are shown. (**C**) Compound 459 docked onto the NSP10 structure (PDB: 6W4H). (**D**) Chemical structure of compound 459. (**E**) Assay principle of the microscale thermophoresis (MST) assay. The fluorescence intensity change of the labeled molecule (purple) after temperature change induced by an infrared laser (red) is measured. The binding of a non-fluorescent molecule (blue) can influence the movement of the labeled molecule. (**F**) Representative MST traces of labeled NSP10 and different concentrations of unlabeled compound 459. The bound fraction is calculated from the ratio between the fluorescence after heating (F_1_) and before heating (F_0_). (**G**) Scatter plot showing the 459 concentrations (x-axis) against the fraction of 459 bound to NSP10 (y-axis). Non-linear regression was fitted through the data using the ‘One-Site – Total’ equation of GraphPad Prism (n = 3). (**H**) Scatter plot showing the 459 concentrations (x-axis) against the normalized BRET ratio (nBRET ratio) for the interaction between NSP10-NL and mCit-NSP16. Non-linear regression was fitted through the data using the ‘log(inhibitor) vs. response – Variable slope (four parameters)’ equation of GraphPad Prism (n = 4). (**I**) Scatter plot showing the 459 concentrations (x-axis) against the relative luminescence measured from icSARS-CoV-2-nanoluciferase in HEK293-ACE2 cells. Non-linear regression was fitted through the data using the ‘log(inhibitor) vs. normalized response’ equation of GraphPad Prism (n = 9). (**J**) Bar plot showing the relative luminescence measured from icSARS-CoV-2-nanoluciferase in HEK293-ACE2 cells upon incubation with 0, 25, 50 or 100 µM of compound 459 together with 2.5 µM AZ1 or without AZ1 (0.0 µM). Statistical significance was calculated in GraphPad Prism by a ‘Two-way Anova’, where each cell mean was compared to the other cell mean in that row using ‘Bonferroni’s multiple comparisons test’ (n = 3; source of variation: 57.91% 459 concentration, p<0.0001; 28.33% AZ1 concentration, p<0.0001; 11.40% 459/AZ1 interaction, p<0.0001).

To confirm the binding of the virtually docked compound 459 to the NSP10 protein (**Figure 5C,D)**, we applied a microscale thermophoresis (MST) assay, which monitors the temperature-induced movement of fluorescently labeled molecules (Seidel et al, 2012) (**Figure 5E**). To that end, we fluorescently labeled purified NSP10 protein and monitored its movement upon non-fluorescent compound addition. From the MST traces (**Figure 5F**) we calculated the fraction of bound compound 459, which allowed us to determine a binding affinity of ∼12.97 µM (**Figure 5G**). To confirm that compound 459 could disrupt the NSP10-NSP16 interaction, we tested its effect in cells using the LuTHy-BRET assay. When incubating cells that express NL-NSP10 and mCit-NSP16 with compound 459, we observed a modest but significant concentration-dependent reduction in the BRET ratio with a half-maximal inhibitory concentration (IC_50_) of 9.2 μM. This result indicated that the compound inhibits the binding of the two proteins in live cells (**Figure 5H**).

Since it was previously shown that normal MTase activity is required to ensure proper viral proliferation (Daffis et al, 2010), we evaluated the effect of compound 459 on SARS-CoV-2 replication using an infectious cDNA clone-derived reporter assay (Hou et al, 2020; Kim et al, 2022). We observed a concentration dependent decrease of the luminescence signal in the SARS-CoV-2 replication assay, indicating an inhibition of viral replication with an IC_50_ of 39.5 µM (**Figure 5I**).

Finally, to investigate if NSP10-NSP16 inhibition by compound 459 would confer additive effects upon combination with AZ1, an established inhibitor of the human ubiquitin-specific peptidase 25 (USP25) reported to impair SARS-CoV-2 replication (Kim et al, 2022), we tested both compounds in the reporter assay. We assessed SARS-CoV-2 viral replication upon treatment with 2.5 µM AZ1 and increasing concentrations of compound 459, and observed an additive, concentration dependent effect of the two molecules (**Figure 5J**). These results indicate that 459 and AZ1 affect viral replication through distinct mechanisms, and that combinatorial therapies involving distinct drug targets could potentially improve the efficacy of treatments using small molecule drugs.

## DISCUSSION

Targeting PPIs offers great opportunities to tackle various diseases, but it remains a great challenge to reliably identify and modulate protein complexes. To improve comparisons between binary PPI datasets generated in different experiments and laboratories and confidently prioritize potential targets for PPI drug discovery, we have utilized a maSVM learning algorithm (Chang & Lin, 2011; Yang et al, 2017) to coherently score interactions of quantitative PPI datasets. Traditional approaches involve i) selecting a cutoff of maximal specificity, i.e. at which none of the random pairs used as negative controls are scored positive (Choi et al, 2019), ii) ROC analyses with selected false-positive rates of ∼5% (Yao et al, 2020), ∼3% (Trepte et al, 2015), or ∼1-2% (Trepte et al, 2018; Cassonnet et al, 2011), and iii) Gaussian distributions (Taipale et al, 2012). With the maSVM approach, we show for the first time that the automatic classification of large quantitative interaction datasets can be performed with high confidence. Notably, such a classification strategy has been successfully used to predict kinase substrates from phosphoproteomics data (Yang et al, 2019; Kim et al, 2021), or to classify cell types from single cell RNA-sequencing (Abdelaal et al, 2019). We show that the maSVM algorithm can reduce the variability between recovered interactions in different experiments, and provide probabilities for being a true interaction for every protein pair tested. For example, we observed that in the LuTHy-BRET and mN2H assays, one pair from the random reference set hsRRS-v2 (SLC6A1+TM4SF4) was classified as a true-positive interaction with >99% probability (**Figure 1J**). Due to its definition as a negative or random interaction, a traditional scoring approach would have classified this pair as strictly negative and thus classify similar or lower scoring pairs also as negative. However, it is very likely that in the LuTHy-BRET and mN2H assays, these two proteins interact biophysically when overexpressed in HEK293 cells, making the pair a potential pseudo-interaction, i.e. a true biophysical interaction without *in vivo* biological relevance (Braun et al, 2009). The maSVM algorithm is able to deal with such exceptions and thus increases robustness in scoring PPIs even when inconsistencies among assays or in reference sets are present. We also show that the maSVM algorithm is universally applicable to classify binary PPIs from different quantitative datasets, including the interface analysis of AFM-predicted complex structures. Thus, it provides a framework to directly compare the results of various binary interaction assays and *in silico* predictions, which will lead to increased reproducibility and interpretability of results between experiments and methods.

We have further expanded the use of the unbiased maSVM algorithm scoring approach to map binary PPIs in three established multiprotein complexes, and to identify potential targets for drug discovery among SARS-CoV-2 proteins. Interestingly, due to the high amino acid conservation of interaction interfaces (Guharoy & Chakrabarti, 2010; Gupta et al, 2020), PPI-targeting drugs could provide unique advantages over other types of drugs such as vaccines or antivirals circumventing mutations in the pathogens’ genomes that can result in immune evasive properties. Of the 26 detected high-confidence (≥95% probability) SARS-Cov-2 PPIs we detected here, 18 are known interactions according to the IMEx database (Orchard et al, 2014), while eight were not reported before (**Supplementary Table 5**). Further characterizing these previously undescribed interactions and understanding their biological functions could result in the identification of novel drug targets for SARS-CoV-2. In particular, our approach helped in prioritizing the NSP10-NSP16 interaction, where NSP10 serves as a cofactor for NSP16s’ MTase activity (Decroly et al, 2011), an enzyme crucial to single-stranded RNA viruses (Ramdhan & Li, 2022). This enzymatic complex has been a target of previous drug screening campaigns (Nencka et al, 2022) and peptides inhibiting the interaction could be successfully identified (Wang et al, 2015; Ke et al, 2012). Even though the experimental structure of the SARS-CoV-2 methyltransferase complex was becoming available during the course of this study (Rosas-Lemus et al, 2020) and was already described for SARS-CoV-1 (Chen et al, 2011), we demonstrate that our pipeline of AI-guided experimental PPI mapping, structure prediction and experimental validation is able to define such protein complexes with high-confidence and to identify hot spots on their interaction interfaces. We used this information to apply a virtual PPI inhibitor screening strategy with VirtualFlow, an *in silico* method which allows to virtually screen billions of compounds and assess their different binding poses on the targeted surface area. VirtualFlow was already used to target 17 SARS-CoV-2 proteins, including the NSP10-NSP16 interface (Gorgulla et al, 2021). Here, we targeted the same interaction interface by virtual screening but further experimentally validated hit compounds for enzymatic inhibition of the NSP10-NSP16 protein complex and binding of hit compound 459 to the NSP10 target site. We also show that this compound inhibits the NSP10-NSP16 interaction and prevents SARS-CoV-2 replication with additive effects when combined with AZ1, a human USP25 inhibitor disrupting SARS-CoV-2 replication (Kim et al, 2022). Interestingly, the predicted target binding site of compound 459 (**Figure 5C**) is highly conserved among coronavirus groups (Lugari et al, 2010) and thus could potentially also inhibit the replication of other viruses belonging to the Coronaviridae family. However, despite its effects, compound 459 is an experimental compound with micromolar affinity that will require structure-activity relationship studies to further improve its chemical scaffold and associated affinity and efficacy for further investigations.

A prerequisite to enable such virtual drug and PPI inhibitor screening approaches is the availability of high-resolution protein and protein complex structures. Protein and protein complex structure prediction has exploded in the last few years with the development of AlphaFold and RoseTTAFold that allow structure prediction of entire proteomes (Jumper et al, 2021; Baek et al, 2021), as well as the prediction of protein-complexes and protein-protein interactions (Humphreys et al, 2021; Burke et al, 2021; Evans et al, 2022; Gao et al, 2022; Bryant et al, 2022). Through ColabFold, which combines such algorithms with the fast homology search MMseqs2, the immense computing power needed was reduced and is now available within the Google Colaboratory, which makes protein structure prediction accessible to all (Mirdita et al, 2022). As current approaches have already predicted tens of thousands of interactions (Burke et al, 2021; Humphreys et al, 2021), it seems feasible to predict complex structures of the entire theoretical SARS-CoV-2 binary interactome, i.e. all 26 proteins against each other for a total of 650 pairwise combinations. Overall, combining such *in silico* and wet lab techniques for both the identification and validation of PPIs as well as for drug screening and validation of drug effects, should help to speed up the process of developing PPI modulating therapeutics.

## AUTHOR CONTRIBUTIONS

Conceptualization of benchmarking approach: P.T., S.G.C., J.O., M.V., E.E.W.; Conceptualization of machine learning, AFM, SARS-CoV-2 PPI screening: P.T., C.S.; Conceptualization of SARS-CoV-2 VirtualFlow: C.S.; Methodology: P.T., C.S., S.K., S.B.M, S.G.C., J.B., I.M., T.H., J.O.; Software: P.T.; Software VitualFlow: C.S.; Software LuTHy-Python: M.S.; Formal analysis: P.T., C.S.; Investigation: P.T., C.S., S.K., S.B.M, S.G.C., J.B, I.M., P.C., S.G., M.Z., S.B., Ma.Li., N.S., A.S., K.S., T.H., J.O.; Resources: A.S., M.L., J.C.T., M.V., E.E.W.; Data Curation: P.T., C.S., S.G.C., E.S.R., Mi.Li., Y.W., T.H., J.O.; Writing - Original Draft: P.T., C.S., J.O.; Writing - Review & Editing: P.T., S.G.C., A.S., J.O., M.A.C., M.V., E.E.W.; Visualization: P.T., C.S., Supervision: P.T., C.S., S.G.C, M.A.C., D.E.H., M.L., J.O., J.C.T., M.V., E.E.W., Project administration: P.T., C.S., M.L., J.O., J.C.T., M.V., E.E.W.; Funding acquisition: P.T., S.G.C., M.A.C., D.E.H., M.L., J.O., J.C.T., M.V., E.E.W.

## ACKNOWLEDGEMENTS

The authors would like to thank all members of the Wanker, Vidal, Twizere and Landthaler laboratories for helpful discussions throughout this project. We would also like to thank Luke Lambourne for critically reading the manuscript and providing feedback on the machine learning and AlphaFold predictions. We also thank the team of the Protein Production & Characterization Technology Platform of the Max Delbrück Center (MDC), Berlin, Germany, for excellent technical assistance. Additional gratitude to Sebastian Lührs (JURECA supercomputer, Forschungszentrum Jülich) and Martin Siegert (MaxCluster, MDC) for technical support on cluster usage. This work was supported by the Helmholtz Association, iMed and Helmholtz-Israel Initiative on Personalized Medicine (Germany); the Federal Ministry of Education and Research and e:med Systems Medicine – IntegraMent 01GS0844 (Germany) all to E.E.W.; as well as by the CHDI Foundation (USA); the German Cancer Consortium DKTK (Germany) and the Deutsche Krebshilfe, ENABLE (Germany) to E.E.W. and P.T. I.M. and M.L. were in part funded by DFG LA 2941/17-1. This work was also supported by a Claudia Adams Barr Award to S.G.C., Fonds de la Recherche Scientifique (F.R.S.-FNRS) Grants FC27371 to J.O., FC38907 to J.B., FC34947 to S.B.M. and PER-40003579 to J.C.T. and a Wallonia-Brussels International (WBI)-World Excellence Fellowship to J.O. Additional support was provided by NIH grants P50HG004233 and R01GM130885 awarded to M.V. and U41HG001715 awarded to M.V., D.E.H. and M.A.C. This work was also supported by the LabEx IBEID (grant 10-LABX-0062). J.O. thanks the Rega Institute for Medical Research for financial support. M.V. is a Chercheur Qualifié Honoraire, and J.C.T. a Maître de Recherche of the Fonds de la Recherche Scientifique (F.R.S.-FNRS, Wallonia-Brussels Federation, Belgium).

## DATA AVAILABILITY

The protein interactions from this publication have been submitted to the IMEx (http://www.imexconsortium.org) consortium through IntAct (Orchard et al, 2014) and assigned the identifier IM-29174. All UniProt and RCSB-PDB accession codes, as well as the AlphaFold-Multimer predictions are provided in the Source Data. Python and R codes are available upon request.

## MATERIALS AND METHODS

### ORF sequencing and plasmid generation

For hsPRS-v2 and hsRRS-v2 proteins, the corresponding sequence-verified entry vectors published in Choi et al(Choi et al, 2019) were Gateway cloned into the different LuTHy destination plasmids. ORFs for subunits of the LAMTOR, MIS12 and BRISC complexes were taken from the CCSB human ORFeome 8.1, which is a sequence confirmed clonal collection of human ORFs in a Gateway entry vector system(Yang et al, 2011). In total, 16 entry plasmids were picked from the collection, single clones were isolated, and ORFs were PCR-amplified and confirmed by bi-directional Sanger DNA sequencing. Entry clones were shuttled into LuTHy (Addgene #113446, #113447, #113448, #113449) and N2H (Addgene #125547, #125548, #125549, #125559) destination vectors using the Gateway Cloning Technology. SARS-CoV-2 ORF cDNA library was obtained from Kim et al. (Kim et al, 2020) via Addgene. NSP10, NSP14, NSP16 and NSP10 mutant cDNA entry clones were generated by gene synthesis and subcloning into pDONR221 (GeneArt, Thermo Fisher Scientific). All cDNA clones were sequence verified and shuttled into LuTHy destination plasmids. All resulting vectors were analyzed by PCR-amplification of cloned ORFs and DNA gel electrophoresis (N2H plasmids), or restriction digestion and sequence validation (LuTHy plasmids). For the LuTHy assay, additional control plasmids (PA-NL, Addgene #113445; PA-mCit-NL, Addgene #113444; PA-mCit, Addgene #113443; NL, Addgene #113442) were used, as previously described (Trepte et al, 2018). For the mN2H assay, additional control plasmids (pDEST-N2H-F1-empty vector, Addgene #125551; pDEST-N2H-F2-empty vector, Addgene #125552) were used, as previously described (Choi et al, 2019).

### LuTHy assay procedure

The LuTHy assay was performed as previously described (Trepte et al, 2018). In brief, HEK293 cells were reversely transfected in white 96-well microtiter plates (Greiner, #655983) at a density of 4.0-4.5×10^4^ cells per well with plasmids encoding donor and acceptor proteins. After incubation for 48 h, mCitrine fluorescence was measured in intact cells (mCit_cell_, Ex/Em: 500 nm/530 nm). For LuTHy-BRET assays, coelenterazine-h (pjk, #102182) was added to a final concentration of 5 μM (5 mM stock dissolved in methanol). Next, cells were incubated for an additional 15 min and total luminescence as well as luminescences at short (370-480 nm) and long (520-570 nm) wavelengths were measured using the Infinite^®^ microplate readers M200, M1000, or M1000 PRO (Tecan). After luminescence measurements, the luminescence-based co-precipitation (LuC) assay was performed. Cells were lysed in 50-100 μL HEPES-phospho-lysis buffer (50 mM HEPES, 150 mM NaCl, 10% glycerol, 1% NP-40, 0.5% deoxycholate, 20 mM NaF, 1.5 mM MgCl_2_, 1 mM EDTA, 1 mM DTT, 1 U Benzonase, protease inhibitor cocktail (Roche, EDTA-free), 1 mM PMSF, 25 mM glycerol-2-phosphate, 1 mM sodium orthovanadate, 2 mM sodium pyrophosphate) for 30 min at 4°C. Lysates (7.5 µL) were transferred into small volume 384-well microtiter plates (Greiner, #784074) and fluorescence (mCit_IN_) was measured as previously described. To measure the total luminescence (NL_IN_), 7.5 µL of 20 µM coelenterazine-h in PBS was added to each well and the plates incubated for 15 more minutes. For LuC, small volume 384-well microtiter plates (Greiner, #784074) were coated with sheep gamma globulin (Jackson ImmunoResearch, #013-000-002) in carbonate buffer (70 mM NaHCO_3_, 30 mM Na_2_CO_3_, pH 9.6) for 3 h at room temperature, and blocked with 1% BSA in carbonate buffer before being incubated overnight at 4°C with rabbit anti-sheep IgGs in carbonate buffer (Jackson ImmunoResearch, #313-005-003). 15 µL of cell lysate was incubated for 3 h at 4°C in the IgG-coated 384-well plates. Then, all wells were washed three times with lysis buffer and mCitrine fluorescence (mCit_OUT_) was measured as described. Finally, 15 µL of PBS buffer containing 10 μM coelenterazine-h was added to each well and luminescence (NL_OUT_) was measured after a 15 min incubation period.

### LuTHy data analysis

Data analysis was performed as previously described (Trepte et al, 2018). In brief, the LuTHy-BRET and LuTHy-LuC ratios from BRET and co-precipitation measurements are calculated as follows:

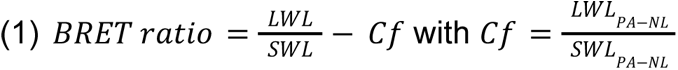

with LWL and SWL being the detected luminescences at long (520–570 nm) and short (370–480 nm) wavelengths, respectively. The correction factor (Cf) represents the donor bleed-through value from the PA-NL only construct. The corrected BRET (cBRET) ratio is calculated by subtracting the maximum BRET ratios of control 1 (NL/PA-mCit-Y), or of control 2 (NL-X/PA-mCit) from the BRET ratio of the studied interaction (NL-X/PA-mCit-Y). For the LuC readout, the obtained luminescence precipitation ratio (PIR) of the control protein PA-NL (PIR_PA-NL_) is used for data normalization, and is calculated as follows:

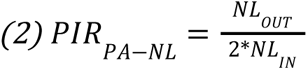

with NL_OUT_ being the total luminescence measured after co-IP and NL_IN_ the luminescence measured in the cell extracts, directly after lysis. Subsequently, LuC ratios are calculated for all interactions of interest, and normalized to the PIR_PA-NL_ ratio:

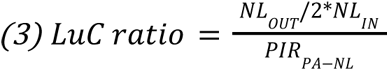

Finally, a corrected LuC (cLuC) ratio is calculated by subtracting either the LuC ratio of control 1 (NL/PA-mCit-Y), or of control 2 (NL-X/PA-mCit) from the LuC ratio of the studied interaction (NL-X/PA-mCit-Y). The calculated LuC ratios obtained for controls 1 and 2 are then compared to each other, and the highest value is used to correct the LuC ratio of the respective interaction. The described analysis was semi-automated, by using a Python script that copied the raw data from the Excel files generated by the Tecan plate readers into Excel templates that were manually controlled for missing values and outliers. Following, all Excel files were imported into R to calculate cBRET and cLuC ratios as described above.

### Mammalian cell-based version of the N2H assay (mN2H)

HEK293T cells were seeded at 6×10^4^ cells per well in 96-well, flat-bottom, cell culture microplates (Greiner Bio-One, #655083), and cultured in Dulbecco’s modified Eagle’s medium (DMEM) supplemented with 10% fetal calf serum at 37 °C and 5% CO_2_. 24 h later, cells were transfected with 100 ng of each N2H plasmid (pDEST-N2H-N1, -N2, -C1 or -C2) using linear polyethylenimine (PEI) to co-express proteins fused with complementary NanoLuc fragments, F1 and F2. The stock solution of PEI HCl (PEI MAX 40000; Polysciences Inc; Cat# 24765) was prepared according to the manufacturer’s instructions. Briefly, 200 mg of PEI HCl powder were added to 170 mL of water, stirred until complete dissolution, and pH was adjusted to 7 with 1 M NaOH. Water was added to obtain a final concentration of 1 mg/mL, and the stock solution was filtered through a 0.22 µm membrane. The DNA/PEI ratio used for transfection was 1:3 (mass:mass). 24 h after transfection, the culture medium was removed and 50 µL of 100x diluted NanoLuc substrate (Furimazine, Promega Nano-Glo, N1120) was added to each well of a 96-well microplate containing the transfected cells. Plates were incubated for 3 min at room temperature. Luciferase enzymatic activity was measured using a TriStar or CentroXS luminometer (Berthold; 2 s integration time).

### LuTHy-BRET donor saturation assays

LuTHy donor saturation assays were performed as previously described (Trepte et al, 2018). In brief, increasing acceptor expression plasmids were transfected to a constant amount of donor plasmids as described above (see “LuTHy procedure”). After 48 h, coelenterazine-h (pjk, #102182) was added to a final concentration of 5 μM. Cells were incubated for an additional 15 min and in-cell mCitrine and luminescence signals were quantified. Infinite® microplate readers M1000 or M1000Pro (Tecan) were used for the readouts with the following settings: fluorescence of mCitrine recorded at Ex 500 nm/Em 530 nm, luminescence measured using blue (370–480 nm) and green (520–570 nm) band pass filters with 1,000 ms (LuTHy-BRET). For data analysis, BRET ratios were calculated as described above. Acceptor to donor ratios were estimated by calculating the ratio of the fluorescence intensity of the acceptor to the total luminescence of the donor and normalization to the acceptor to donor signal intensities of the PA-mCit-NL tandem construct (4).

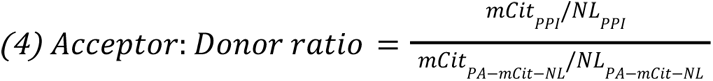

### Processing publicly available data and selecting multiprotein complexes

Reference PDB structures and homologous structures for interactions in the hsPRS-v2 were obtained from interactome insider http://interactomeinsider.yulab.org/ (Meyer et al, 2018). Publicly available binary protein interaction datasets used in this study came from the original Choi et al (Choi et al, 2019) and Yao et al (Yao et al, 2020) publications. Human protein complexes used in this study were selected based on the following criteria. First, human protein complexes should have at least one experimentally determined structure in PDB (Berman et al, 2000). Second, the complex should have at least four subunits. Third, at least 80% of entry clones for individual subunits of a complex should be present in the human ORFeome 8.1 collection (Yang et al, 2011). A total of 24 distinct complexes (**Supplementary Table 2**) with different PDB structures met those criteria, and three protein complexes with well-documented biological functions were selected from this list: LAMTOR, BRISC and MIS12.

Published SARS-CoV-2 interactions were extracted from the IMEx database using the search term ‘coronavirus’ (https://www.ebi.ac.uk/intact/imex) as of 2023-01-20 (Orchard et al, 2014).

### Multi-adaptive support vector machine learning algorithm

The multi adaptive supporting vector machine learning algorithm was adapted from Yang et al (Yang et al, 2019) and Kim et al (Kim et al, 2021). Standardization of datasets is a common requirement for many machine learning algorithms. We observed a strong construct specific variance in the multiprotein complex reference set (**Figure EV5**) and the binary SARS-CoV-2 mapping (**Figure EV6**). We argued that constructs with a high variance are unlikely to form significantly more or less interactions than constructs with a low variance, but rather assumed that the observed variance is probably a technical artifact that could for example be explained by “sticky” proteins. To be able to apply a machine learning algorithm that universally applies to all constructs, we used a percentile-based scaling approach (RobustScaler, https://scikit-learn.org/), that is not influenced by a small number of very large marginal outliers, i.e. not influenced by high scoring, e.g. true-positive interactions. Initially, all interaction scores per construct were median centered and the interquartile range (IQR) between the 25th to 75th quartile defined as the reference IQR (IQR_ref_) of all constructs. Next, interaction scores from constructs with a higher IQR than the IQR_ref_ were scaled to the IQR_ref_.

Machine learning with adaptive sampling (AdaSampling) is a framework developed for both positive-unlabeled (PU) learning and learning with class label noise (LN) (Yang et al, 2019). For binary interaction assays, class LN can refer to protein pairs in the PRS that score negative or protein pairs in the RRS that score positive. Through adaptive sampling, the class mislabeling probability can be estimated, which progressively reduces the risk to select mislabeled instances for model training (Yang et al, 2019). Reference sets were assembled as described in the results section (**Figure 1A**, Step 1), and the interaction scores normalized as described above, using the IQR_ref_ of the combined training and test sets (**Figure 1A**, Step 2). Next, for the LuTHy-BRET assay, we selected the cBRET ratio and mCit_cell_ as training features since the BRET ratio is directly influenced by acceptor expression (mCit). For the LuTHy-LuC we selected the cLuC ratio and NL_OUT_ as training features, since the NL_OUT_ luminescence provides direct information on the total amount of protein precipitated that is not influenced by expression values. As training features for the other assays, we used for the SIMPL assay: ratio.mean, SIMPL.mean and for the vN2H, yN2H, mN2H, GPCA, KISS, MAPPIT and NanoBiT assays each respective interaction score (**Figure 1A**, Step 2). For AlphaFold-Multimer the training features were the “Predicted Aligned Error” (PAE) and the interface area (iA), the latter being derived from the AFM predicted structures using PDBePISA.

Positive training sets were assembled from the reference set by weighted sampling j protein pairs (minimum 30) including only interactions above the median of all interactions scores. Negative training sets were assembled by non-weighted sampling the same number of protein pairs as in the positive training sets. Next, a support vector machine learning algorithm was trained using the ‘svm’ function of the ‘e1071’ package (version 1.7-11) for R and the following parameters: type = “C-classification”; kernel = “linear”; cost = 100.

The resulting SVM model was used to re-classify the training set in 5 iterations, which was used to predict the classifier probabilities of the protein pairs in the test set, from which protein pairs in the training set were excluded. Training and predictions were repeated 50 times in a paired fashion, whereas if no classifier probabilities for protein pairs after 50 cycles were obtained, these protein pairs were excluded from the training set and the training repeated an additional 10 times. Finally, the average classifier probabilities from all models were calculated.

### AlphaFold-Multimer protein complex prediction

Protein complex structures were predicted by AlphaFold-Multimer (Evans et al, 2022) using ColabFold (Mirdita et al, 2022) with the following parameters: use_amber: ‘no’; template_mode: ‘none’ (no pdb template information is used); msa_mode: ‘MMseq2 (UniRef+Environmental)’, pair_mode: ‘unpaired+paired’ (pair sequences from same species); model_type: ‘auto’ (AlphaFold2-multimer-v2); num_models: 5; num_recycles: 3; rank_by: ‘auto’; stop_at_score: 100. For each protein complex, 5 models were predicted, and the resulting pLDDT and PAE scores extracted from .json files using the ‘fromJSON’ function of the ‘rjson’ library for R. For each model, inter-subunit PAE values with pLLDT scores >50 and containing at least 10 amino acids, were k-means clustered row-wise and column-wise into each 2 clusters (4 total), using the ‘kmeans’ function of the ‘stats’ package for R. For 4 models of hsPRS-v2 PPIs and 6 models of hsRRS-v2 protein pairs the PAE clustering failed. For all others, the average PAE was calculated for each cluster in order to identify the interaction interface region with the closest average distance.

PDBePISA was used to determine the interaction interface and solvation free energy of the AlphaFold-Multimer predicted structures (Krissinel & Henrick, 2007) using the python script PisaPy (https://github.com/hocinebib/PisaPy) for batch analysis. The PisaPy output interfacetable.xml files were processed in R using the ‘read_xml’ function of the ‘xml2 package’. PDBePISA prediction failed for all five models of the hsRRS-v2 protein pair PSMD12+CRIPT, and for 13 hsPRS-v2 interactions and 56 hsRRS-v2 proteins at least one, but not all five models were predicted (see **Figure EV3B**). Overall, 4.75±0.8 models were successfully predicted for all hsPRS-v2 interactions and 4.23±1.4 models for hsRRS-v2 protein pairs.

### Ultra-large virtual screening with VirtualFlow

Ultra-large virtual screening was performed using the VirtualFlow workflow engine (Gorgulla et al, 2020) on a Sun Grid Engine (MaxCluster, Max Delbrück Center) or a SLURM-managed (JURECA supercomputer, Forschungszentrum Jülich) high performance computing cluster (Krause & Thörnig, 2018). A subset (∼350M ligands) of the “ready-to-dock” Enamine REAL library was docked onto the experimentally validated NSP10 interaction interface. For primary ultra-large docking, Quick Vina 2 (Alhossary et al, 2015) was used with exhaustiveness set to 1. Ligands were ranked based on their predicted binding free energies in kcal/mol. Then, the top 10M scoring ligands were re-docked using Smina Vinardo (Apr 2, 2016, based on Autodock Vina version 1.1.2) and Autodock Vina (version 1.1.2) (Trott & Olson, 2010) with flexible residues at the target region (Val, Met, Phe, Ser, Cys, Cys, Arg, His, Tyr, Lys, Lys, His). AutoDockTools was used to generate the rigid and flexible structures in the PDBQT format. Exhaustiveness in the re-docking was also set to 1 and two iterations for each docking scenario were conducted. The size of the cuboid docking box for all scenarios and docking runs was set to 75.647 x 16.822 x 17.631 Å. Ligands were ranked by the mean scores of the replica of the Smina Vinardo and Autodock Vina dockings, respectively. Finally, the ligands, which were present in the top 10K of both docking scenarios were selected (∼2K ligands). The ligands were then chemically clustered to identify cluster representatives. Clustering was performed using the Cluster Molecules component embedded in the Pipeline Pilot Software (BIOVIA Pipeline Pilot, Release 2018, San Diego, Dassault Systèmes) using FCFP_4 Fingerprints and an average of 60 compounds per cluster in order to obtain 30 clusters. Molecules were filtered based on reactivity, toxicity and drug-likeness using Pipeline Pilot Software according to Horvath et al. (Horvath et al, 2014) and solubility was predicted according to Cheng & Merz (Cheng & Merz, 2003). From each cluster, a representative molecule was selected based on reactivity, toxicity and drug-likeness properties as well as their predicted solubility. In total, 20 molecules were selected and ordered for synthesis at Enamine Ltd. (Kiev, Ukraine). From the 20 compounds, 15 (#456 to #470) were successfully synthesized and delivered. Compounds were diluted to 10 or 50 mM stock solutions in DMSO, and stored at −20°C.

### Recombinant protein production

The NSP10 expression construct comprising amino acids 23-145 was cloned into a modified pET28a vector, resulting in the expression of an N-terminal His_6_-tagged protein (MGSDKIHHHHHHNSTVLS…GCSCDQ). The protein was produced at 17°C using *E. coli* BL21-AI cells (Thermo Fisher Scientific), induced with 0.5 mM isopropyl β-D-1-thiogalactopyranoside (IPTG) and 0.2% (v/v) L-arabinose. For purification, cells were resuspended in lysis buffer (50 mM sodium phosphate pH 7.8, 0.5 M NaCl, 5% glycerol) supplemented with 0.25% (w/v) 3-[(3-cholamidopropyl)-dimethylammonio]-1-propane-sulfonate (CHAPS), 1 mM phenylmethyl-sulfonyl fluoride (PMSF), 3000 U/mL lysozyme (Serva) and 7.5 U/mL RNase-free DNase I (AppliChem), lysed by multiple freeze-thaw cycles and the extract was cleared by 1 h centrifugation at 34.000×g. The His_6_-fusion protein was captured from the supernatant using metal affinity chromatography on a HisTrap™ FF Crude Column (Cytiva) equilibrated with 20 mM Tris-HCl pH 8.0 and 0.5 M NaCl. After several wash steps, the protein was eluted with 20 mM Tris-HCl pH 8.0, 0.5 M NaCl, and 250 mM imidazole, and further purified by a size-exclusion chromatography step on a 26/600 Superdex 75 prep grade column (Cytiva) equilibrated with 20 mM HEPES pH 7.5 and 0.2 M NaCl. The purified protein was concentrated to 8 mg/mL, flash-frozen with liquid nitrogen, and stored at −80°C until further use.

The NSP10-NSP16 co-expression plasmid (cloned into pETDuet-1, Novagen) was transformed into *E. coli* BL21-AI cells (Thermo Fisher Scientific). The protein was produced at 17°C upon induction with 0.5 mM IPTG and 0.2% (v/v) L-arabinose. For purification, cells were resuspended in lysis buffer (50 mM Tris-HCl pH 7.6, 0.5 M NaCl, 5% glycerol) supplemented with 0.25% (w/v) CHAPS, 1 mM PMSF, 3000 U/mL lysozyme (Serva), 7.5 U/mL RNase-free DNase I (AppliChem), 1 mM MgCl_2_, lysed by multiple freeze-thaw cycles, and the extract was cleared by centrifugation. The His_6_-tagged NSP16, complexed with co-expressed NSP10, was captured from the supernatant using metal affinity chromatography on a HisTrap™ FF Crude Column (Cytiva) equilibrated with 50 mM Tris-HCl pH 7.6, 0.5 M NaCl, and 5% glycerol. After several wash steps, the protein was eluted with the same buffer including 250 mM imidazole, and further purified by a size-exclusion chromatography step on a 16/600 Superdex 75 prep grade column (Cytiva) equilibrated with PBS pH 7.4 and 0.3 M NaCl. The purified protein was supplemented with 5% (v/v) glycerol and 1 mM DTT.

The purified proteins were flash-frozen with liquid nitrogen and stored at −80°C until further use. The molecular mass of all purified proteins was confirmed by intact mass analyses using an Agilent 1290 Infinity II UHPLC system coupled to an Agilent 6230B time-of-flight (TOF) instrument.

### Methyltransferase assay

Methyltransferase activity of the NSP10-NSP16 complex and inhibitor screening was performed using the MTase-Glo assay (Cat. No. V7601, Promega) (Hsiao et al, 2016) according to the manufacturer’s instructions. In brief, 18.75 µM RNA cap-0 oligo (5’-(N7-MeGppp)ACAUUUGCUUCUGAC-3’) or no RNA as control was incubated in the presence of 740 nM co-purified NSP10-NSP16 protein complexes or no protein as control in reaction buffer (20 mM Tris-HCl pH 8.0, 50 mM NaCl, 1 mM EDTA, 3 mM MgCl2, 0.1 mg/ml BSA, 1 mM DTT) supplemented with 0.2 U/µL SUPERase RNAse inhibitor and 40 µM S-adenosylmethionine (SAM). NSP10-NSP16 complex was incubated with 100 µM inhibitor compound or DMSO as control for 10 min before adding the remaining components. Next, enzymatic reactions were kept for 2 h at 37 °C in 8 µl total volume in 384-well plates (CLS3824, Corning). After incubation, 2 µl of 5x MTase-Glo Reagent (Promega) was added to wells, plates were shaken for 1 min at 1000 rpm and further incubated for 30 min at RT. Then, 10 µl of MTase-Glo Detection Solution (Promega) was added, plates were again shaken for 1 min at 1000 rpm and further incubated for 30 min at RT. Finally, firefly luminescence intensity (∼565 nm) was quantified in a SpectraMax iD5 microplate reader (Molecular Devices).

### Microscale thermophoresis

For microscale thermophoresis (MST) assays, NSP10 protein was fluorescently labeled using *N*-Hydroxysuccinimide (*NHS*)-ester fluorophores according to the manufacturer’s instructions (Protein Labeling Kit RED-NHS 2nd Generation, Nanotemper). Labeling was performed in the NSP16 protein storage buffer (20 mM HEPES pH 7.5 and 0.3 M NaCl). Prior to use in MST experiments, the labeled protein solution (2 µM) was centrifuged at 15.000xg for 10 min at 4°C to remove potential aggregated protein species. NSP16/NSP10 MST experiments were performed in 20 mM HEPES, 0.3 M NaCl with 0.5% Tween, compound/NSP10 experiments in 20 mM HEPES, 0.3 M NaCl with 0.05% Tween. For the binding studies, 100 or 200 nM NSP10-RED (depending on labeling efficiency) were incubated with increasing amounts of NSP16 or compound for 15 min at RT. Measurements were performed in standard capillaries (Nanotemper) using a Monolith MST device (Nanotemper) with 80% LED and medium infrared (IR) laser power. Binding was analyzed from MST signals after 5 or 20 s compared to relative fluorescence before IR laser pulse.

### LuTHy-BRET compound assay

The LuTHy assay procedure was performed as described above using the in-cell BRET readout. After transfection and expression of the LuTHy constructs for 48 h, cells were treated with indicated concentrations of compound dissolved in DMSO infused into the cell culture media. Control wells were treated with DMSO only. After 3 h of compound incubation, fluorescence of mCitrine was recorded at Ex 500 nm/Em 530 nm using the Infinite^®^ microplate reader M1000 PRO (Tecan), and cell morphology (data not shown) was analyzed by automated imaging using a Spark multimode microplate reader (Tecan). Then, coelenterazine-h (pjk, #102182) was added to a final concentration of 5 μM (5 mM stock dissolved in methanol), cells were incubated for an additional 15 min and total luminescence as well as luminescences at short (370-480 nm) and long (520-570 nm) wavelengths were measured using the Infinite^®^ microplate readers M1000 PRO (Tecan). BRET ratios were calculated as described above and normalized to solvent control wells (normalized BRET (nBRET) ratio).

### SARS-CoV-2 replication assay

HEK293-ACE2 (3 × 10^4^ cells per well) were plated in white 96-well plates. The cells were then infected with SARS-CoV-2 (Hou et al, 2020; Kim et al, 2022) (0.01 MOI) containing a nanoluciferase reporter and were simultaneously treated with the compounds (100-0µM concentration). Cells were further cultured for another 24 hours and luminescence was measured (Coutant et al, 2020). Cell Titer-Glo Luminescent Cell Viability Assay kit (Promega, G7750) was used for cell viability.

---

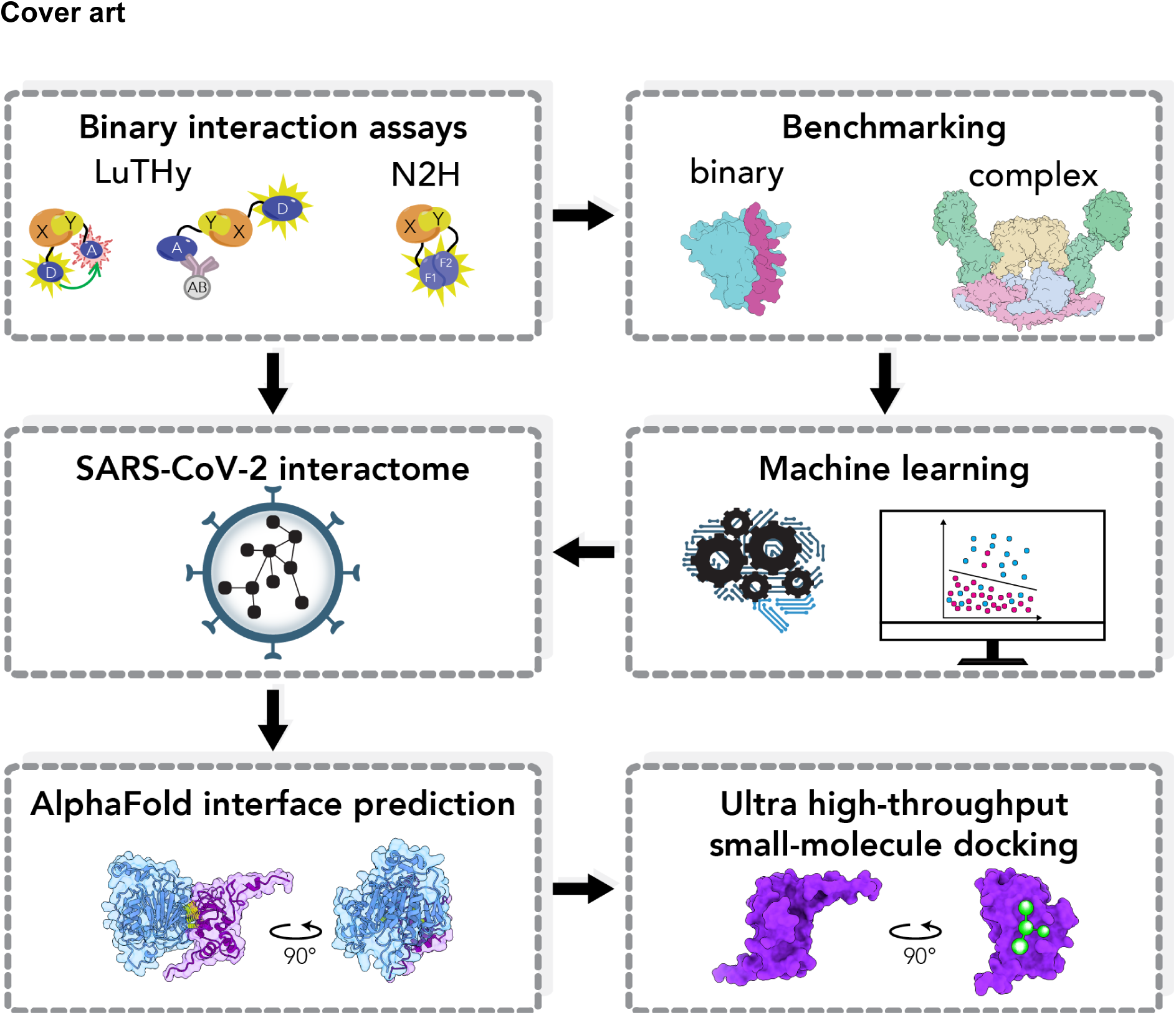

**Figure EV1 (related to Figure 1).**
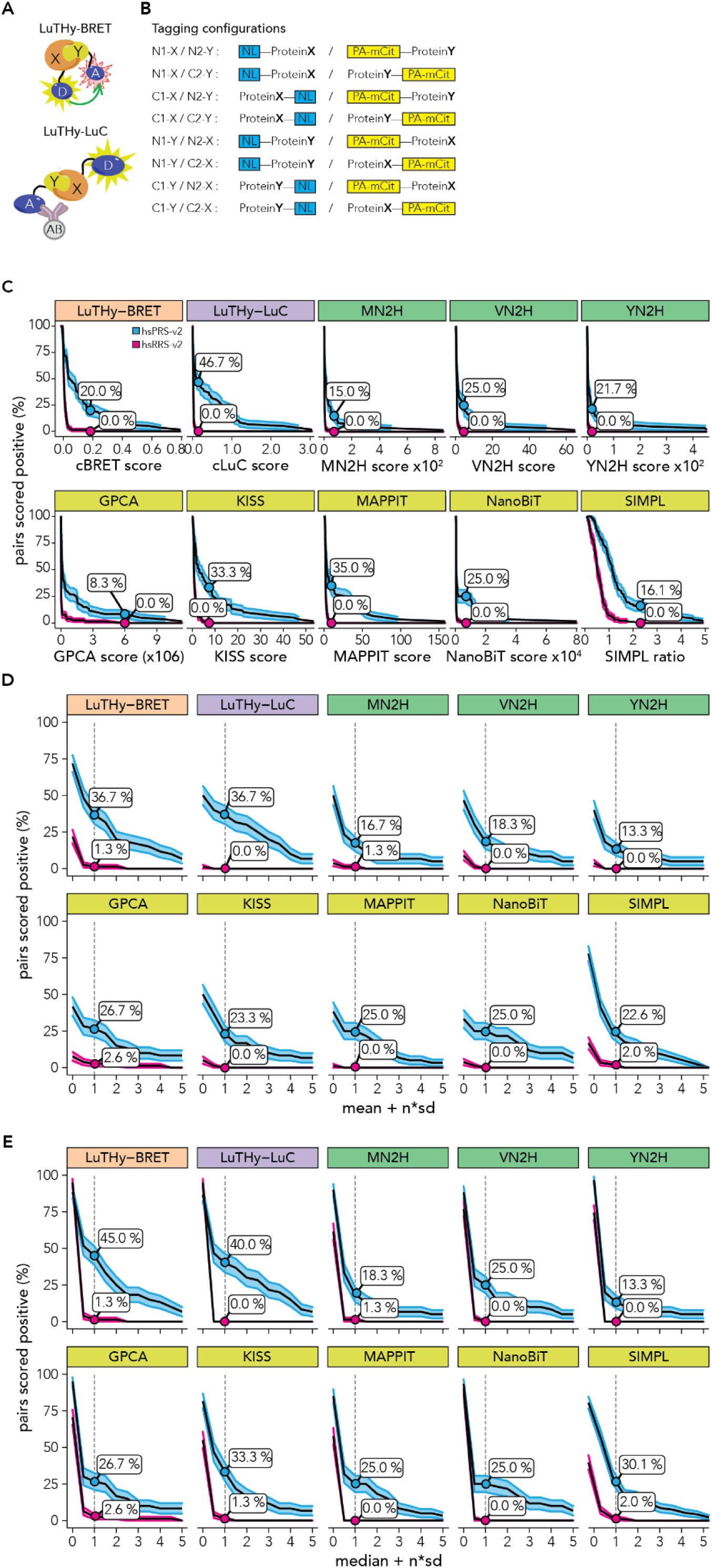
Effect of different scoring approaches on recovery rates. (**A**) Schematic overview of the LuTHy-BRET and LuTHy-LuC assays. X: Protein X, Y: Protein Y, D: NanoLuc donor, A: mCitrine acceptor, AB: antibody. (**B**) With the LuTHy assay, each protein pair X-Y can be tested in eight possible configurations (N- vs. C-terminal fusion for each protein), and proteins can be swapped from one tag to the other resulting in 16 quantitative scores for each protein pair, i.e. eight for LuTHy-BRET and eight for LuTHy-LuC. (**C**) Line plots showing the fraction of protein pairs that scored positive (y-axis) dependent on the quantitative interaction scores (x-axis) for 10 binary PPI assay versions. For each tested protein pair, the tagging configuration with the highest interaction score is used. For LuTHy all eight tagging configurations were tested, whereas for MN2H, VN2H, YN2H, GPCA, NanoBi four and for KISS, MAPPIT and SIMPL two tagging configurations were tested. Recovery rates at maximum specificity, i.e. where none of the protein pairs in the RRS scored positive (0%), are indicated. Note that in Choi et al. (Choi et al, 2019) recovery rates at maximum specificity were calculated by using distinct cut-offs for each tagging configuration. (**D**) Line plots showing the fraction of protein pairs that scored positive (y-axis) dependent on the distribution of interaction scores, i.e. the mean of all interaction scores + n*(sd) (x-axis) for 10 binary PPI assay versions. Recovery rates at mean + 1 standard deviation are indicated (**E**) Line plots showing the fraction of protein pairs that scored positive (y-axis) dependent on the distribution of interaction scores, i.e. the median of all interaction scores + n*(sd) (x-axis) for 10 binary PPI assays. Recovery rates at median + 1 standard deviation are indicated. LuTHy data from this study; SIMPL from Yao et al (Yao et al, 2020); all other from Choi et al (Choi et al, 2019). Note that the SIMPL assay was benchmarked by Yao et al (Yao et al, 2020) against 88 positive proteins pairs derived from the hsPRS-v1 (Venkatesan et al, 2009) and as a random reference set against “88 protein pairs with baits and preys selected from the PRS but used in combinations determined computationally to have low probability of interaction” (Yao et al, 2020).

**Figure EV2 (related to Figure 1).**
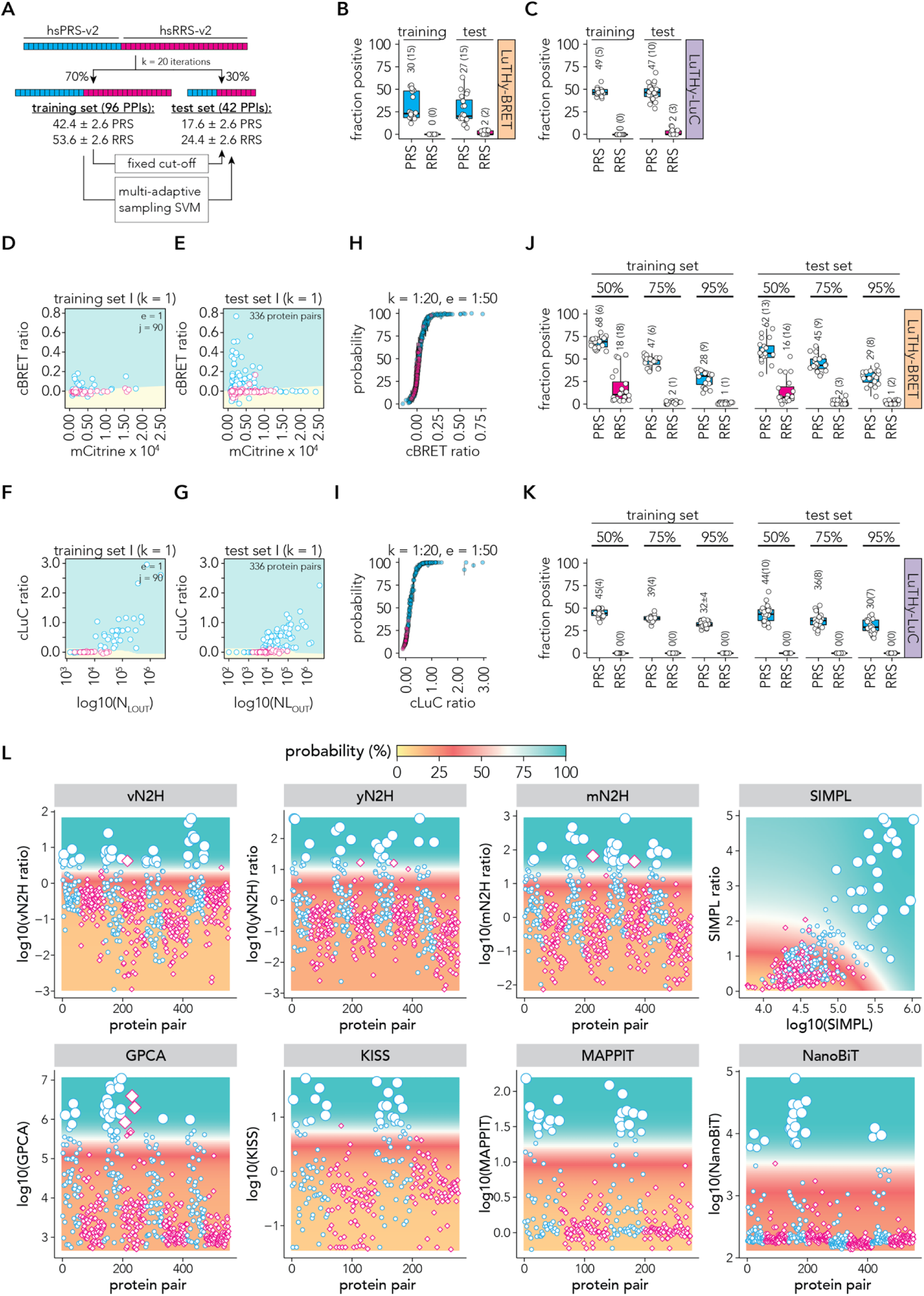
Establishment and performance of the maSVM algorithm. (**A**) Schematic overview to evaluate the performance when using either a fixed cutoff or the maSVM learning algorithm, by separating the hsPRS-v2 and hsRRS-v2 randomly into a training (70%) and a test (30%) set in k = 20 iterations. (**B,C**) Boxplots showing the fraction of protein pairs that scored positive in the training and the test sets (from **A**) using a fixed-cutoff at a maximum specificity, i.e. where none of the interactions of the RRS of the training set scored positive by (**B**) LuTHy-BRET or (**C**) LuTHy-LuC. Each dot represents the recovery rates from one of the 20 iterations. Numbers above boxplot indicate the mean and in brackets the standard deviation. The cutoffs used to determine the fraction of protein pairs that scored positive in the test were used from each respective training set k at 0% RRS. (**D,E**) Representative scatter plot for the first SVM model for the LuTHy-BRET showing in-cell mCitrine expression (x-axis) against cBRET ratios (y-axis) from the (**D**) first training set (k = 1), from which in the first ensemble (e = 1) 90 protein pairs (j) were randomly sampled and reclassified in 5 iterations (i = 5). (**E**) Scatter plot for the first test set (k = 1), that contains 336 protein pair configurations. The classification models from the first SVM model (from **D**) are visualized in (**D,E**) as different colors (negativ = yellow; positive = teal) that are separated by the support vector. The known classification of the protein pairs of the train and test set are indicated by color (blue = PRS; magenta = RRS). (**F,G**) Representative scatter plot for the first SVM model for the LuTHy-LuC showing luminescence after co-precipitation (x-axis) against cLuC ratios (y-axis) from the (**F**) first training set (k = 1), from which in the first ensemble (e = 1) 90 protein pairs (j) were randomly sampled and reclassified in 5 iterations (i = 5). (**G**) Scatter plot for the first test set (k = 1), that contains 336 protein pair configurations. The classification models from the first SVM model (from **F**) are visualized in (**F,G**) as different colors (negativ = yellow; positive = teal) that are separated by the support vector. The known classification of the protein pairs of the train and test set are indicated by color (blue = PRS; magenta = RRS). (**H,I**) Scatter plot showing (**H**) cBRET ratios (x-axis) or (**I**) cLuC ratios (x-axis) against the average classifier probability (y-axis) for all hsPRS-v2 (blue) and hsRRS-v2 (magenta) protein pairs from all eight tagging configurations. The classifier probability was averaged over the twenty assembled training and test sets (k) and standard deviations are indicated. (**J-K**) Box plots showing the fraction of hsPRS-v2 and hsRRS-v2 protein pairs that scored above classifier probabilities of 50%, 75% or 95% by (**J**) LuTHy-BRET and (**K**) LuTHy-LuC. Each dot represents the results from one assembled test and training set (k). Numbers above boxplot indicate the mean and in brackets the standard deviation. (**L**) Scatter plots for eight quantitative PPI assay variants showing the number of protein pairs (x-axis) against their respective interaction scores (y-axis) for hsPRS-v2 (blue) and hsRRS-v2 (magenta) protein pairs. Average classifier probability from the 50 maSVM models is displayed as the size of the data points and as a colored grid in the background. The maSVM algorithm for the SIMPL assay was trained on the SIMPL interaction score (mean SIMPL, x-axis) and the ratio between SIMPL interaction score and bait expression (mean ratio, y-axis) using the published data from Yao et al (Yao et al, 2020). Note that the SIMPL assay was benchmarked against 88 positive proteins pairs derived from the hsPRS-v1 (Venkatesan et al, 2009) and as a random reference set against “88 protein pairs with baits and preys selected from the PRS but used in combinations determined computationally to have low probability of interaction” (Yao et al, 2020). Data for all other assays is from Choi et al (Choi et al, 2019).

**Figure EV3 (related to Figure 1):**
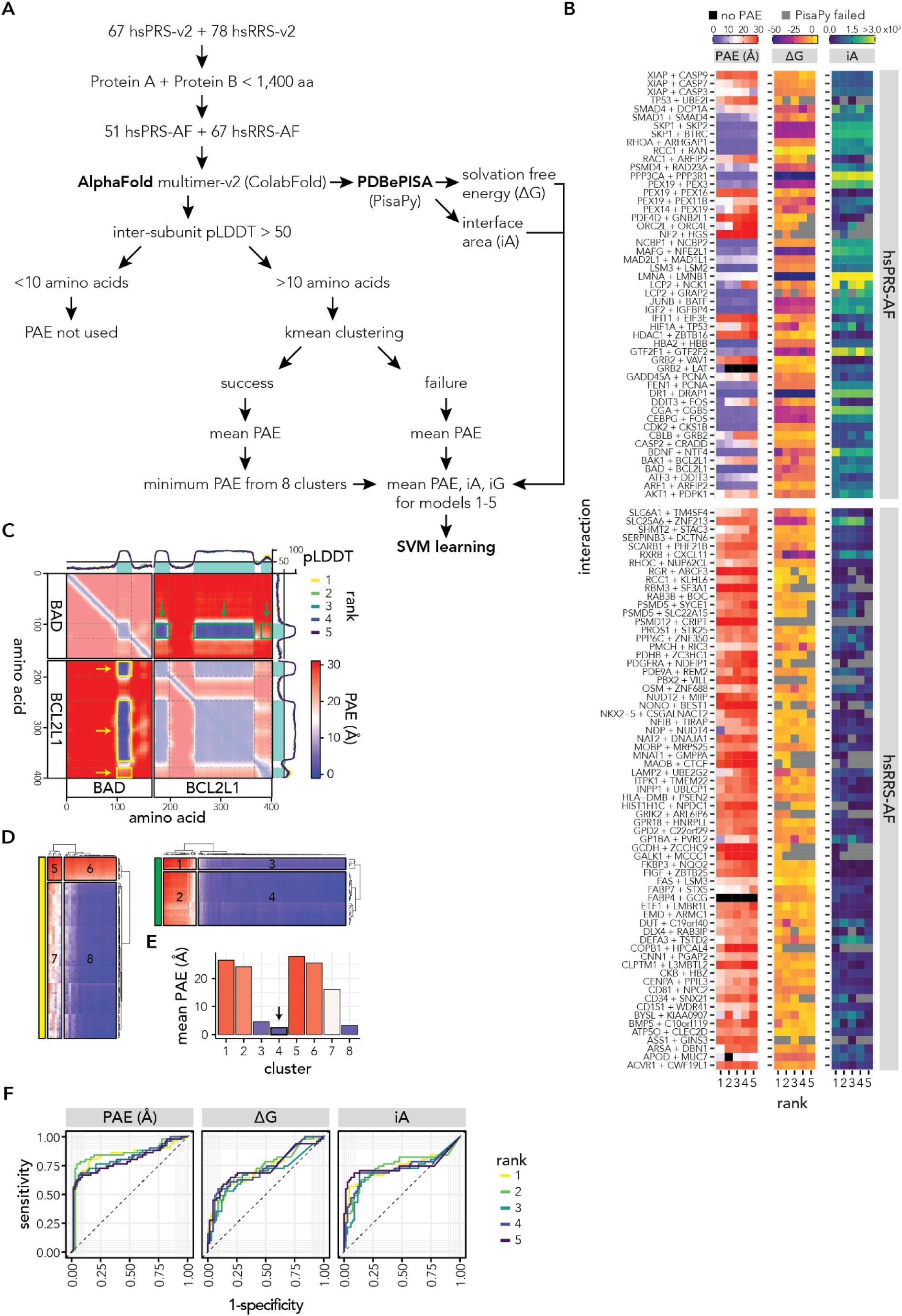
Benchmarking AFM using well-established positive and random reference sets. (**A**) Schematic overview of AlphaFold-multimer (AFM) benchmarking. First, the hsPRS-v2 and hsRRS-v2 were filtered for protein pairs with less than 1,400 amino acids combined, resulting in 51 positive reference set pairs (hsPRS-AF) and 67 random reference set pairs (hsRRS-AF). For these 118 protein pairs, five structural models were predicted using ColabFold through the AFM algorithm (590 total structures). Following, PAE and pLDDT values were extracted from the AFM predicted structures, and inter-subunit amino acids were filtered for pLDDT >50. If >10 inter-subunit amino acids remained, PAE values were k-means clustered. If clustering failed, the mean PAE of the unclustered amino acids was calculated, else the average PAE of the eight clusters were calculated and the minimum PAE selected as the amino acid region with the minimal distance between the two proteins. In addition, PDBePISA was used to determine the solvation free energy (ΔG) and the area (iA) of the interface region (Python script PisaPy was used for batch analysis) for 521 of the 590 structures. For the remaining 69 structures PDBePISA could not identify an interface. Next, the average PAE, iA and ΔG were calculated for the five predicted structural models of the 51 hsPRS-AF and 67 hsRRS-AF protein pairs. Finally, a multi-adaptive maSVM learning algorithm was trained on the PAE and iA features of the hsPRS-AF and hsRRS-AF as outlined in Figure 1A. The 50 trained models were ultimately used to predict the classifier probability of the CoV-2-AF structures in Figure EV7B-E. (**B**) Heatmap of the PAEs, ΔGs and iAs for protein pairs of the hsPRS-AF and hsRRS-AF. Shown are the minimum PAE values after kmeans clustering. If <10 amino acids had pLDDT >50 the PAE values were not used and shown in black. Protein pairs where no interaction interface was detected by PDBePISA are shown in gray. (**C**) Representative example for the kmeans clustering strategy of AFM reported PAE values. Heatmap shows the PAEs for the protein pair BAD+BCL2L1 (hsPRS-AF) rank 1 model. The intra-molecular PAEs are shown with 50% opacity. The predicted local distance difference test (pLDDT) for all five predicted models (rank 1-5) are shown as line graphs on top and on the right of the heatmap. The area with pLDDT scores >50 is highlighted in teal. Inter-molecular regions with pLDDT >50 that were used for kmeans clustering are highlighted with green (BAD>BCL2L1 interface) and yellow (BCL2L1>BAD interface) boxes that are also indicated with arrows. (**D**) Clustering results of green and yellow regions highlighted in panel C. Cluster numbers are indicated. (**E**) Average PAE values for the eight clusters from panels C and D. The arrow indicates the cluster with the lowest average PAE value. (**F**) Receiver characteristic analysis comparing sensitivity and specificity between the five ranked structural models for PAE, ΔG and iA of the hsPRS-AF and hsRRS-AF.

**Figure EV4 (related to Figure 1):**
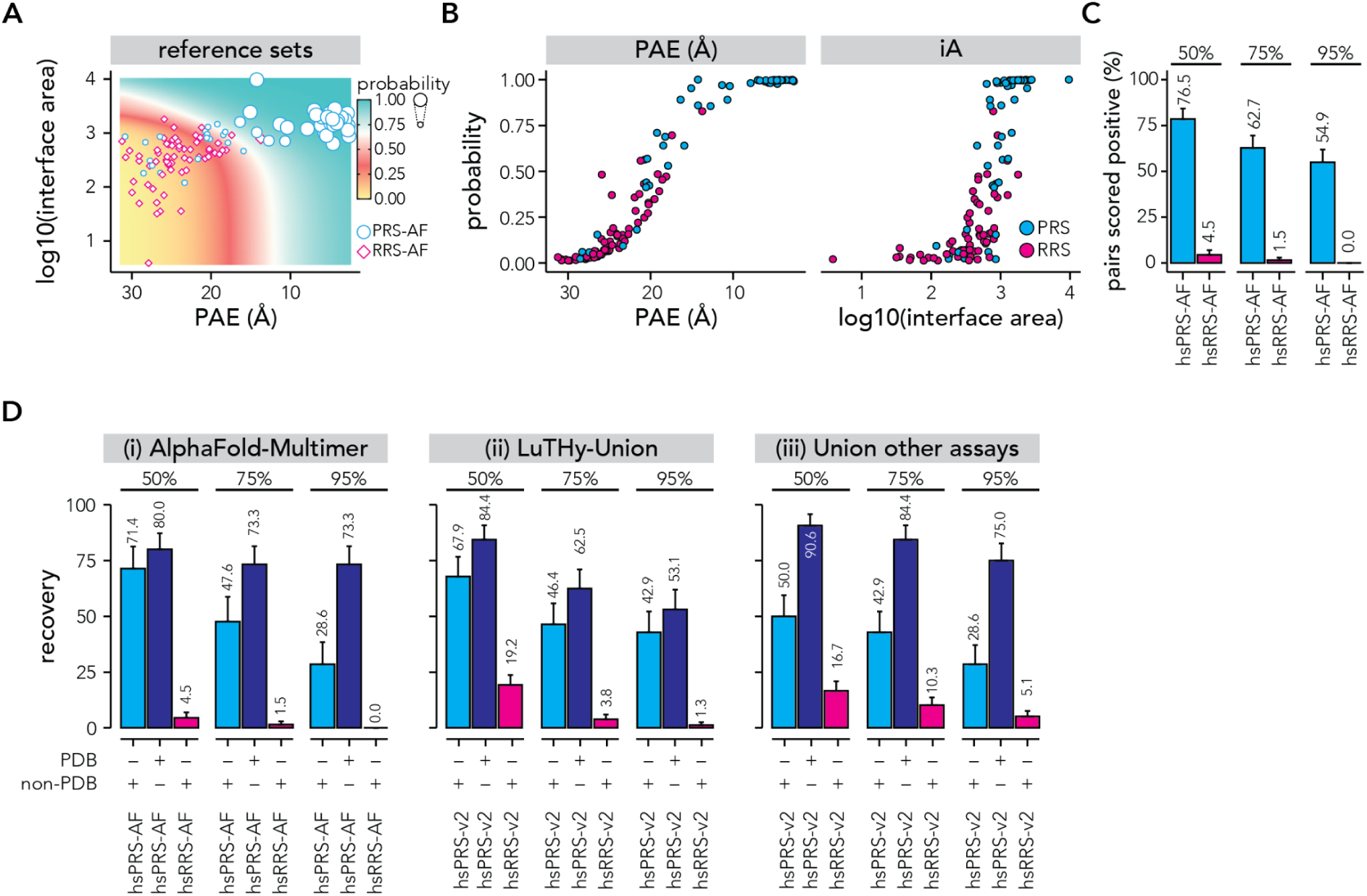
Training a maSVM algorithm to classify AFM predicted structures. (**A**) Scatter plot showing PAE (x-axis) against interface area (y-axis) for all hsPRS-AF (blue) and hsRRS-AF (magenta) protein pairs. Average classifier probability from the 50 maSVM models is displayed as the size of the data points and as a colored grid in the background. (**B**) Scatter plots showing PAE (x-axis, left panel) or interface area (x-axis, right panel) against classifier probability (y-axis) for all hsPRS-AF (blue) and hsRRS-AF (magenta) protein pairs. (**C**) Bar plots showing the fraction of hsPRS-AF and hsRRS-AF protein pairs that scored above classifier probabilities of 50%, 75% and 95%. (**D**) Bar plots showing the fraction of hsPRS-AF and hsRRS-AF interactions with structures deposited in PDB that scored above classifier probabilities of 50%, 75% and 95% by AlphaFold-Multimer (i) and the fraction of hsPRS-v2 and hsRRS-v2 interactions with structures deposited in PDB that scored above classifier probabilities of 50%, 75% or 95% by LuTHy (ii) or the union of five other binary assays (iii), N2H (MN2H, VN2H, YN2H), GPCA, KISS, MAPPIT and NanoBiT. Data for the SIMPL assay was excluded for this analysis due to the different composition of the reference sets.

**Figure EV5 (related to Figure 2).**
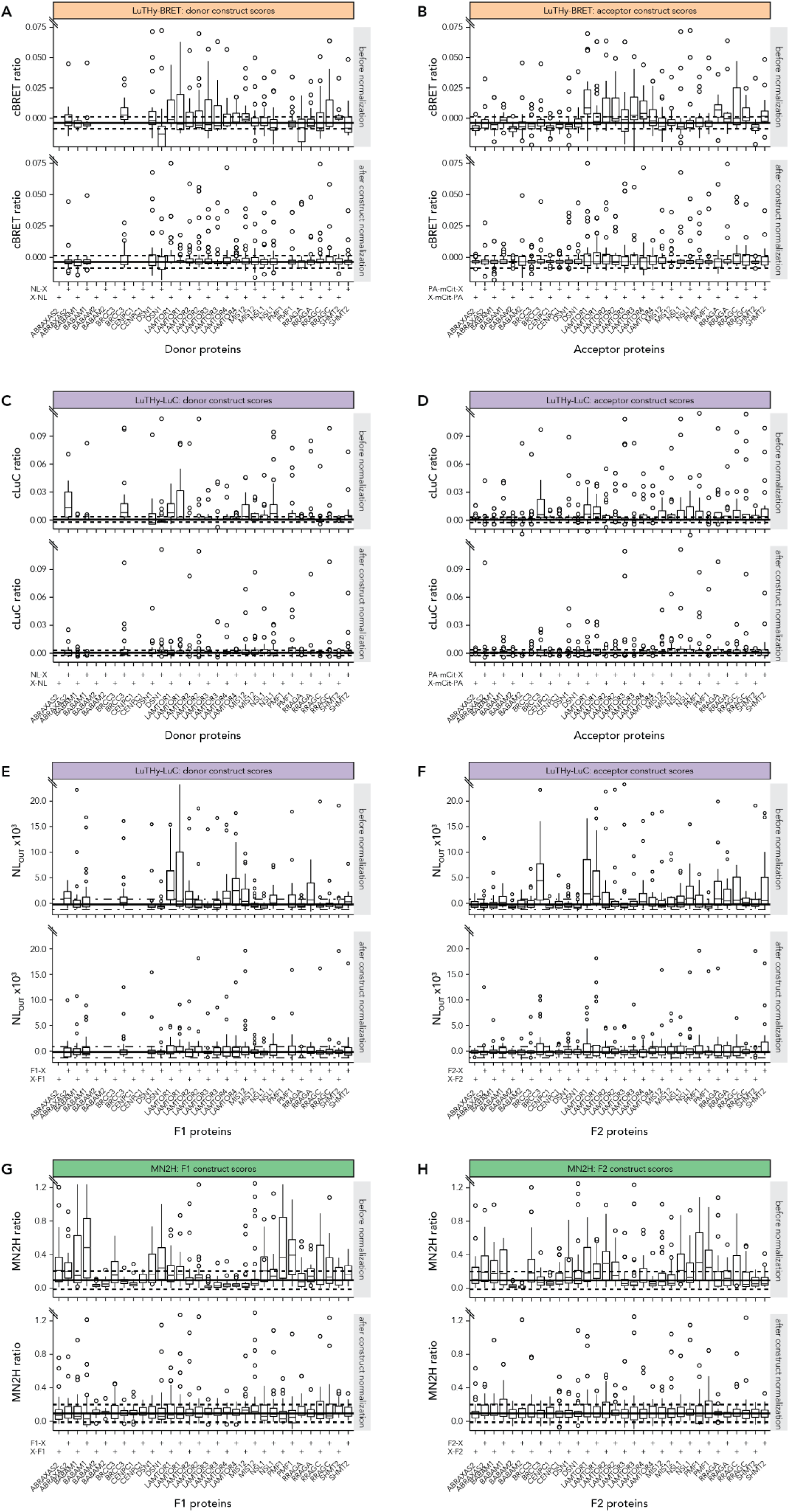
Construct-specific robust scaler normalization for mapping multiprotein complexes. (**A-H**) Boxplots of assay specific interaction scores before and after robust scaler normalization for the multiprotein complex proteins in all tagging configurations of donor and acceptor constructs. Boxplots display the constructs’ median, lower and upper hinges the 25^th^ and 75^th^ percentiles, lower and upper whiskers extending from the hinges with 1.5x the inter-quartile range and outlier points beyond the end of the whiskers. The thick horizontal line indicates the median interaction score over all constructs of the multiprotein complex set and the dashed lines the respective IQR of the 25^th^ and 75^th^ quartiles. Note that the horizontal lines always refer to the median and IQR before normalization and that the range of the y-axis is limited to visualize all boxplots as well as the median and IQR, but high scoring protein pairs (outliers) are hidden. (**A**) cBRET ratios for donor constructs before (top) and after (bottom) robust scaler normalization. (**B**) cBRET ratios for acceptor constructs before (top) and after (bottom) robust scaler normalization. (**C**) cLuC ratios for donor constructs before (top) and after (bottom) robust scaler normalization. (**D**) cLuC ratios for acceptor constructs before (top) and after (bottom) robust scaler normalization. (**E**) Luminescence after co-precipitation (NL_OUT_) for donor constructs before (top) and after (bottom) robust scaler normalization. (**F**) Luminescence after co-precipitation (NL_OUT_) for acceptor constructs before (top) and after (bottom) robust scaler normalization. (**G**) mN2H ratios for F1 constructs before (top) and after (bottom) robust scaler normalization. (**H**) mN2H ratios for F2 constructs before (top) and after (bottom) robust scaler normalization.

**Figure EV6 (related to Figure 3).**
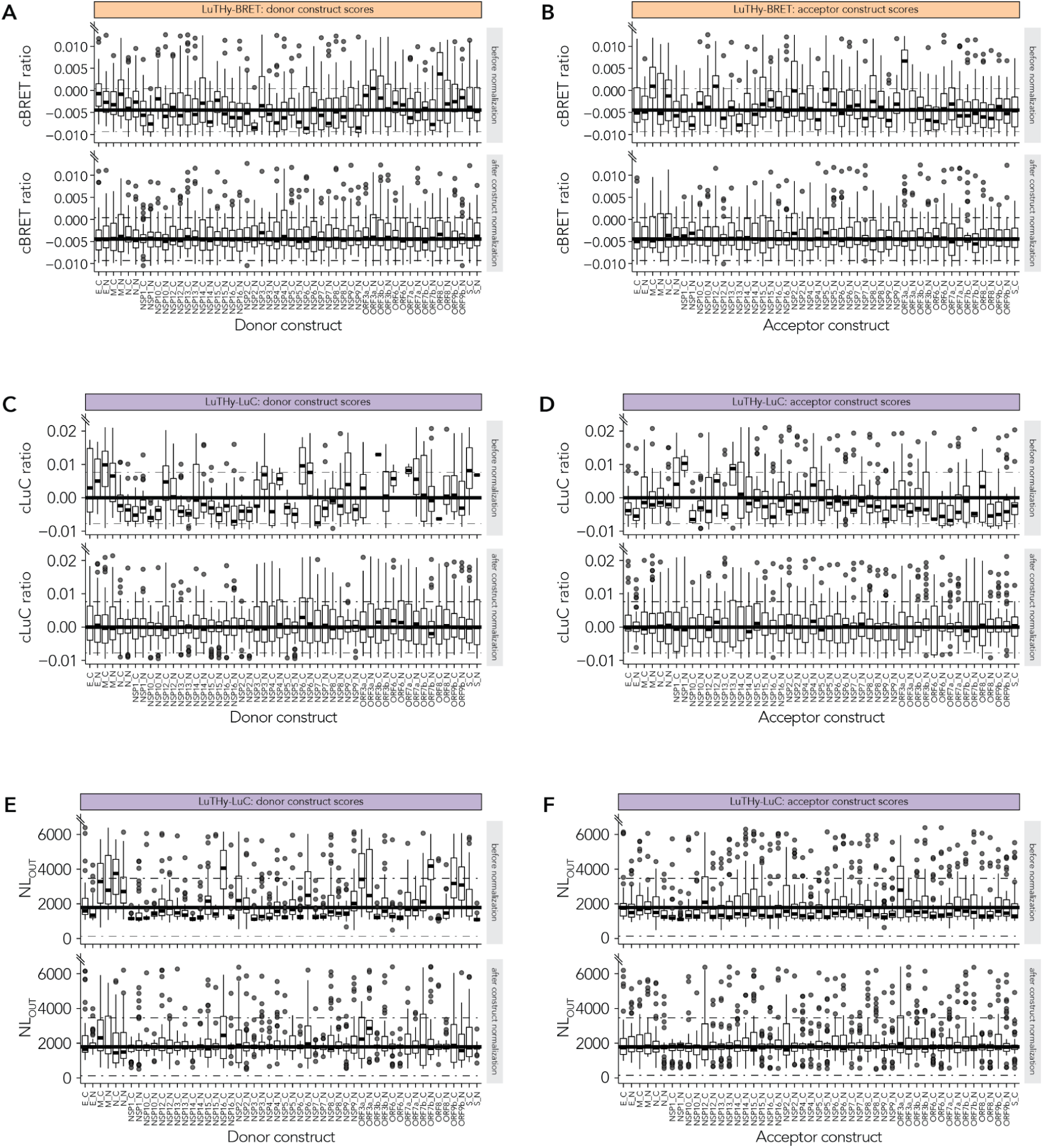
Construct-specific robust scaler normalization for SARS-CoV-2 binary PPI mapping. (**A-F**) Boxplots of LuTHy-BRET and LuTHy-LuC interaction scores before and after robust scaler normalization for the SARS-CoV-2 proteins in all tagging configurations of donor and acceptor constructs. Boxplots display the constructs’ median, lower and upper hinges the 25^th^ and 75^th^ percentiles, lower and upper whiskers extending from the hinges with 1.5x the inter-quartile range and outlier points beyond the end of the whiskers. The thick horizontal line indicates the median interaction score over all constructs of the training (hsPRS-v2, hsRRS-v2, multiprotein complex) and test set (SARS-CoV-2) and the dashed lines the respective IQR of the 25^th^ and 75^th^ quartiles. Note that the horizontal lines always refer to the median and IQR before normalization and that the range of the y-axis is limited to visualize all boxplots as well as the median and IQR, but high scoring protein paris (outliers) are hidden. (**A**) cBRET ratios for donor constructs before (top) and after (bottom) robust scaler normalization. (**B**) cBRET ratios for acceptor constructs before (top) and after (bottom) robust scaler normalization. (**C**) cLuC ratios for donor constructs before (top) and after (bottom) robust scaler normalization. (**D**) cLuC ratios for acceptor constructs before (top) and after (bottom) robust scaler normalization. (**E**) Luminescence after co-precipitation (NL_OUT_) for donor constructs before (top) and after (bottom) robust scaler normalization. (**F**) Luminescence after co-precipitation (NL_OUT_) for acceptor constructs before (top) and after (bottom) robust scaler normalization.

**Figure EV7 (related to Figure 5 and Figure EV5).**
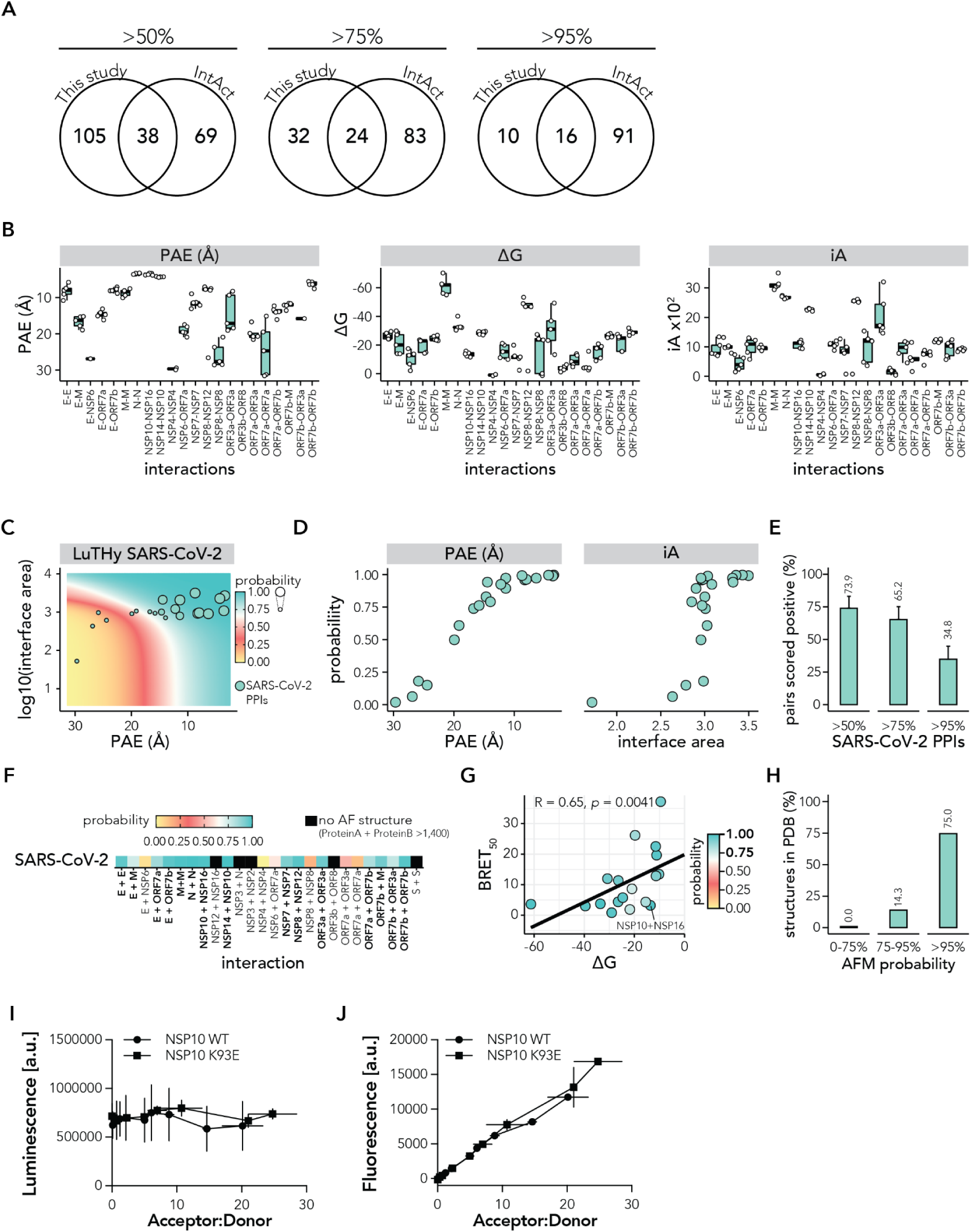
Predicting SARS-CoV-2 protein complex structures using AlphaFold Multimer. (**A**) Venn diagrams showing the overlap between interactions recovered by LuTHy at >50%, >75% and >95% probabilities and interactions deposited in the IntAct database (Orchard et al, 2014) (**B**) Boxplots showing predicte d alignment error (PAE), solvation free energy (ΔG) and interface area (iA) from AlphaFold-Multimer (AFM) predicted SARS-CoV-2-AF structures. Boxplots display the median, lower and upper hinges the 25^th^ and 75^th^ percentiles and lower and upper whiskers extending from the hinges with 1.5x the inter-quartile range. Each dot represents one predicted structural model. (**C**) Scatter plot showing PAE (x-axis) against interface area (y-axis) for all SARS-CoV-2-AF (orange) protein pairs. Average classifier probability predicted by the 50 maSVM models trained by the hsPRS-AF and hsRRS-AF set (see Figure EV4A), is displayed as the size of the data points. Each point in the colored grid in the background displays the average classifier probabilities from the 50 maSVM models. (**D**) Scatter plots showing PAE (x-axis, left panel) or interface area (x-axis, right panel) against classifier probability (y-axis) for all SARS-CoV-2-AF (orange) protein pairs. (**E**) Bar plots showing the fraction of SARS-CoV-2-AF protein pairs that scored above classifier probabilities of 50%, 75% and 95%. (**F**) Heatmap showing the classifier probabilities for the AFM predicted protein pair structures of the SARS-CoV-2-AF protein pairs. (**G**) Scatter plot showing the average ΔG (x-axis) from the five predicted structural SARS-CoV-2-AF models against the LuTHy-BRET determined binding strengths (BRET_50_, see Figure EV8A). Only SARS-CoV-2 predicted AFM structures with classifier probabilities of >75% are shown and the respective classifier probabilities are indicated by the fill color of the data points. A linear regression fit through the data is shown and the Spearman correlation coefficient (R) and p-value are indicated. (**H**) Barplot showing the fraction of AFM predicted structures with 0-75%, 75-95% and >95% classification probability that have an experimentally reported structure deposited to the PDB (Berman et al, 2000) database. (**I,J**) Luminescence (**I**) and fluorescence (**J**) values from LuTHy-BRET donor saturation experiment, where constant amounts of NSP10-NL wt or K93E are co-expressed with increasing amounts of mCitrine-NSP16.

**Figure EV8 (related to Figure 4 and Figure EV7).**
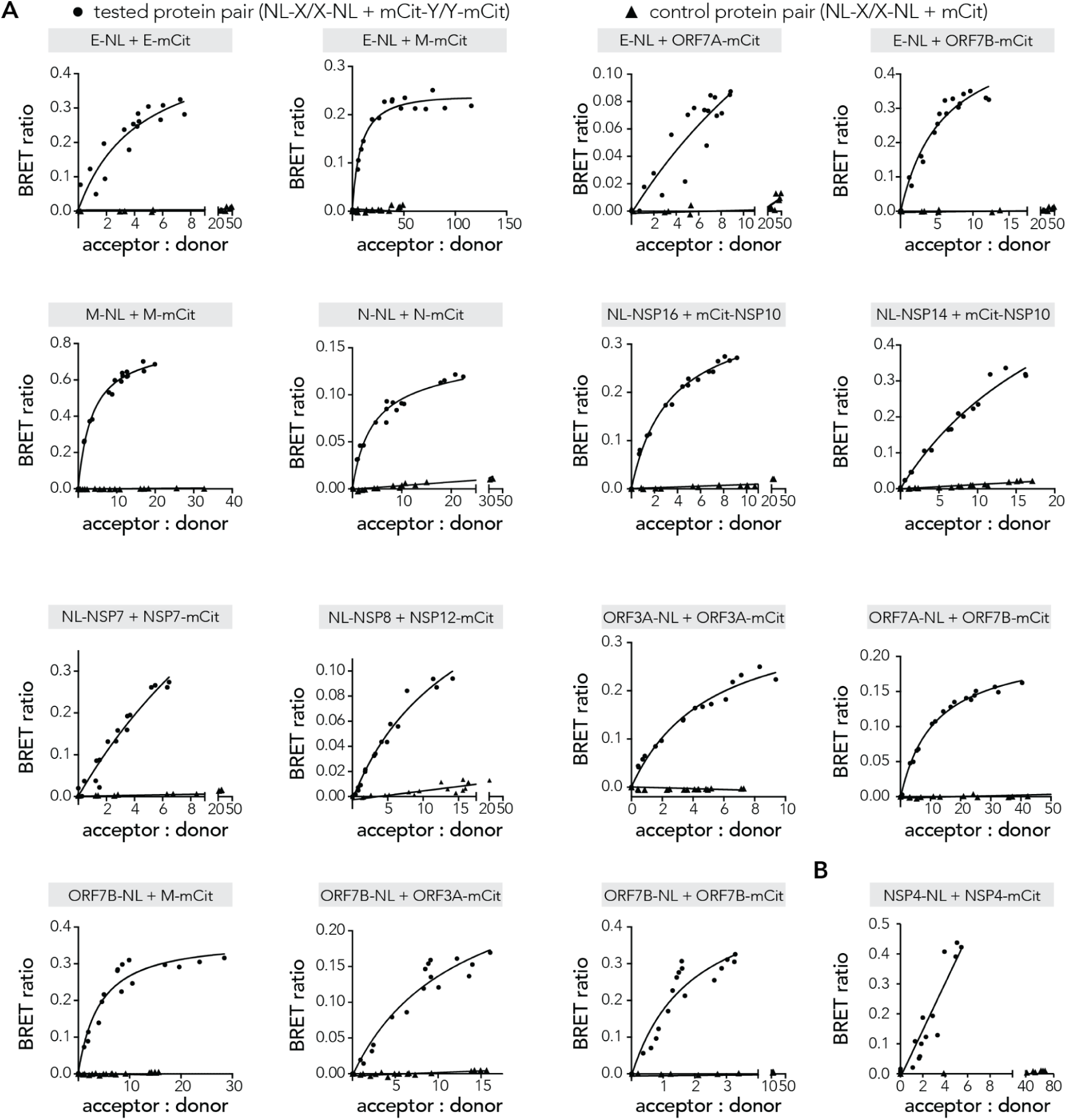
LuTHy-BRET binding strengths for CoV-2-AF protein complexes. (**A,B**) Scatter plots showing LuTHy-BRET donor saturation curves with the acceptor to donor ratio (x-axis) plotted against the BRET ratio (y-axis) for 16 CoV-2-AF protein pairs with classifier probabilities >75%. A non-linear regression fit was performed using the ‘One site – Total and nonspecific binding’ in GraphPad Prism, using the results from the ‘NL-X/X-NL + mCit’ protein pair to subtract nonspecific binding in order to calculate the acceptor to donor ratios at half-maximal BRET ratios (BRET_50_). BRET_50_ values for protein pairs in (**A**) were used in Figure EV7F. (**B**) For the homodimer between NSP4-NSP4 the calculation of a BRET_50_ failed due to the linear relation between acceptor : donor and the BRET ratio. The linear relation can be the result of an unspecific binding between the two proteins or because of a higher order oligomerization of the protein.

## APPENDIX

**Source Data Figure 1**: Raw LuTHy data for interactions in hsPRS-v2 and hsRRS-v2.

**Source Data Figure 2**: Raw LuTHy and mN2H data for the multiprotein complexes.

**Source Data Figure 3**: Raw LuTHy data for SARS-CoV-2.

**Source Data Figure 4**: AlphaFold-Multimer (ColabFold) predicted structures for SARS-CoV2-AF protein pairs and PDBePISA calculated interface information.

**Source Data Figure EV2:** Randomly split hsPRS-v2 and hsRRS-v2 into training (70%) and a test (30%) set.

**Source Data Figure EV3 and EV4:** AlphaFold-Multimer (ColabFold) predicted structures for hsPRS-AF and hsRRS-AF protein pairs and PDBePISA calculated interface information. Due to file size limitation, it contains mostly .pdb files from AFM predictions, whereas .json and .a3m files were removed. Available upon request.

**Supplementary Table 1:** hsPRS-v2 interactions supported by structures or homologous structures

**Supplementary Table 2:** List of the multiprotein complexes meeting the prioritization criteria in this study.

**Supplementary Table 3:** SARS-CoV-2 interactions in IMEx not detected with LuTHy.

**Supplementary Table 4:** LuTHy identified SARS-CoV-2 interactions with >50% interaction probability.

**Supplementary Table 5:** LuTHy identified SARS-CoV-2 interactions with >95% interaction probability.

