## Supplementary figures and images for "AI-guided pipeline for protein-protein interaction drug discovery identifies a SARS-CoV-2 inhibitor"

### AKT1_PDPK1_coverage.png

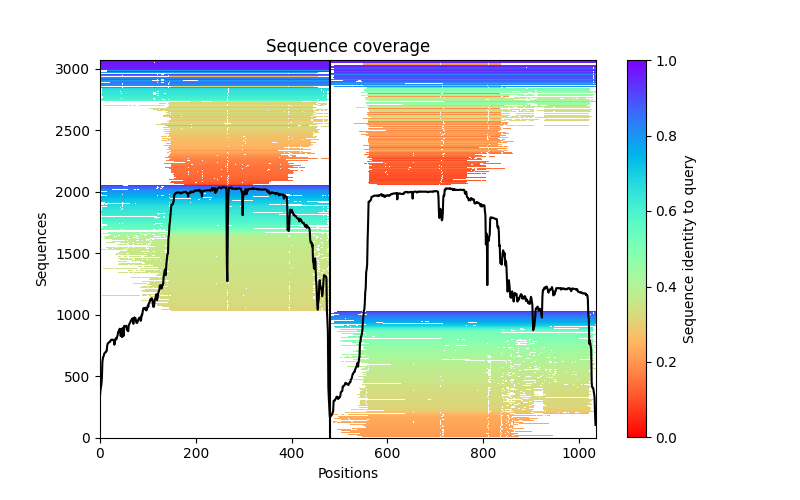

### AKT1_PDPK1_PAE.png

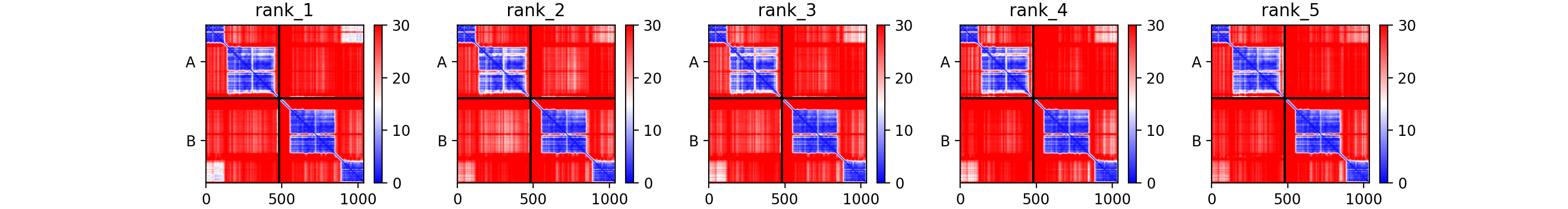

### AKT1_PDPK1_plddt.png

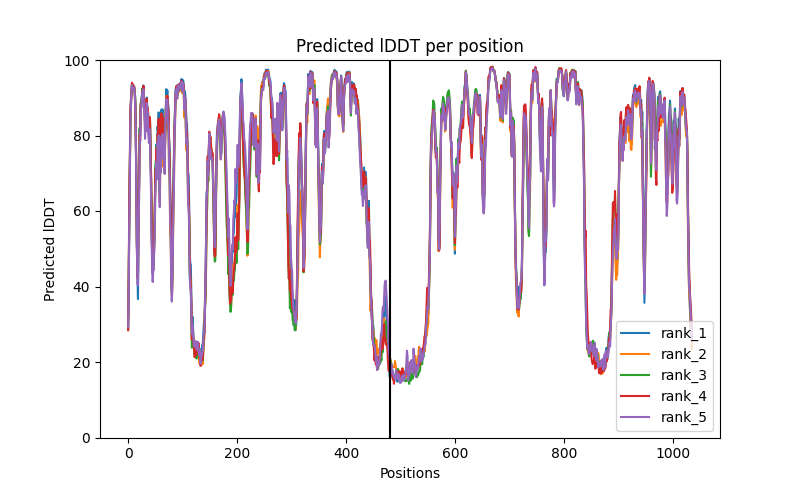

### APOD_MUC7_PAE.png

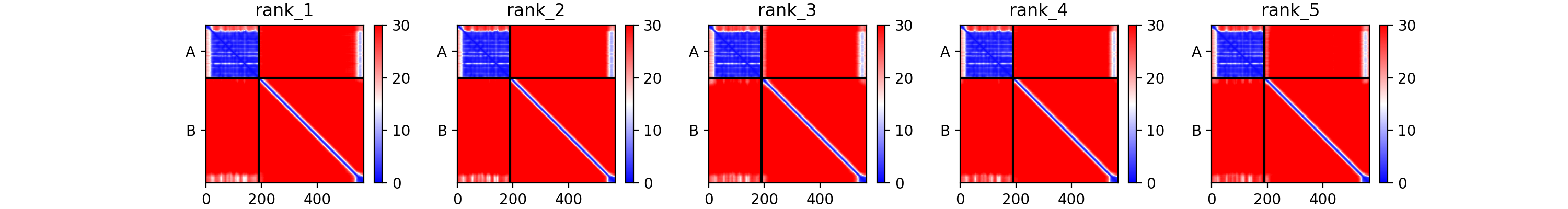

### APOD_MUC7_plddt.png

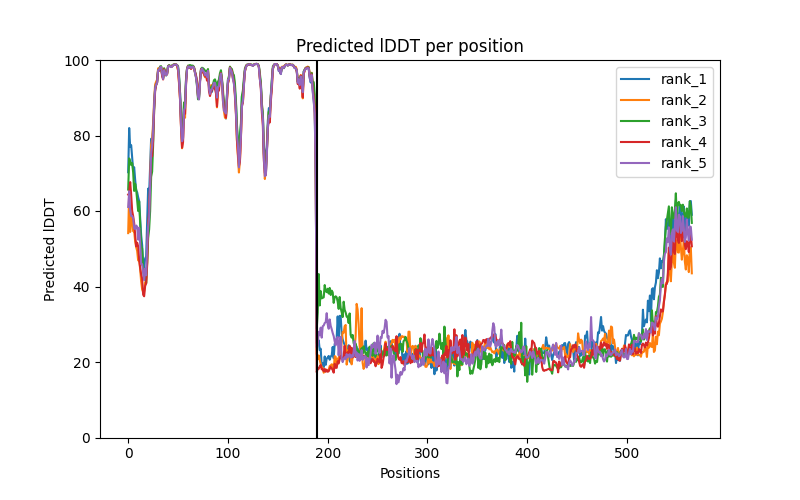

### ARF1_ARFIP2_plddt.png

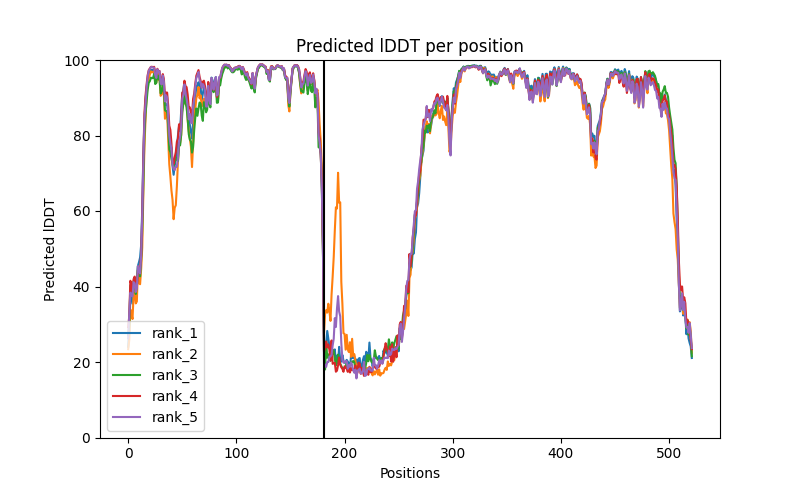

### ARSA_DBN1_coverage.png

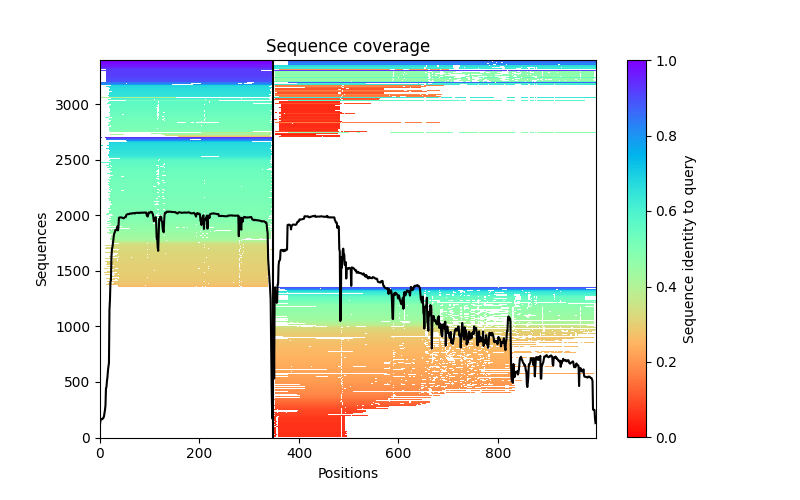

### ATF3_DDIT3_PAE.png

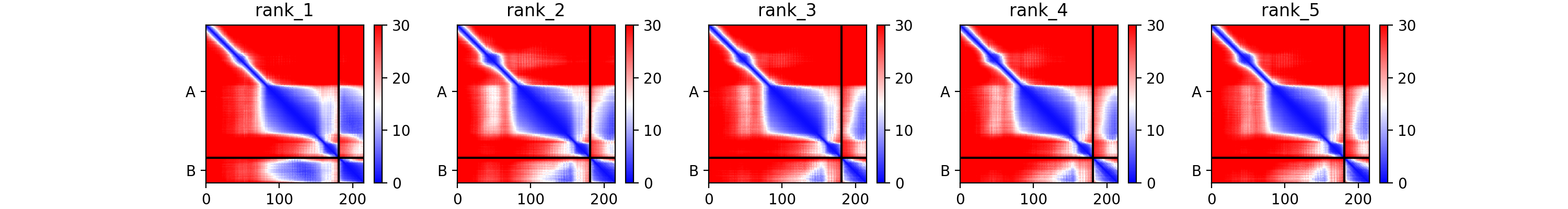

### BAD_BCL2L1_coverage.png

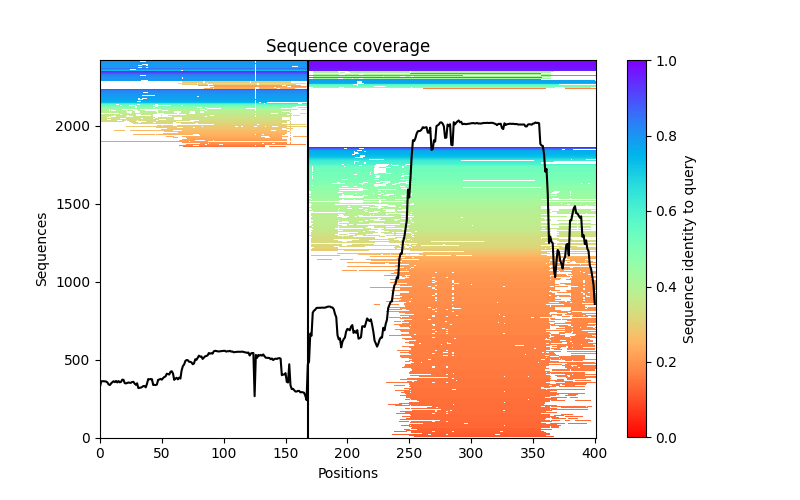

### BYSL_KIAA0907_PAE.png

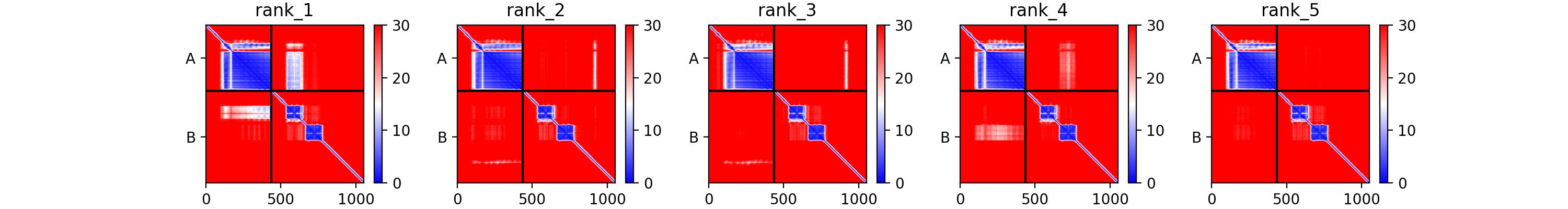

### CD151_WDR41_coverage.png

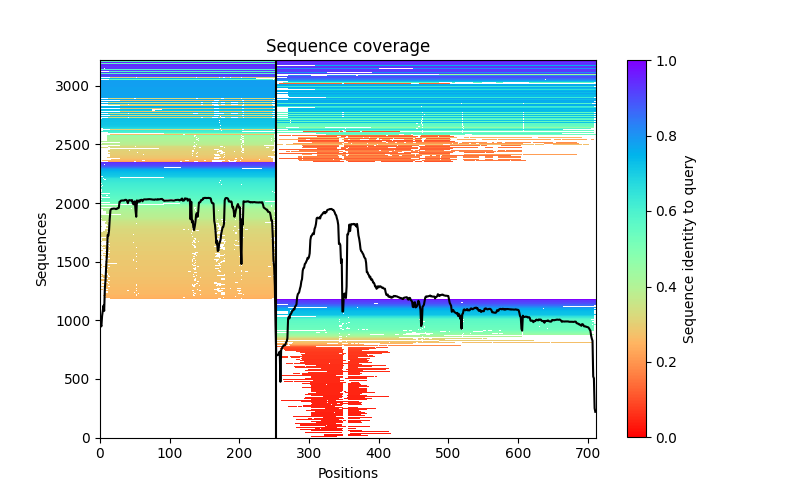

### CGA_CGB5_coverage.png

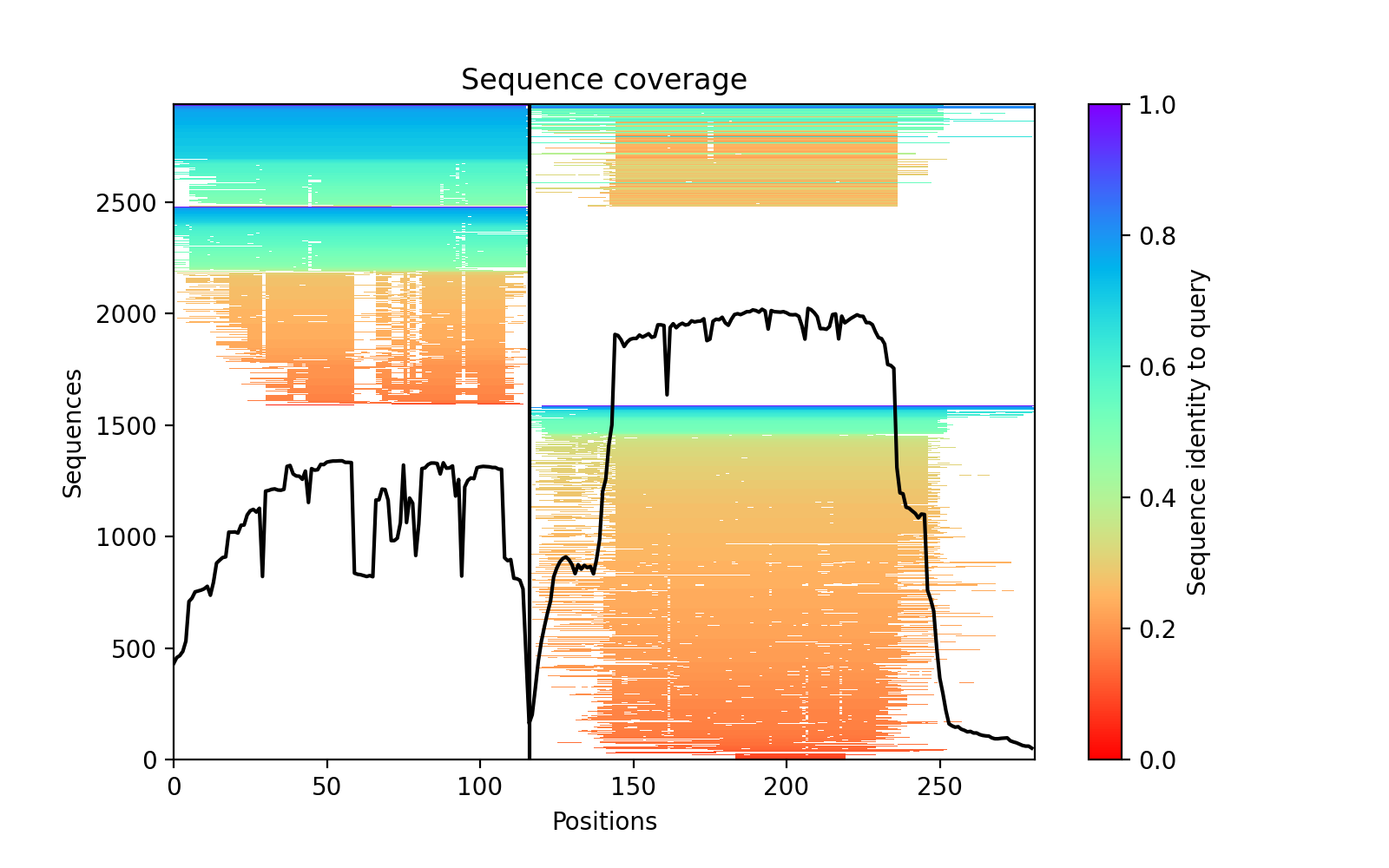

### COPB1_HPCAL4_PAE.png

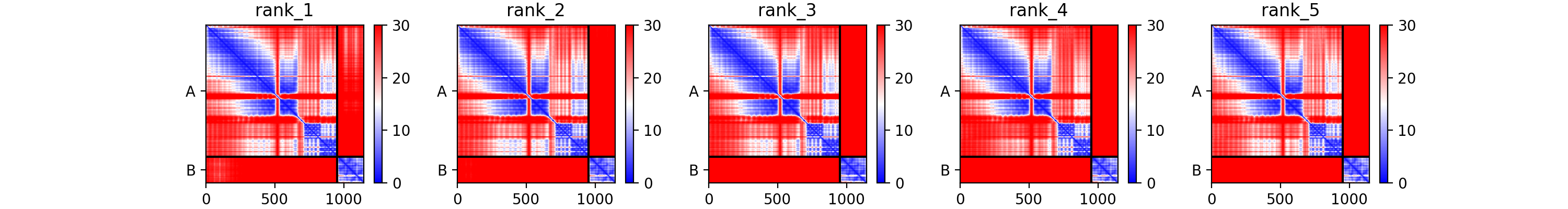

### DEFA3_TSTD2_PAE.png

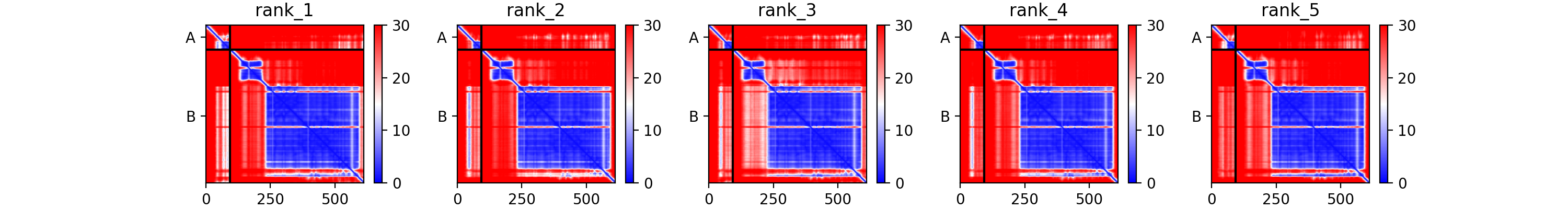

### DLX4_RAB3IP_PAE.png

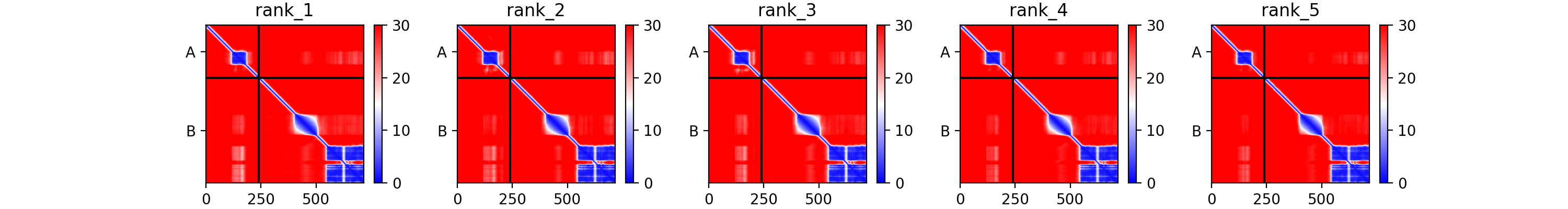

### DUT_C19orf40_PAE.png

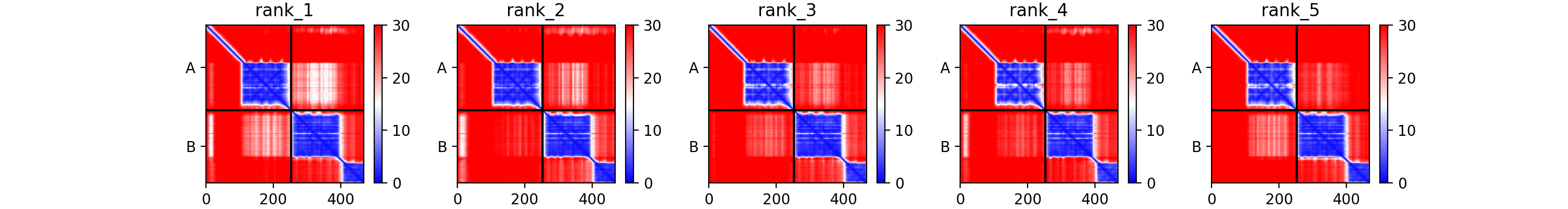

### E_E_420f7_coverage.png

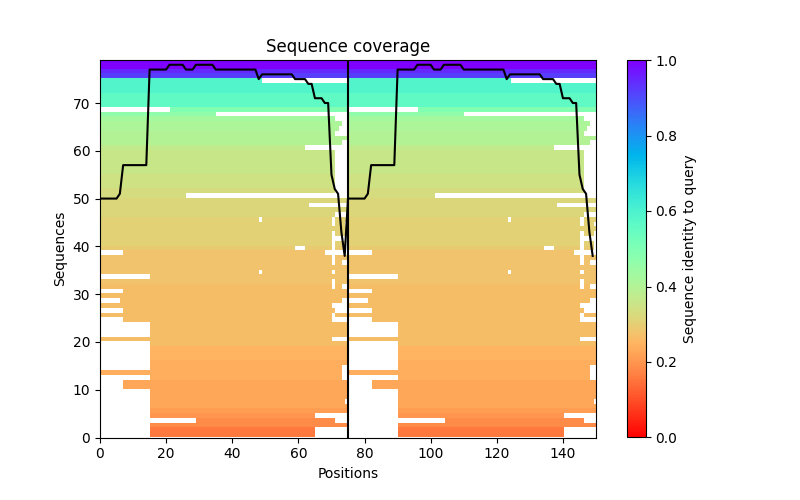

### E_NSP6_9c8a7_coverage.png

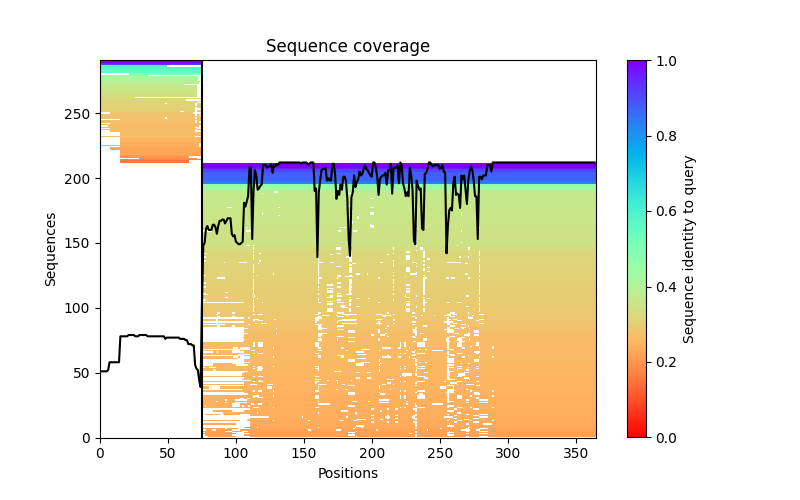

### E_NSP6_9c8a7_PAE.png

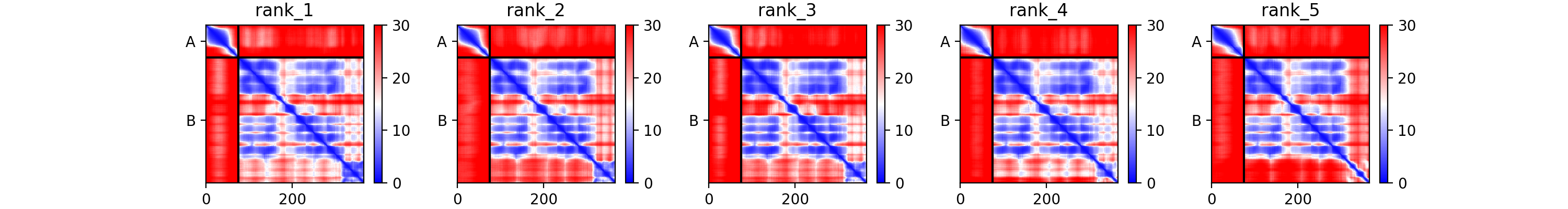

### E_NSP6_9c8a7_plddt.png

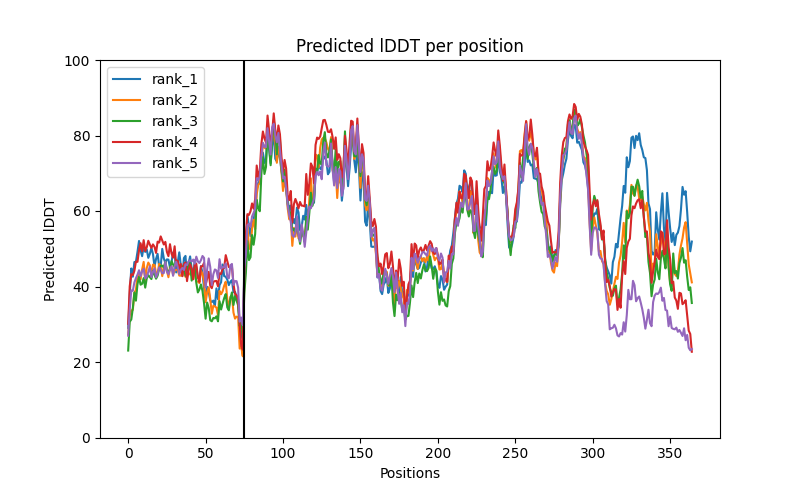

### E_ORF7a_764d1_coverage.png

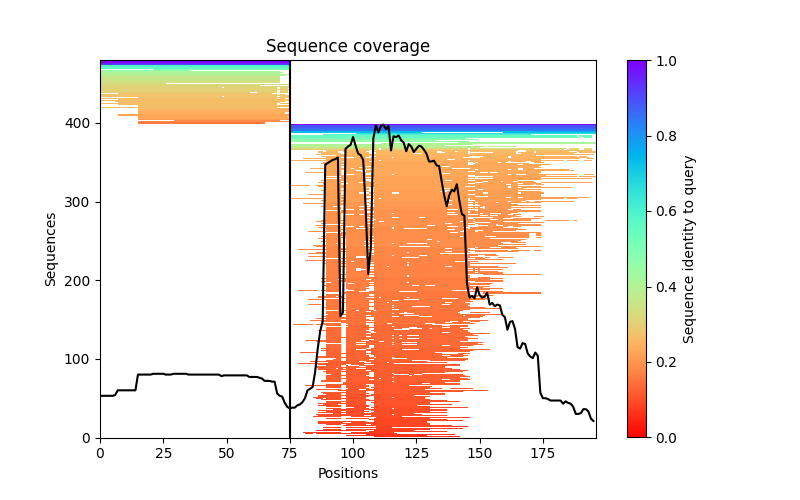

### E_ORF7a_764d1_PAE.png

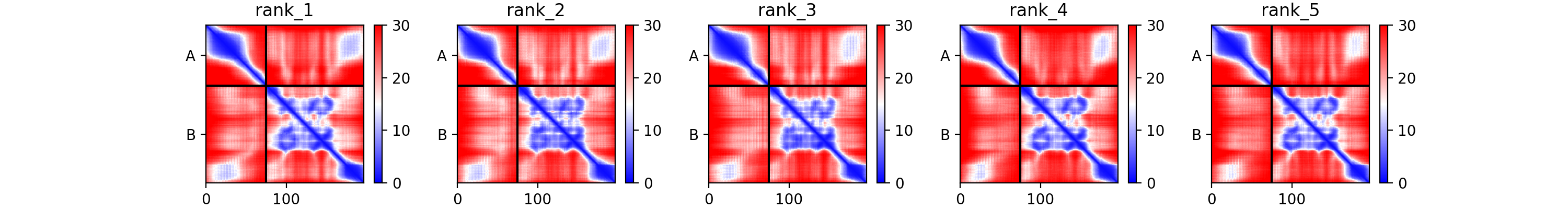

### E_ORF7a_764d1_plddt.png

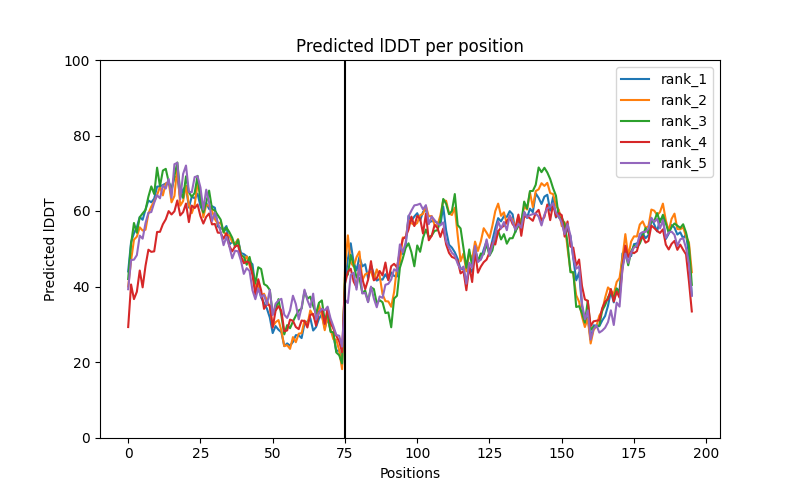

### E_ORF7b_a7aff_coverage.png

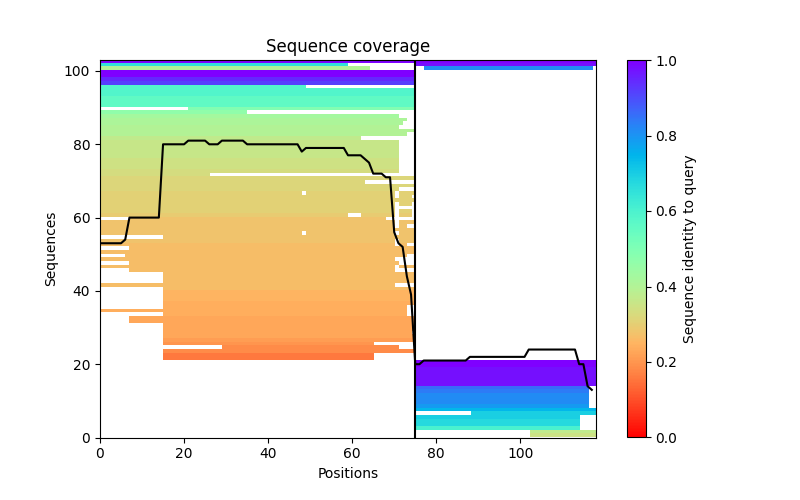

### E_ORF7b_a7aff_PAE.png

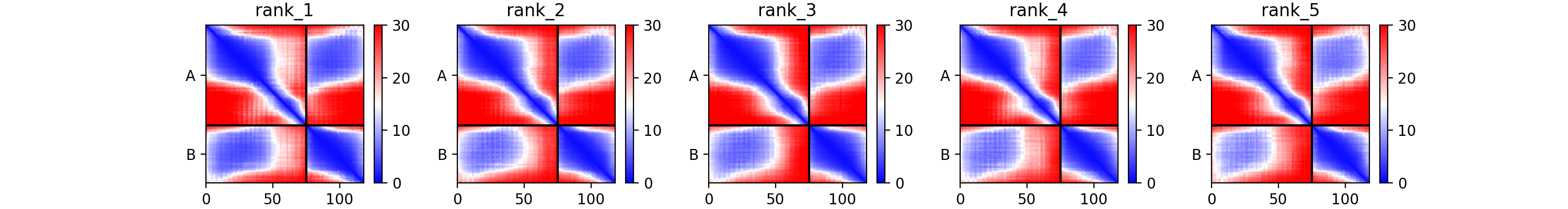

### E_ORF7b_a7aff_plddt.png

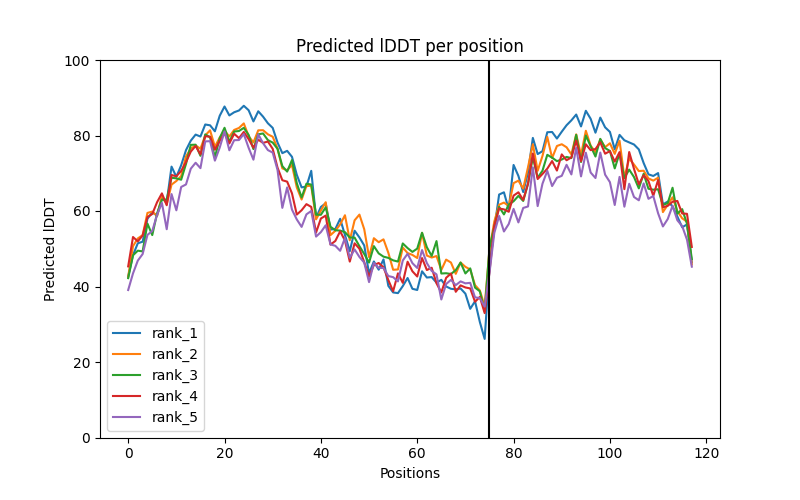

### ETF1_LMBR1L_coverage.png

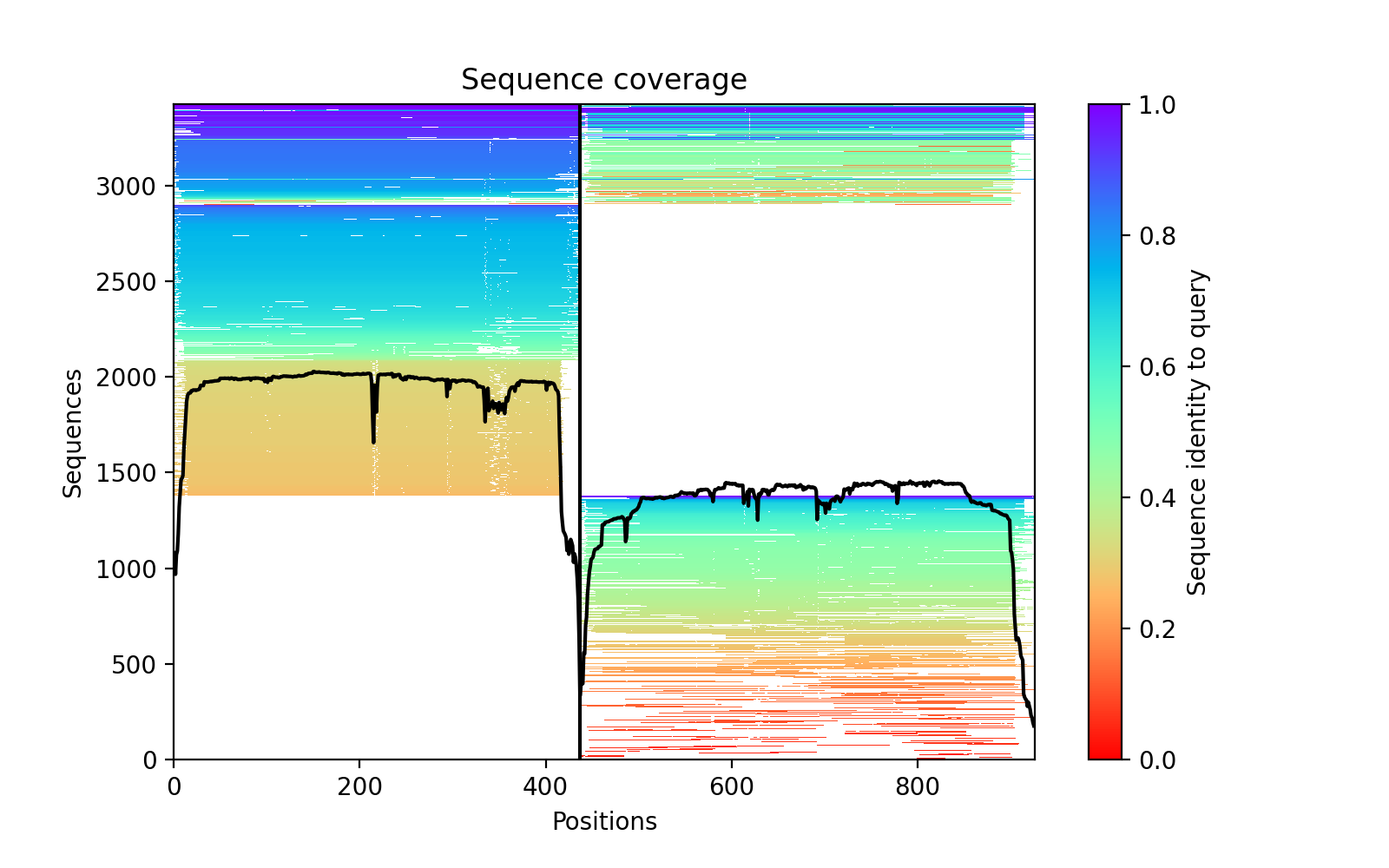

### FABP4_GCG_plddt.png

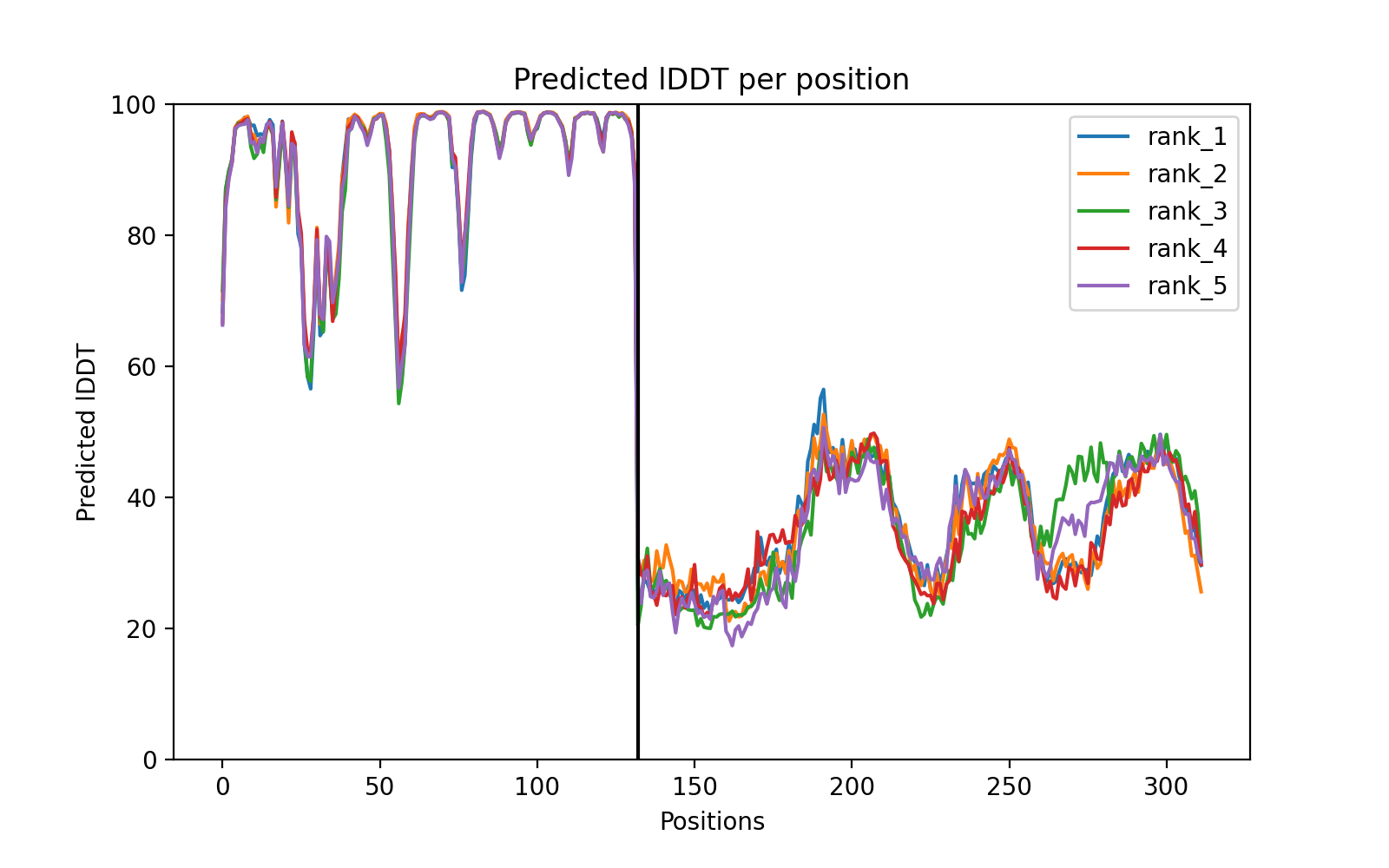

### FABP7_STX5_coverage.png

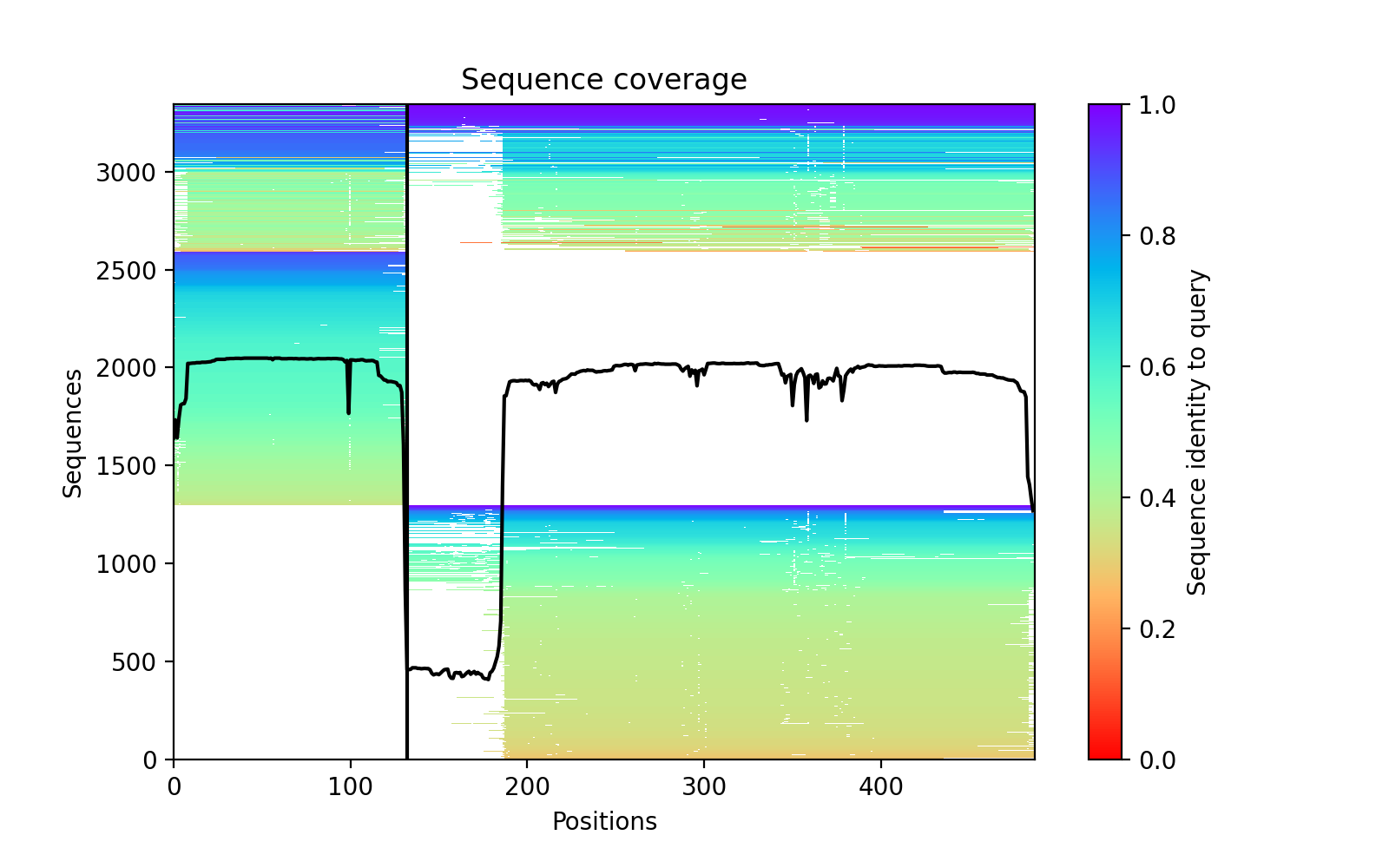

### FIGF_ZBTB25_PAE.png

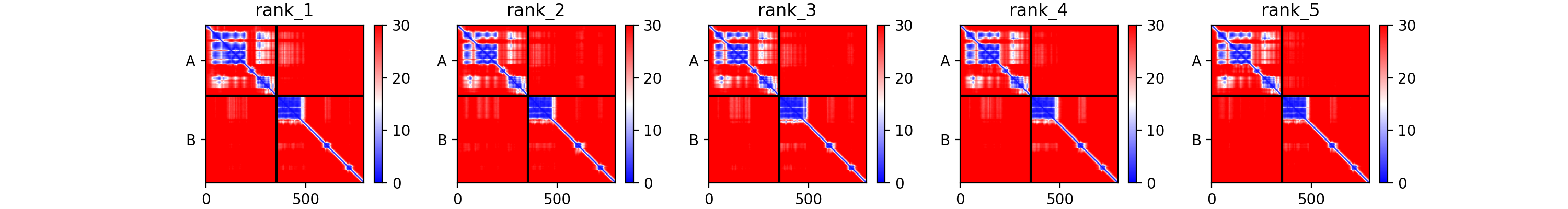

### GALK1_MCCC1_plddt.png

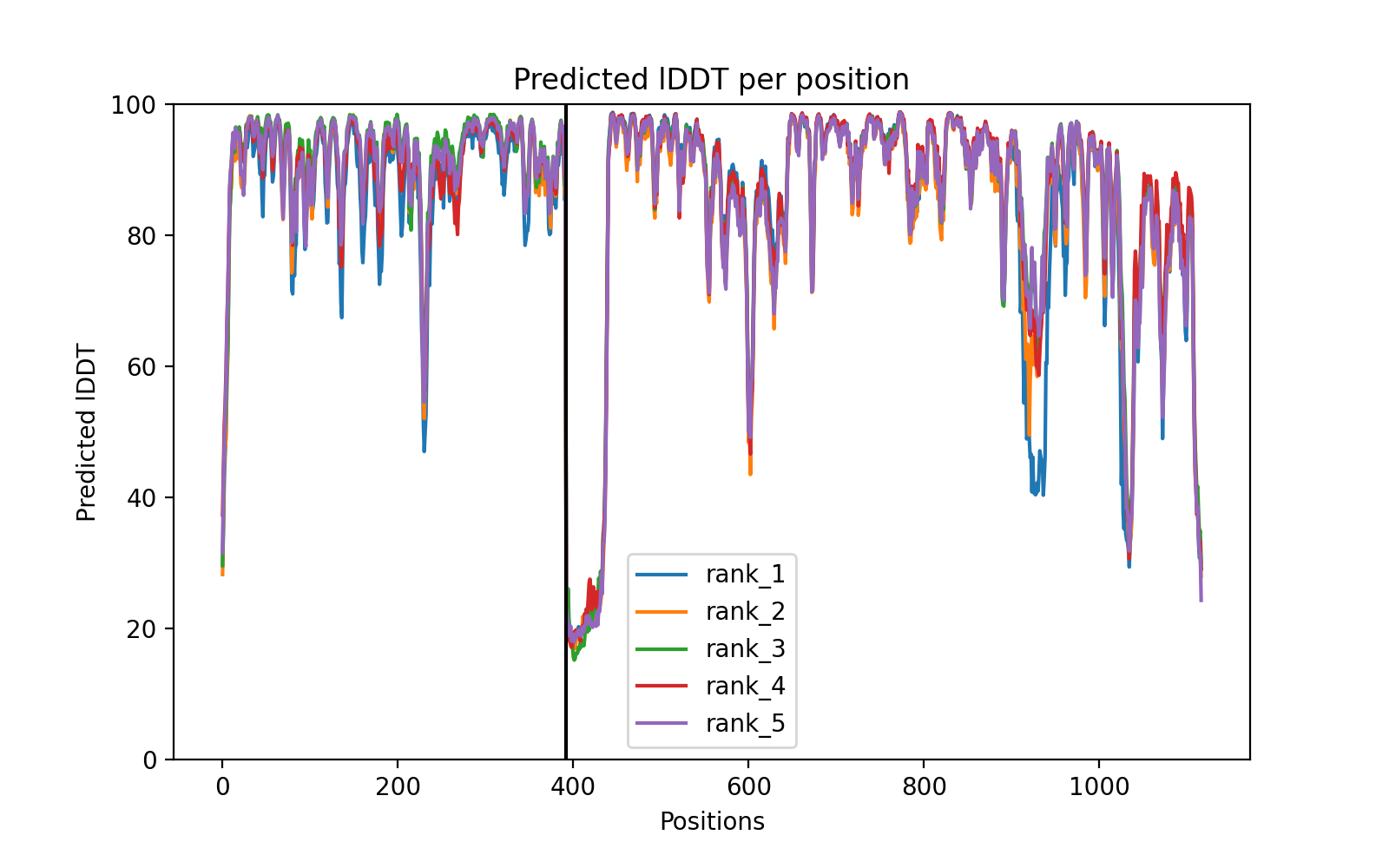

### GPD2_C22orf29_coverage.png

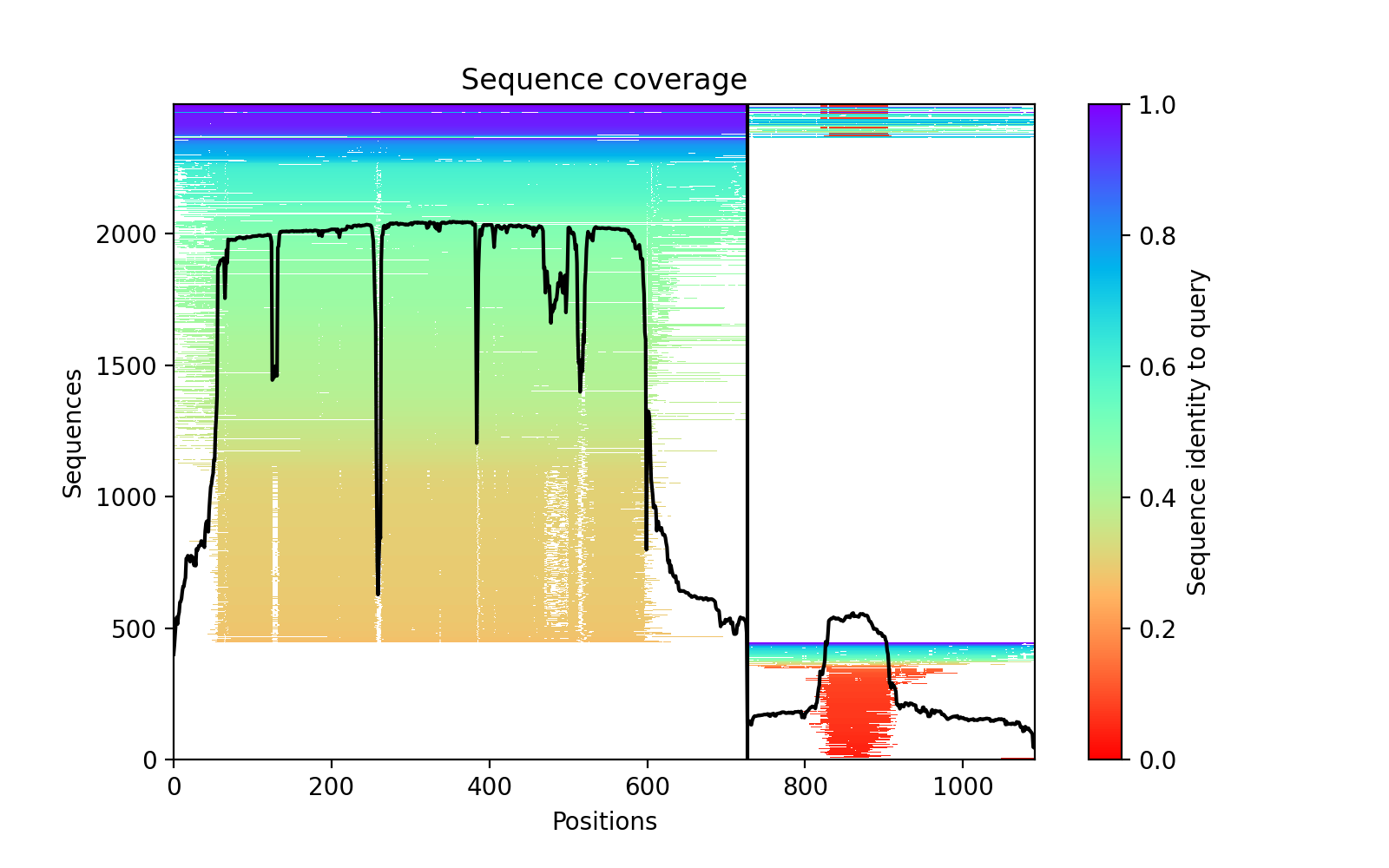

### GRB2_VAV1_coverage.png

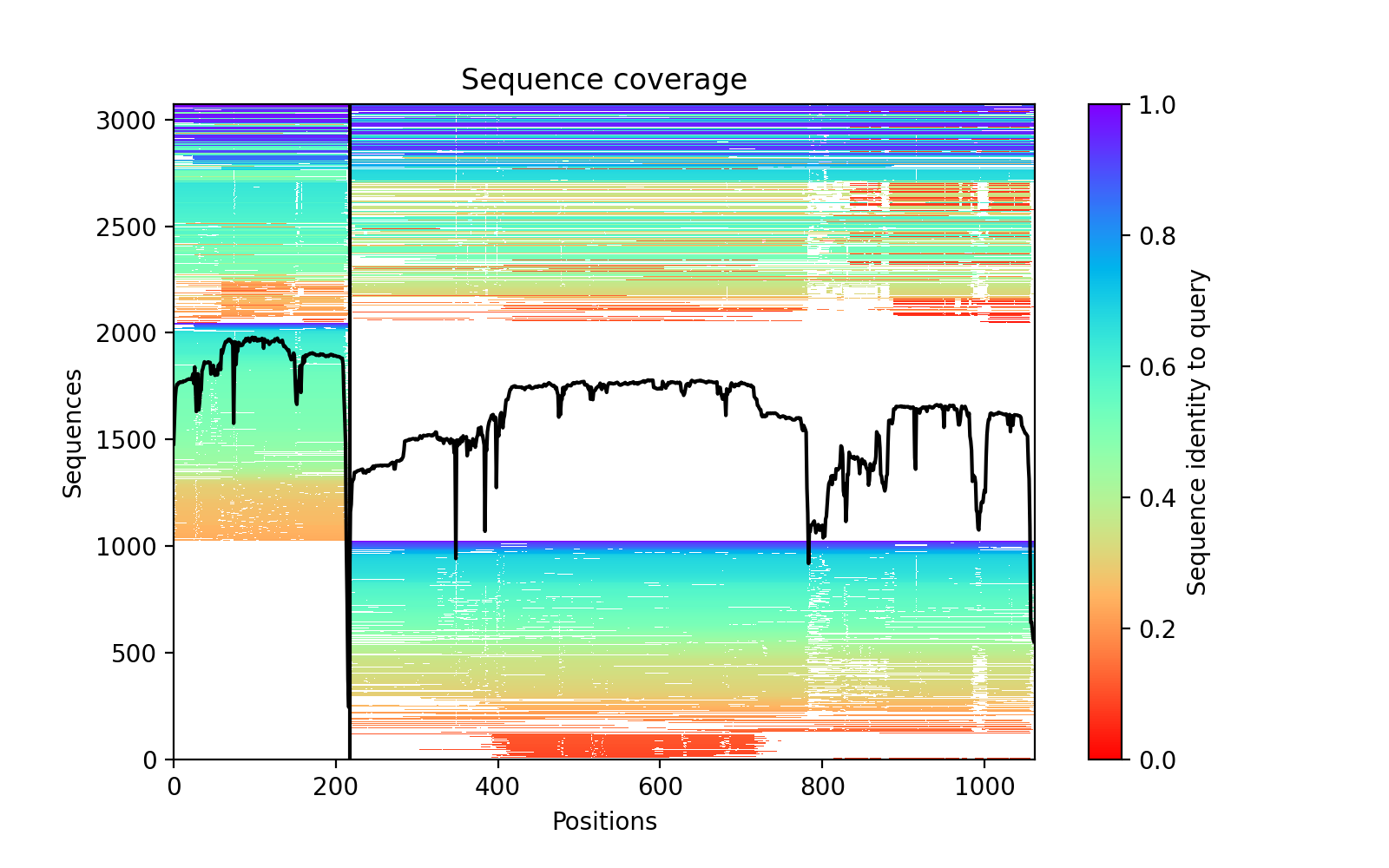

### HBA2_HBB_PAE.png

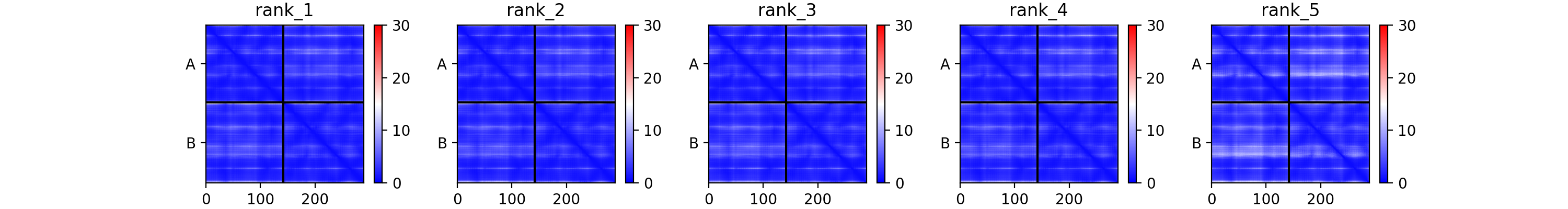

### HIST1H1C_NPDC1_PAE.png

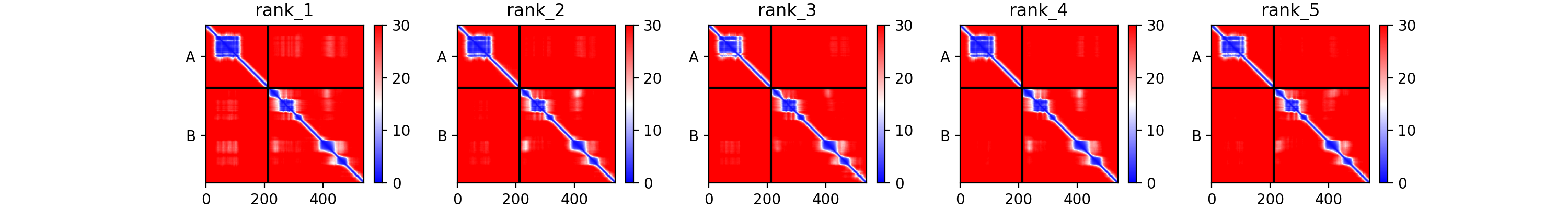

### IFIT1_EIF3E_PAE.png

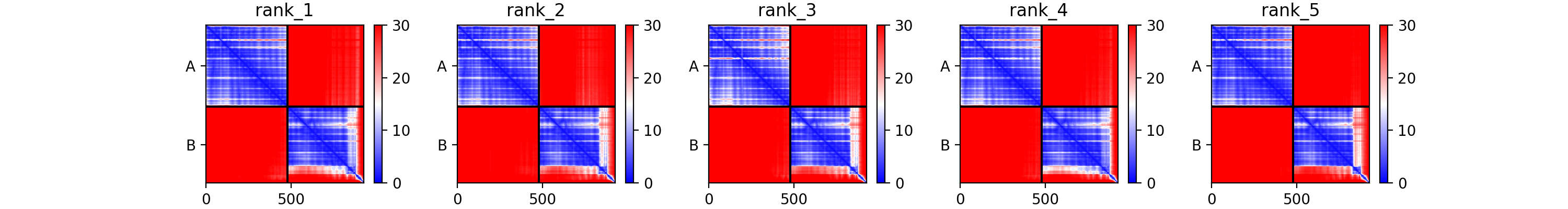

### IFIT1_EIF3E_plddt.png

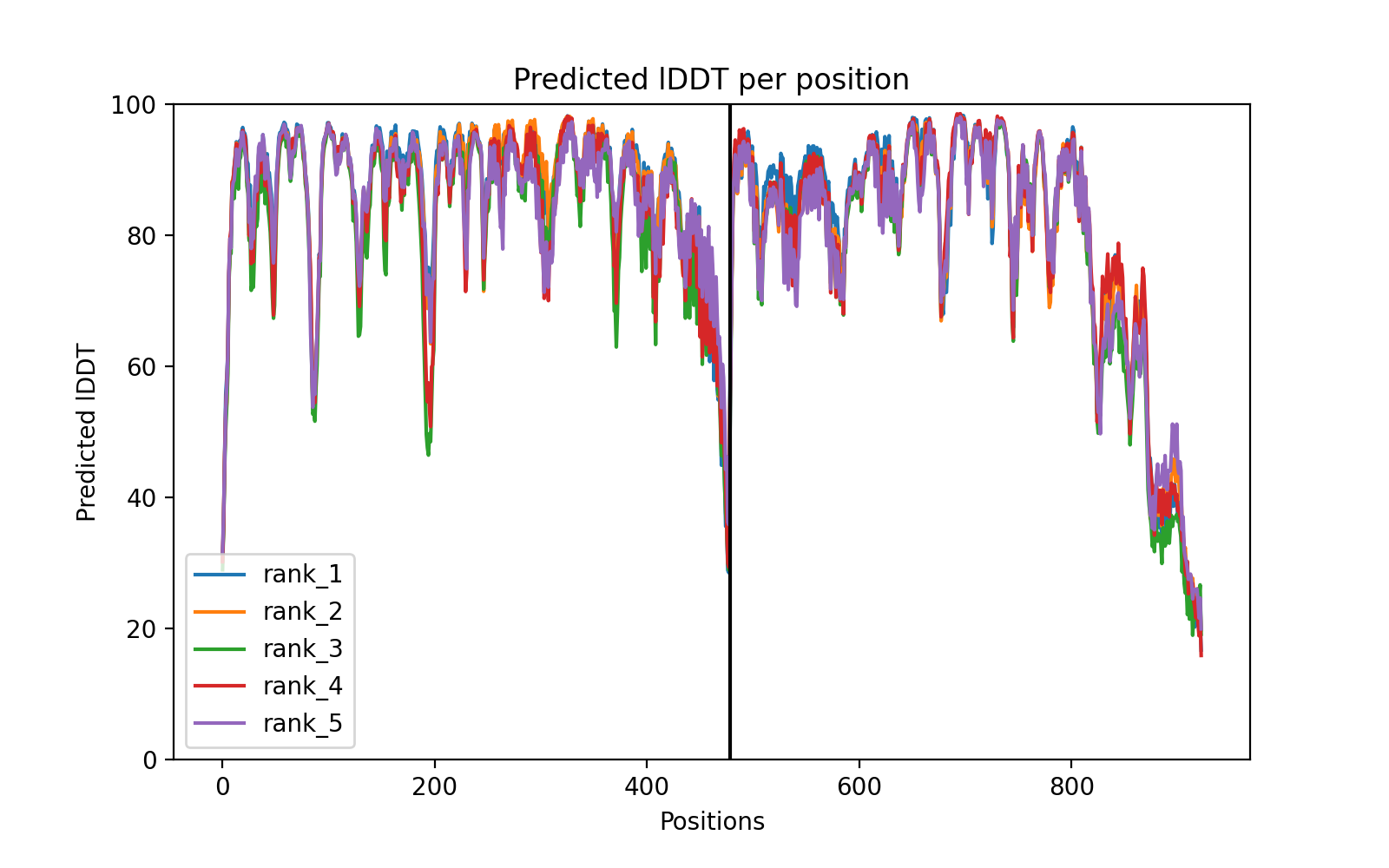

### IGF2_IGFBP4_PAE.png

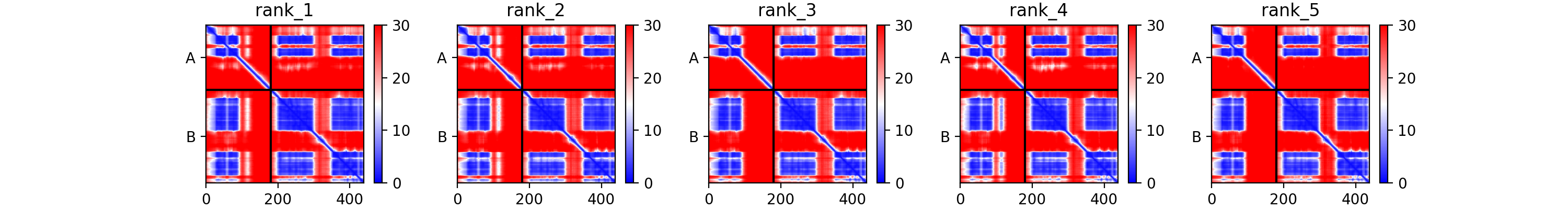

### ITPK1_TMEM22_coverage.png

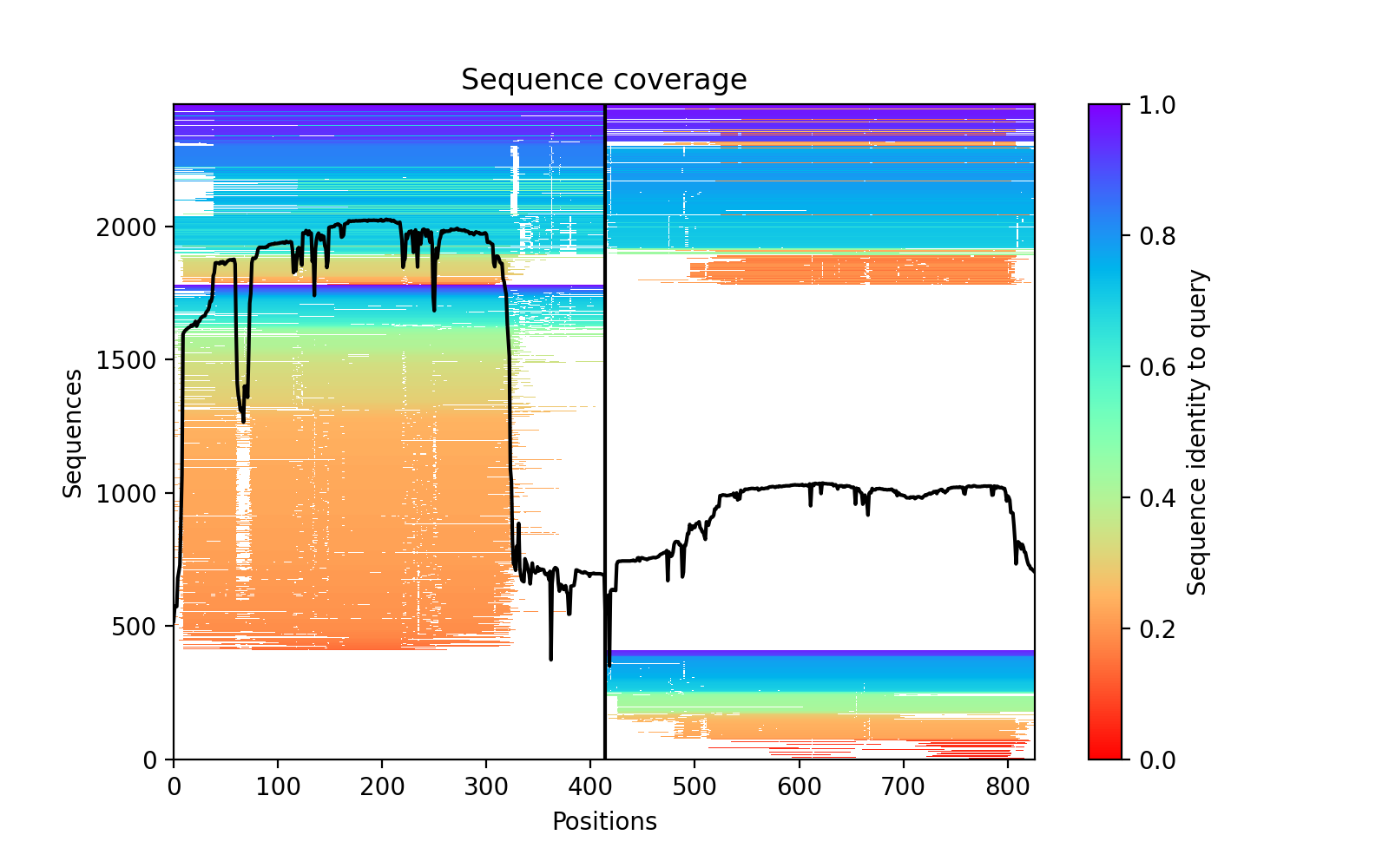

### ITPK1_TMEM22_plddt.png

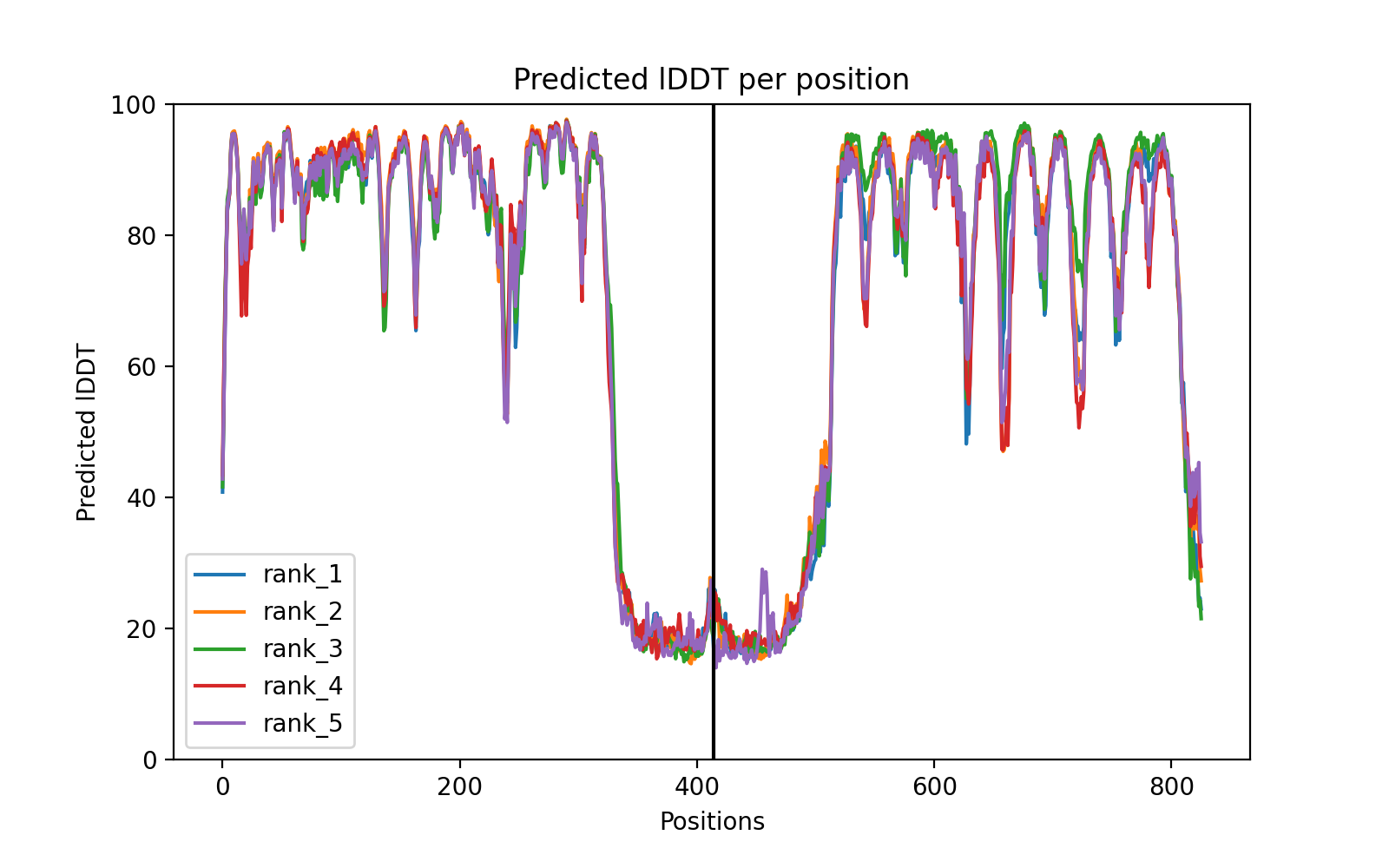

### LAMP2_UBE2G2_PAE.png

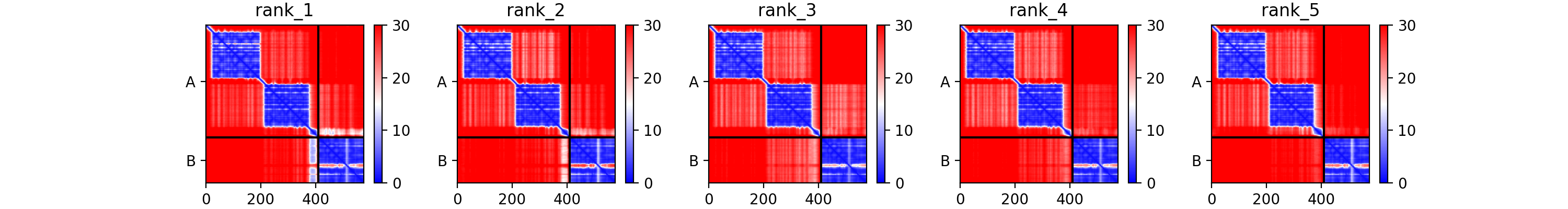

### LCP2_GRAP2_coverage.png

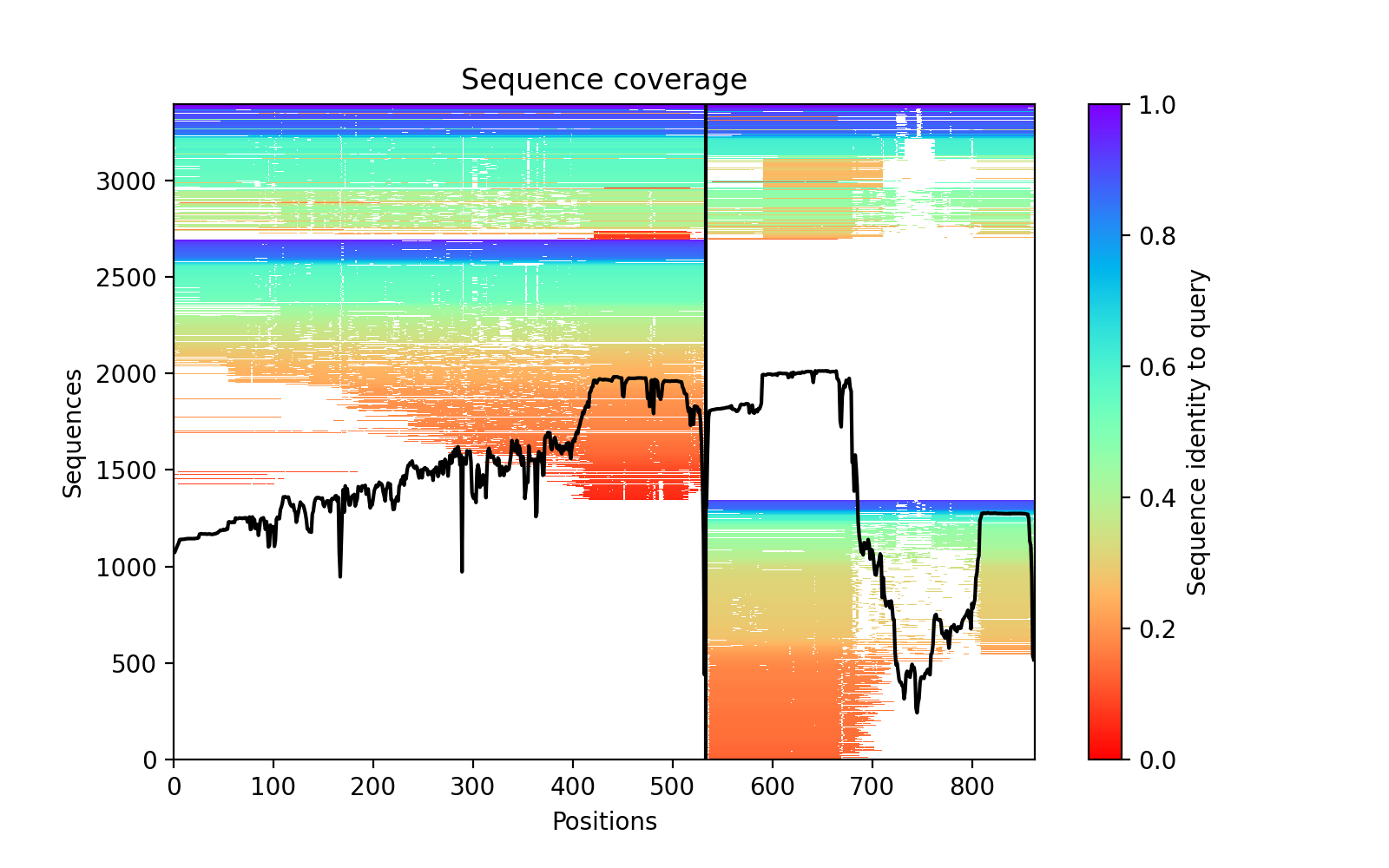

### LCP2_GRAP2_plddt.png

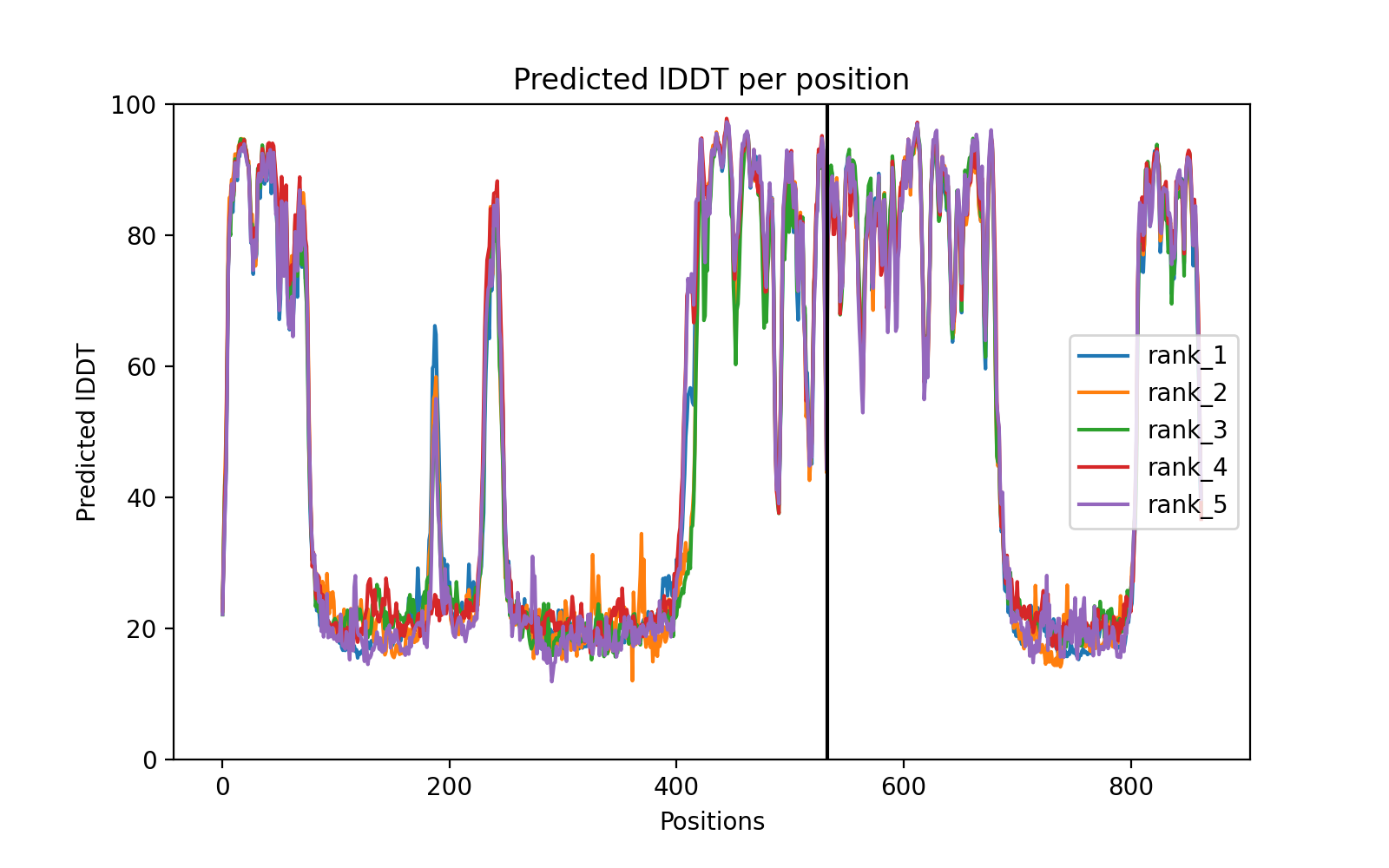

### LCP2_NCK1_coverage.png

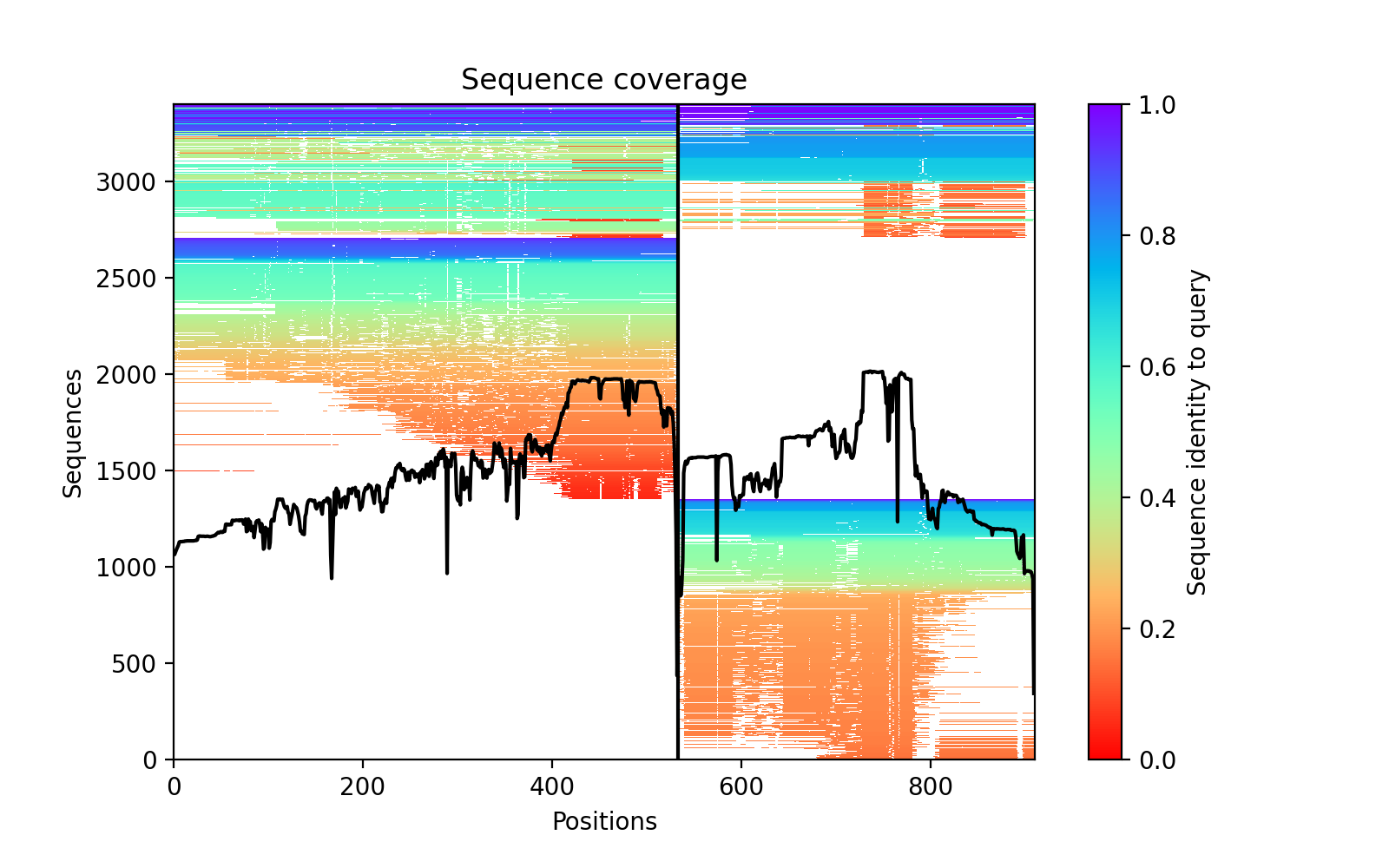

### LCP2_NCK1_PAE.png

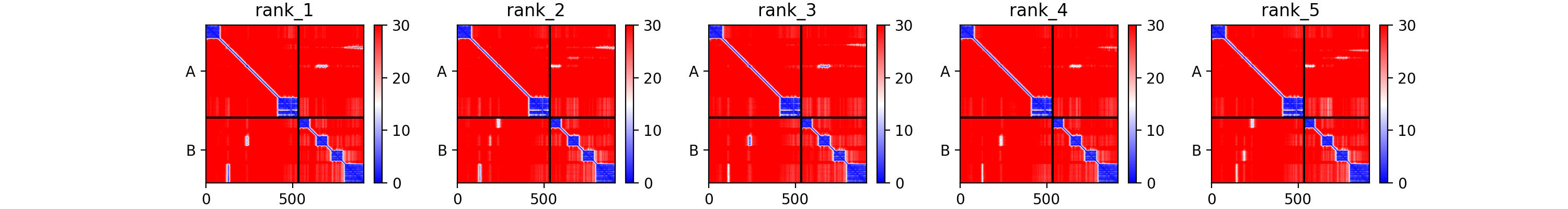

### LMNA_LMNB1_plddt.png

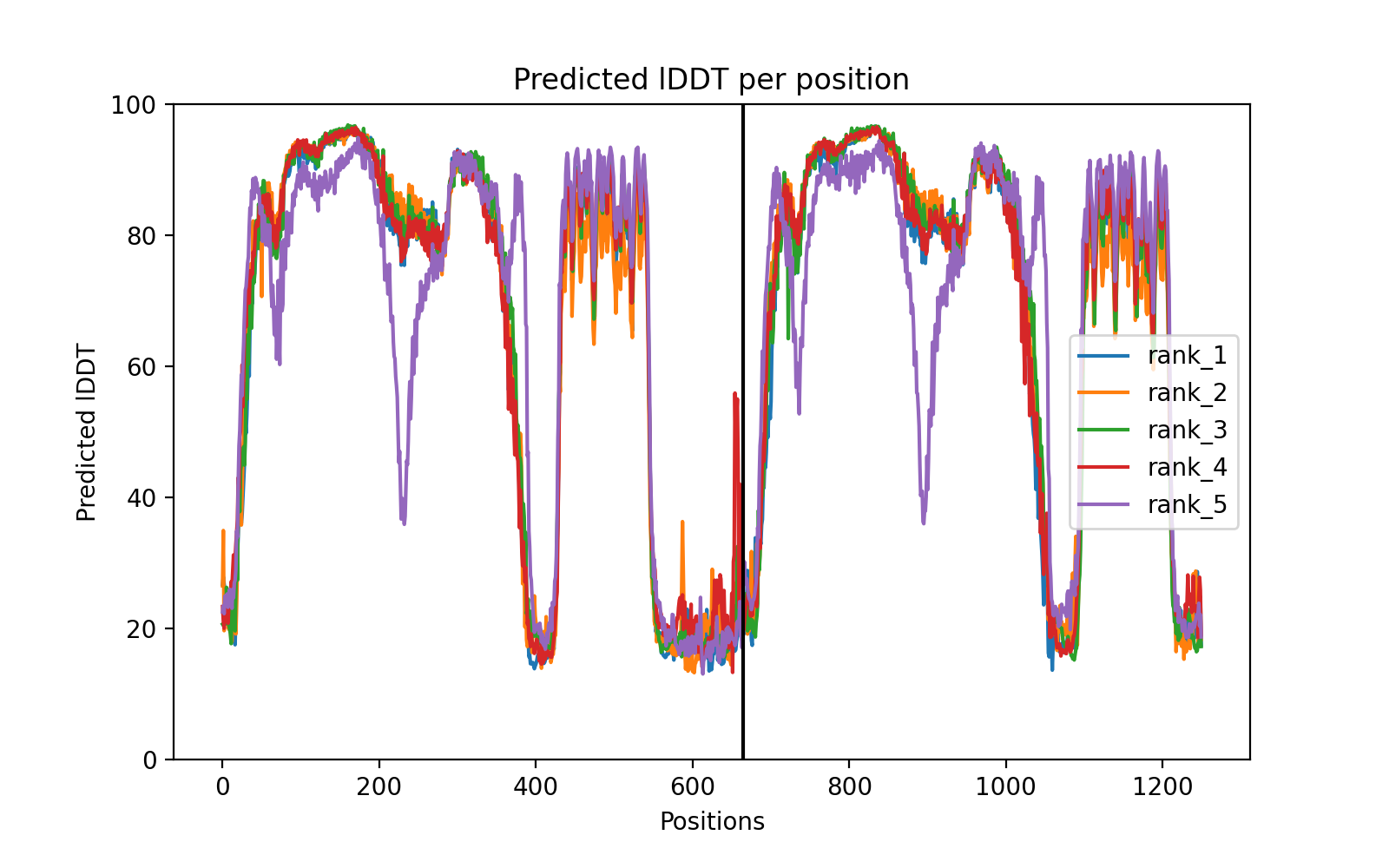

### LSM3_LSM2_plddt.png

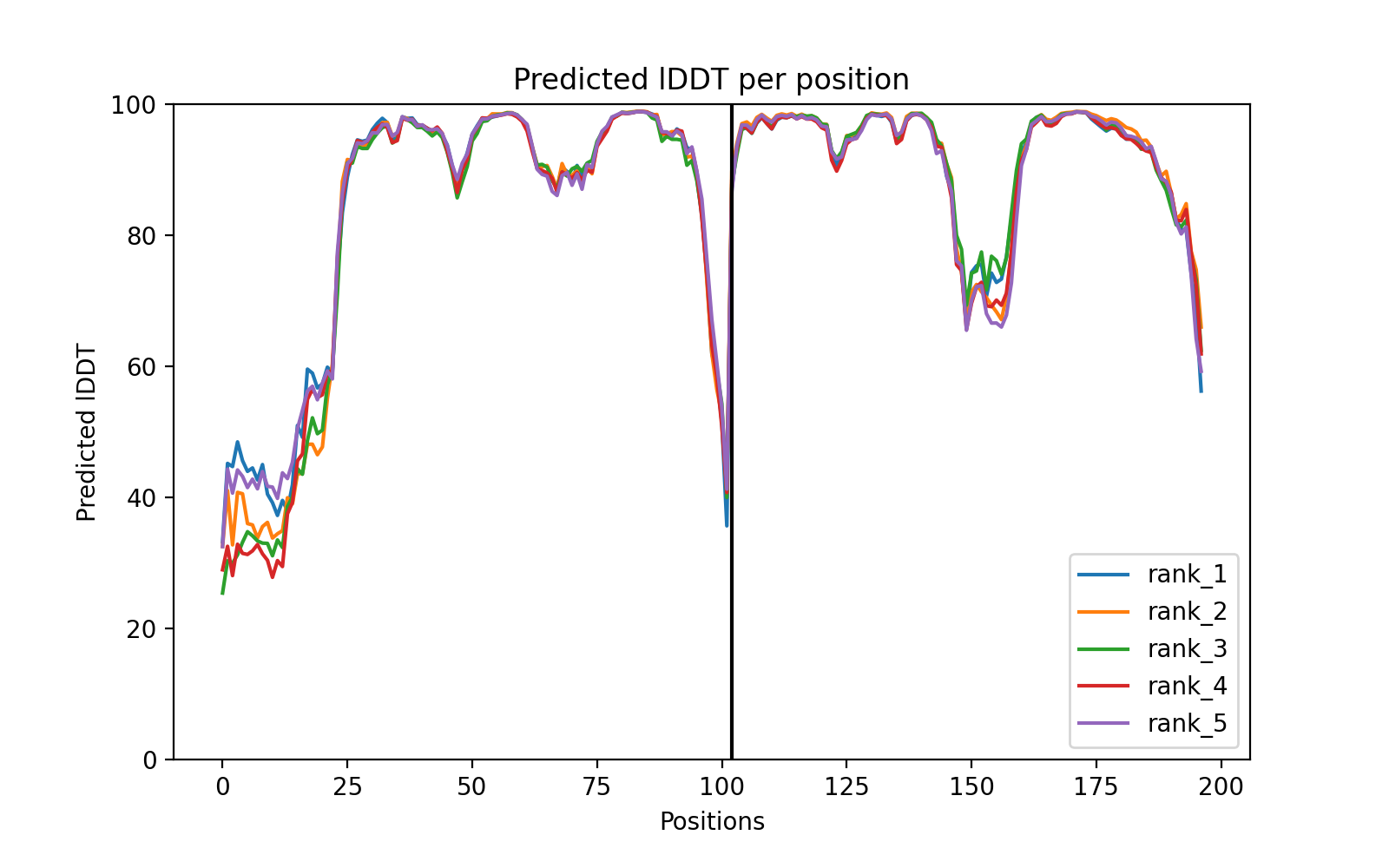

### MAFG_NFE2L1_PAE.png

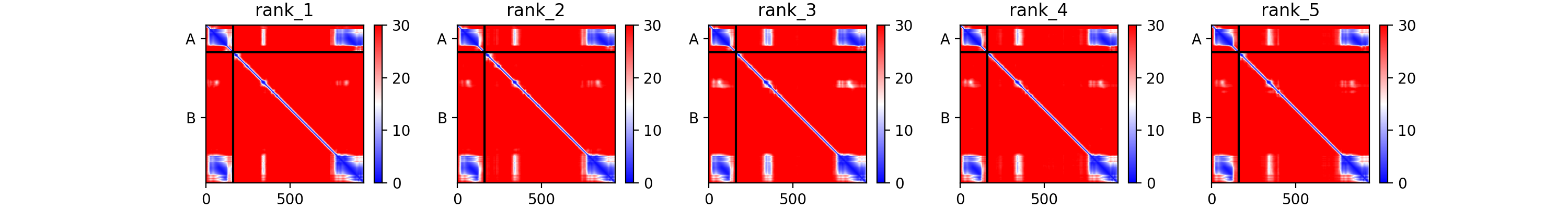

### MAFG_NFE2L1_plddt.png

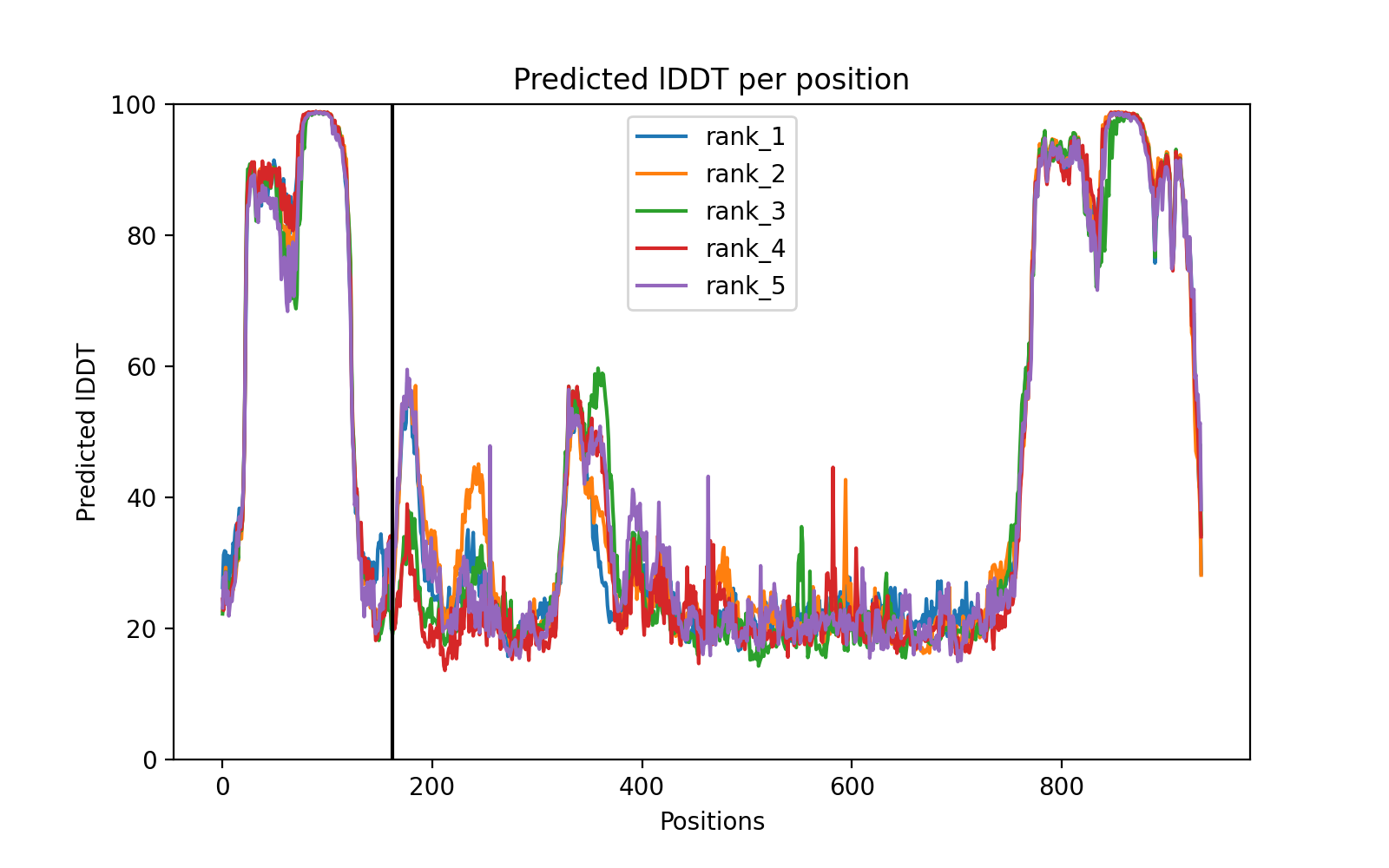

### MNAT1_GMPPA_PAE.png

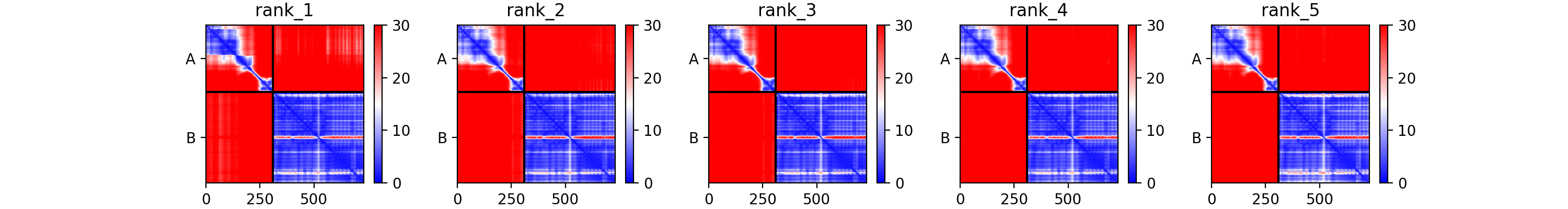

### MOBP_MRPS25_coverage.png

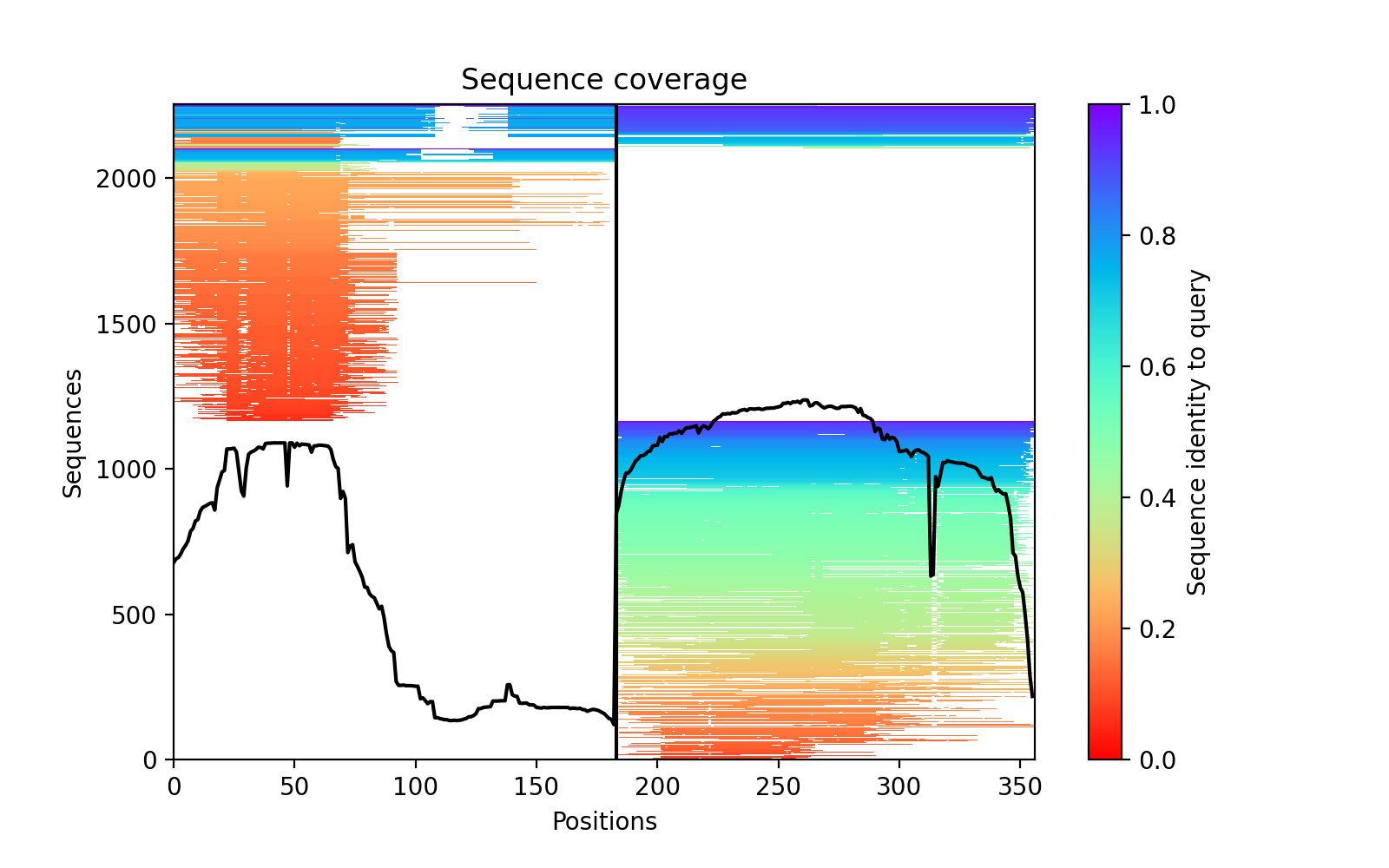

### NAT2_DNAJA1_PAE.png

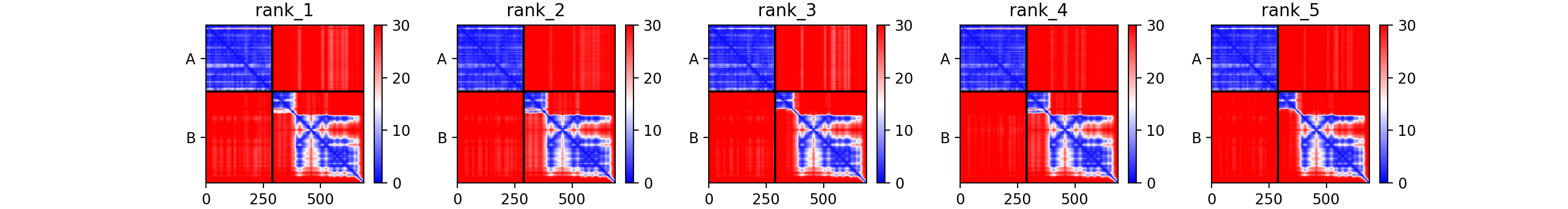

### NAT2_DNAJA1_plddt.png

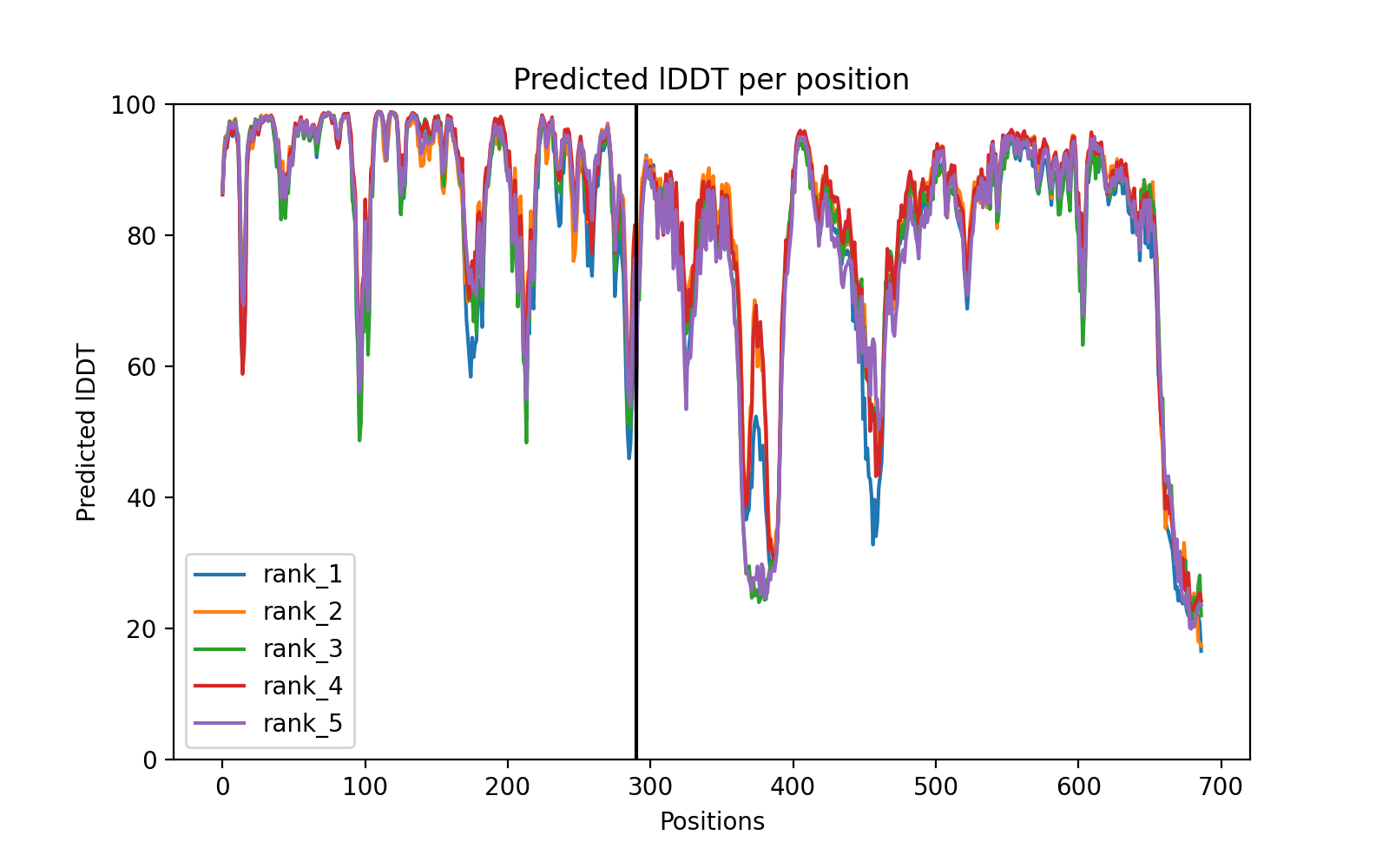

### NDP_NUDT4_PAE.png

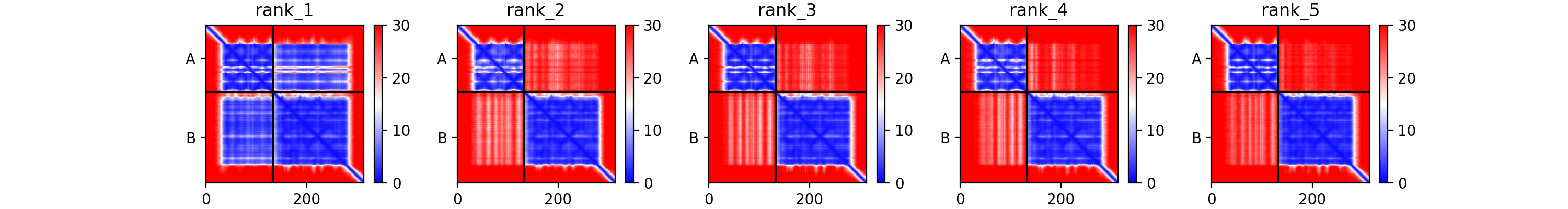

### NKX2-5_CSGALNACT2_coverage.png

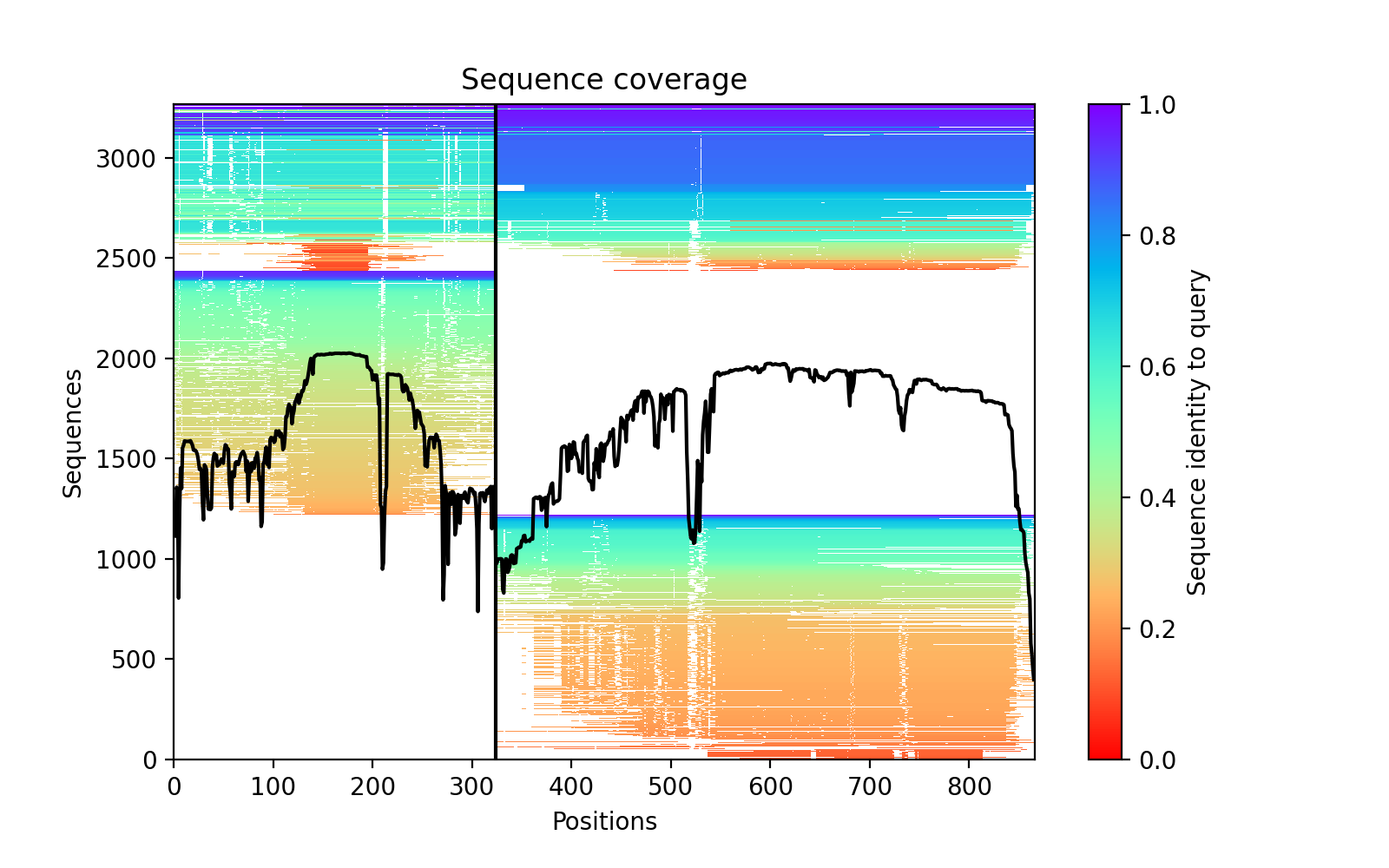

### NSP4_NSP4_8f98f_coverage.png

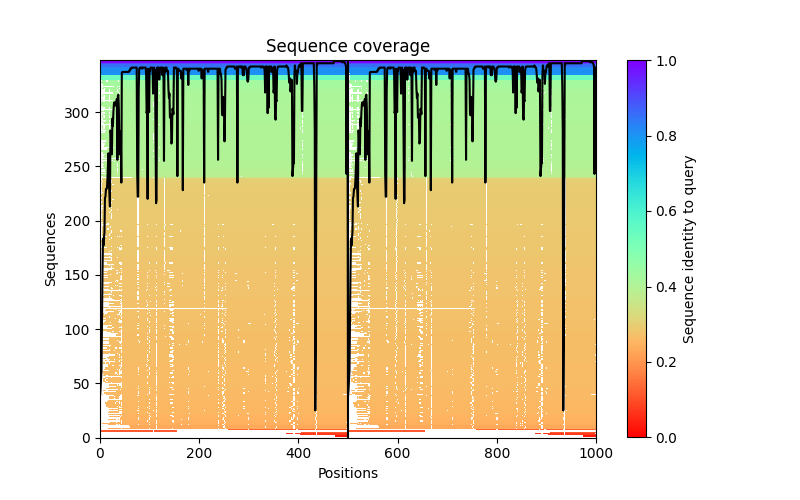

### NSP4_NSP4_8f98f_PAE.png

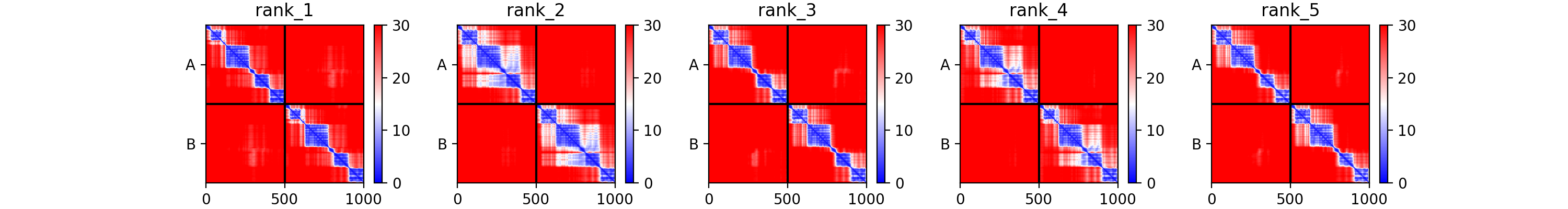

### NSP4_NSP4_8f98f_plddt.png

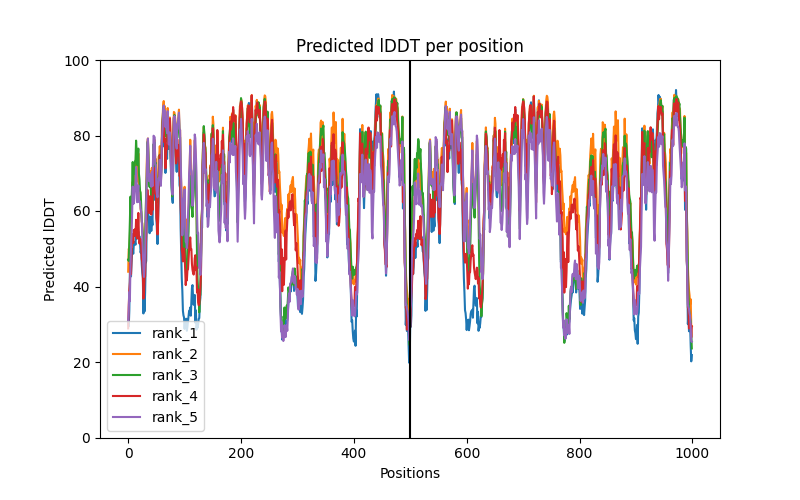

### NSP6_ORF7a_43e6e_coverage.png

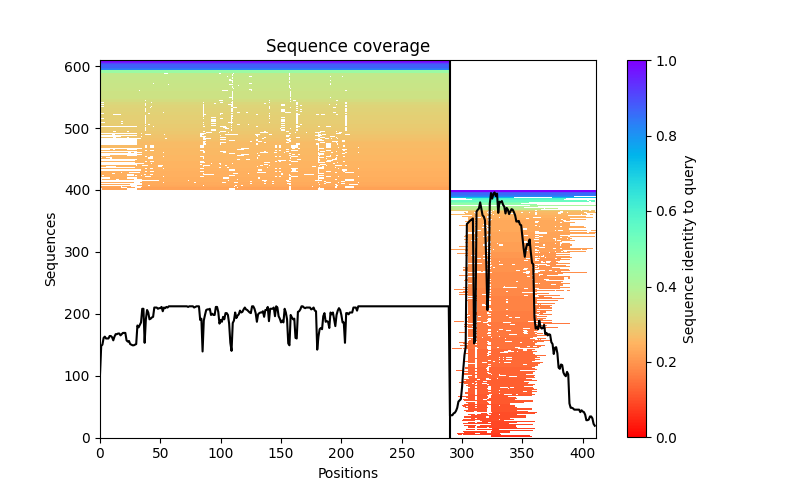

### NSP6_ORF7a_43e6e_PAE.png

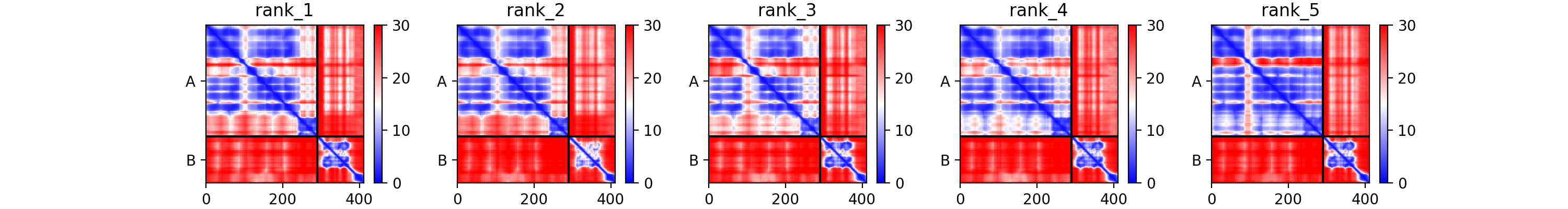

### NSP6_ORF7a_43e6e_plddt.png

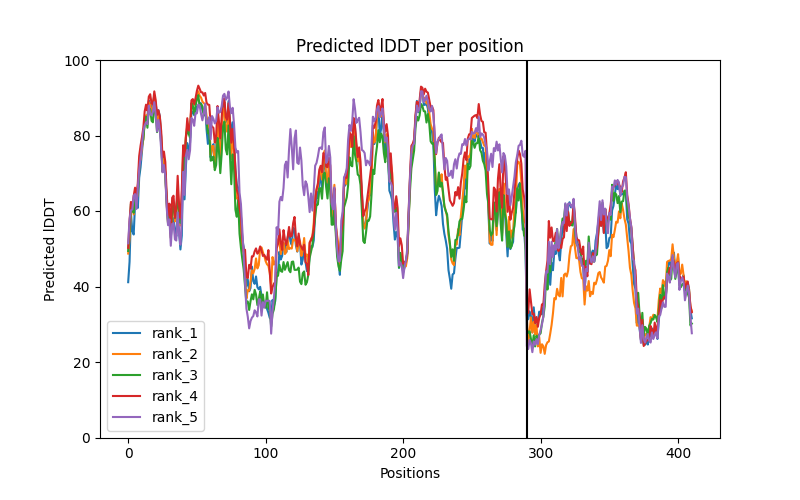

### NSP8_NSP12_db995_coverage.png

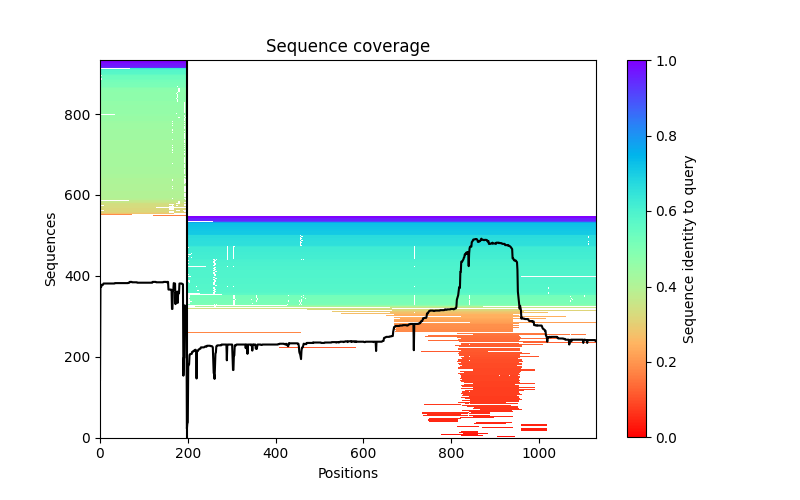

### NSP8_NSP12_db995_PAE.png

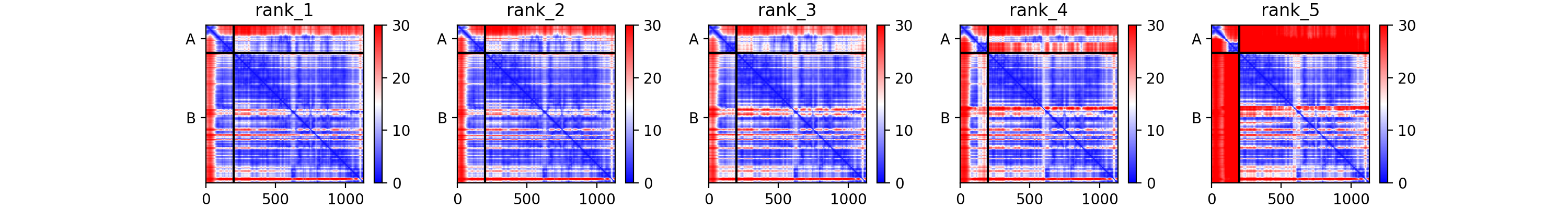

### NSP8_NSP12_db995_plddt.png

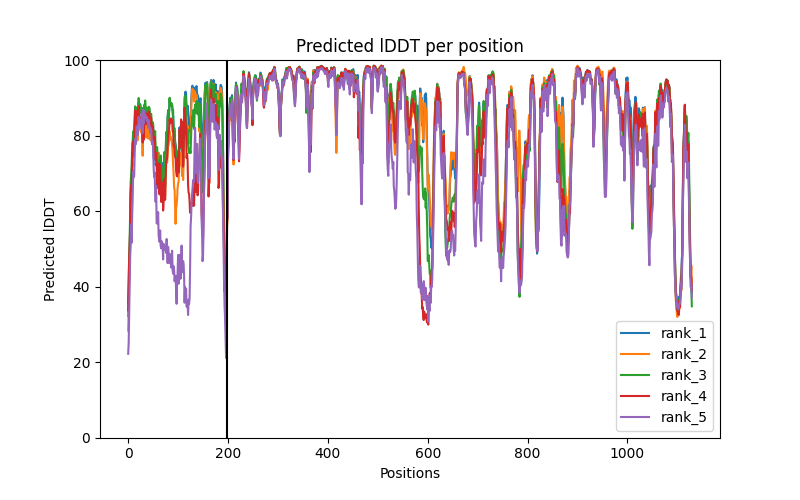

### NSP12_NSP16_40e1a_coverage.png

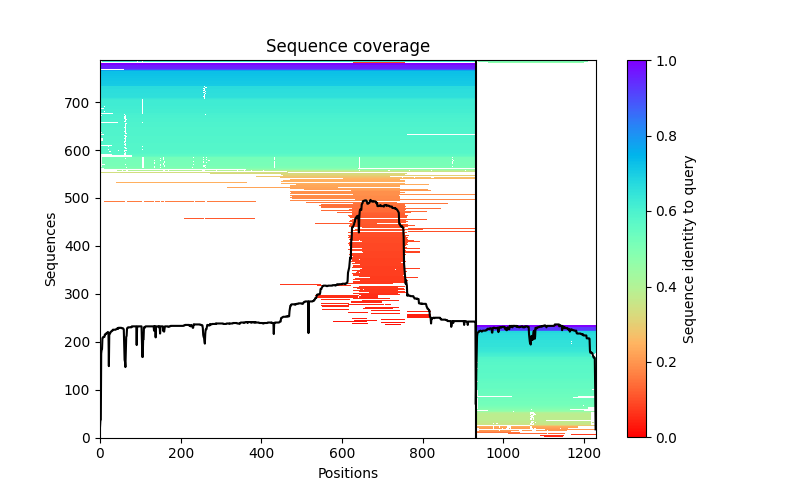

### NSP12_NSP16_40e1a_PAE.png

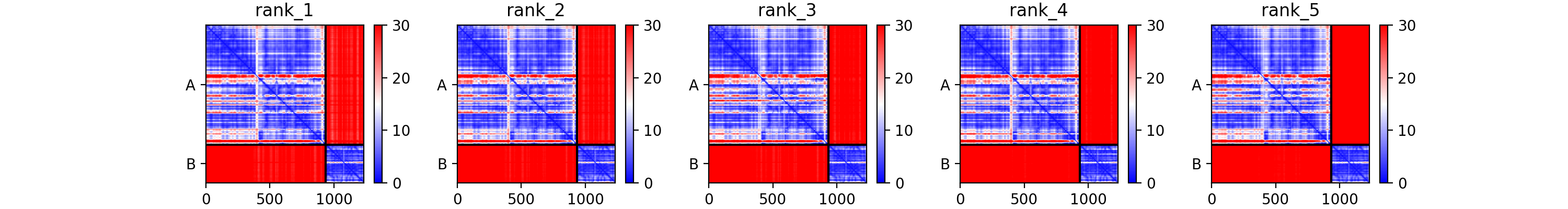

### NSP12_NSP16_40e1a_plddt.png

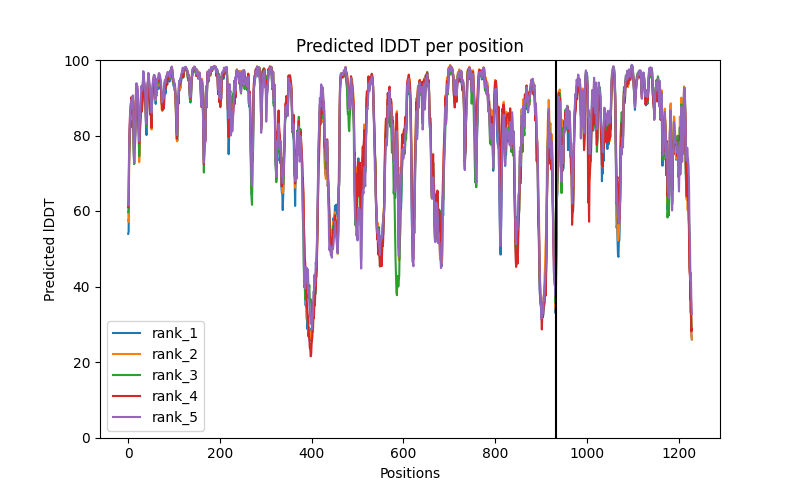

### NUDT2_MIIP_plddt.png

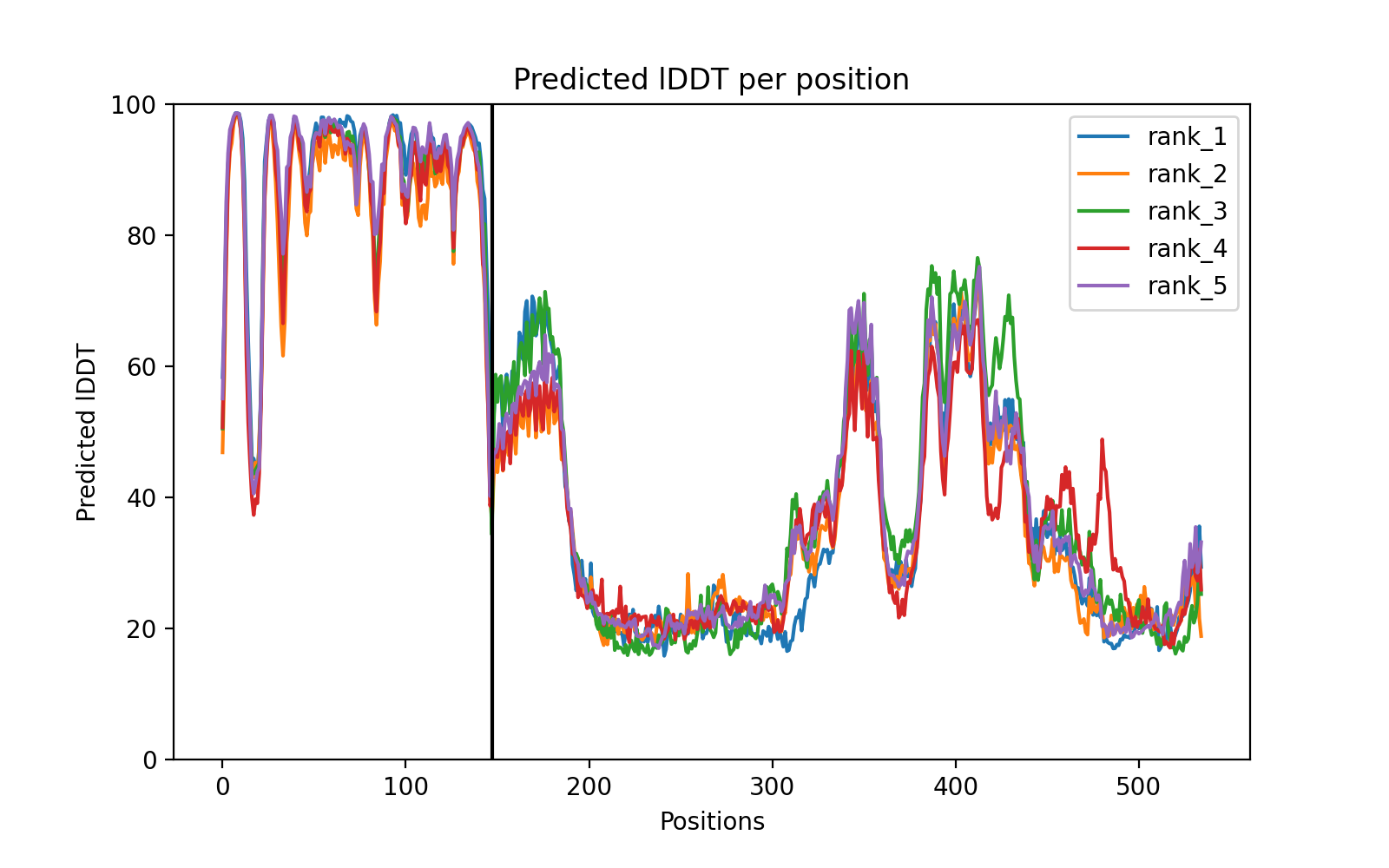

### ORF3a_ORF3a_4d856_coverage.png

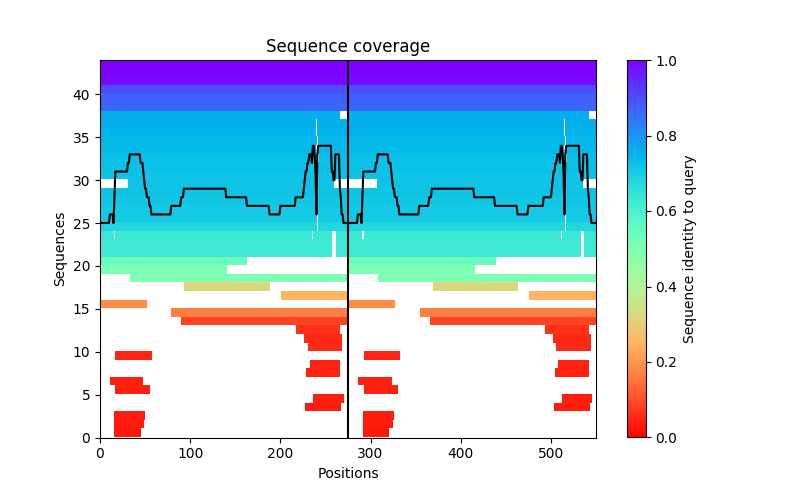

### ORF3a_ORF3a_4d856_PAE.png

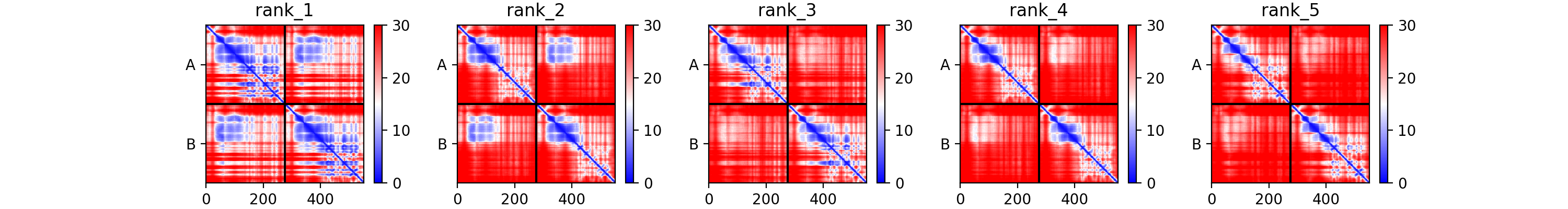

### ORF3a_ORF3a_4d856_plddt.png

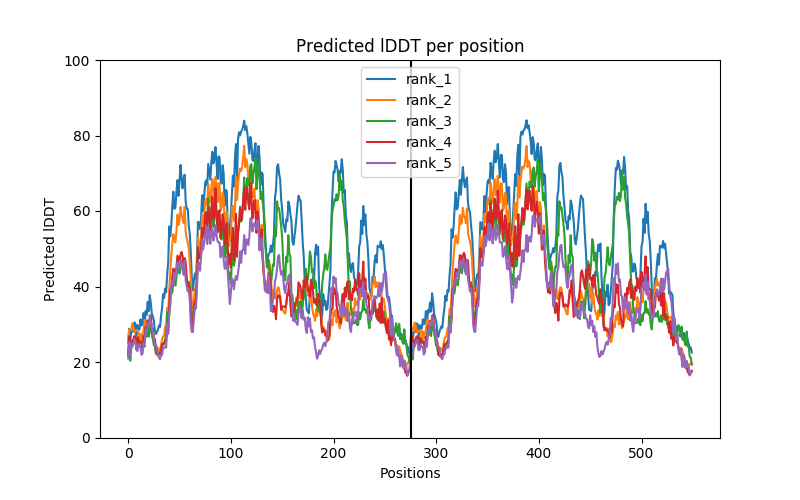

### ORF7a_ORF3a_5fed4_coverage.png

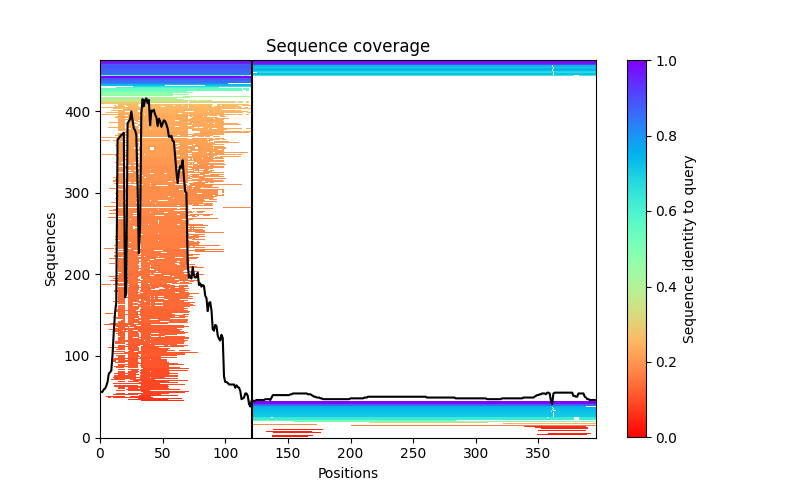

### ORF7a_ORF3a_5fed4_PAE.png

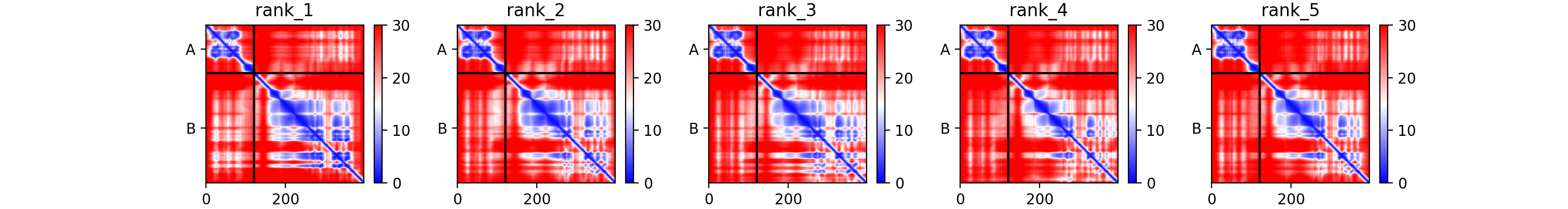

### ORF7a_ORF3a_5fed4_plddt.png

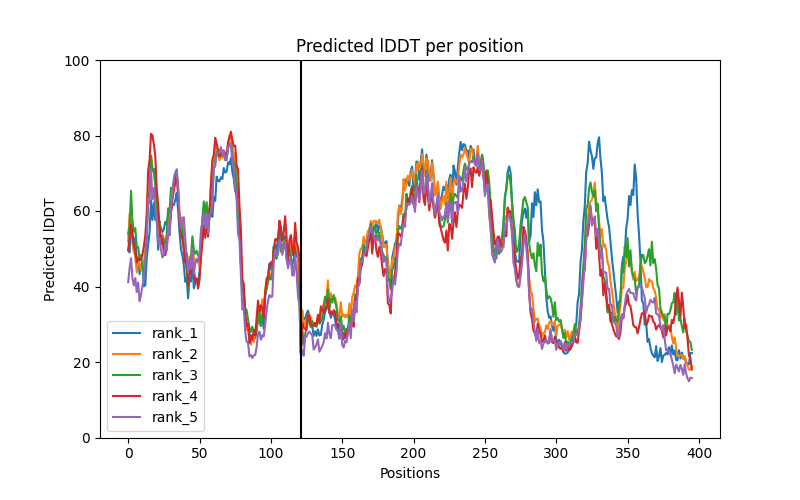

### ORF7a_ORF7a_b4818_coverage.png

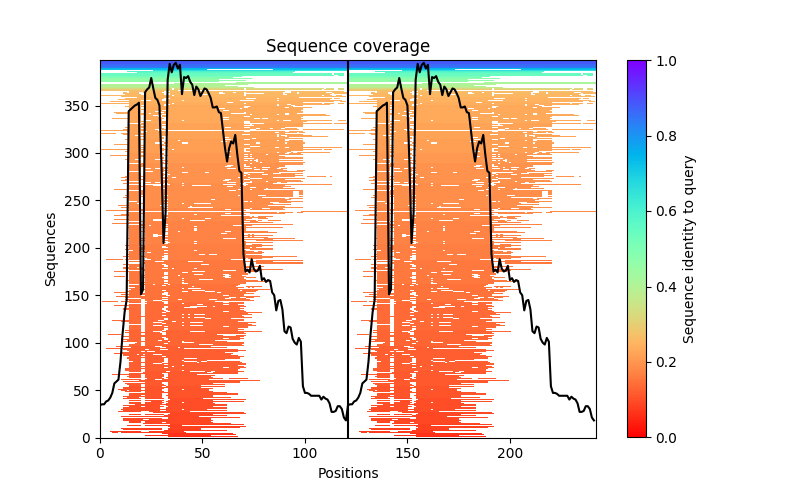

### ORF7a_ORF7a_b4818_PAE.png

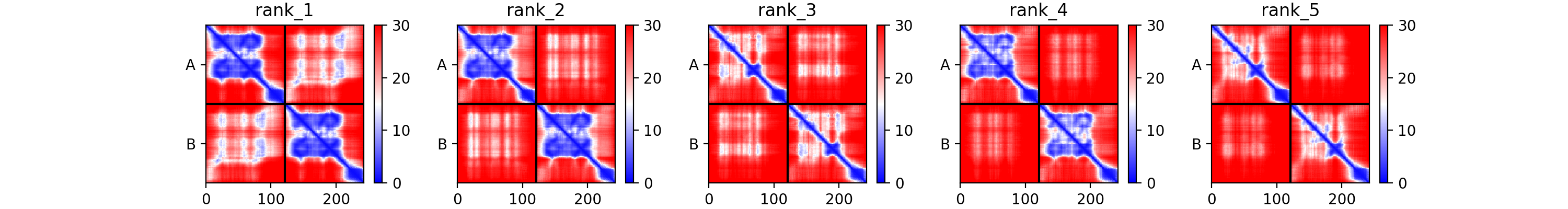

### ORF7a_ORF7a_b4818_plddt.png

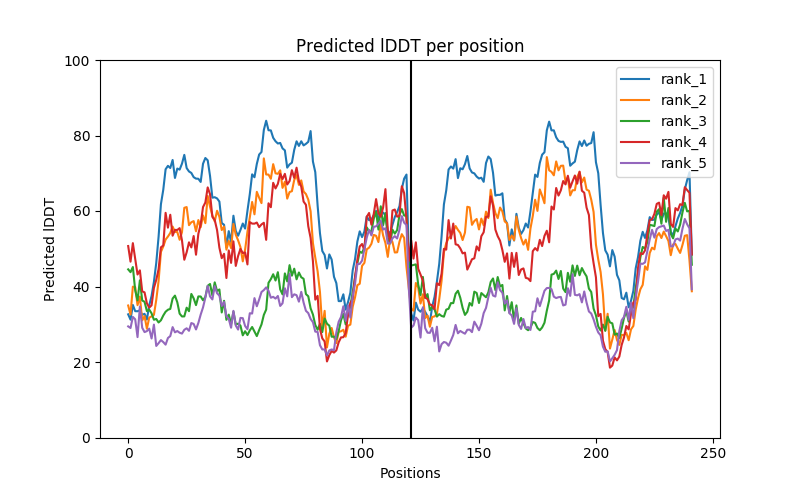

### ORF7b_M_904cd_coverage.png

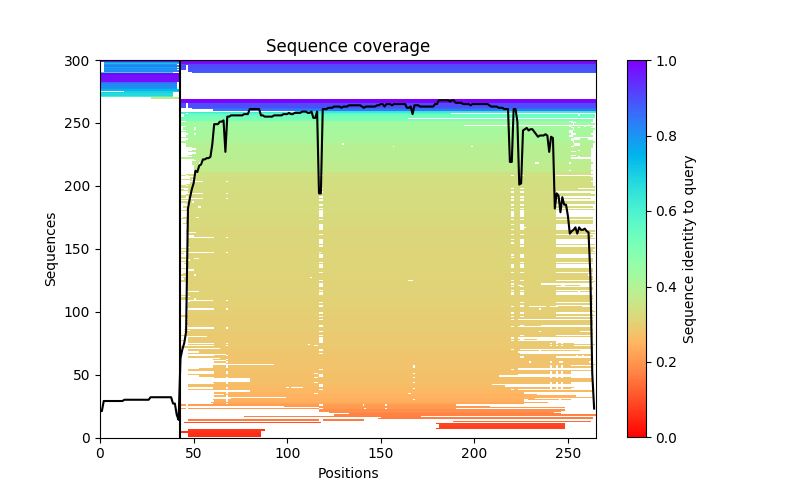

### ORF7b_M_904cd_PAE.png

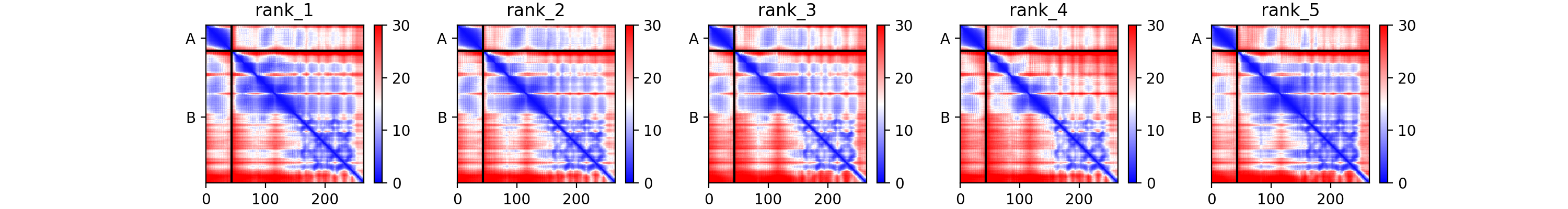

### ORF7b_M_904cd_plddt.png

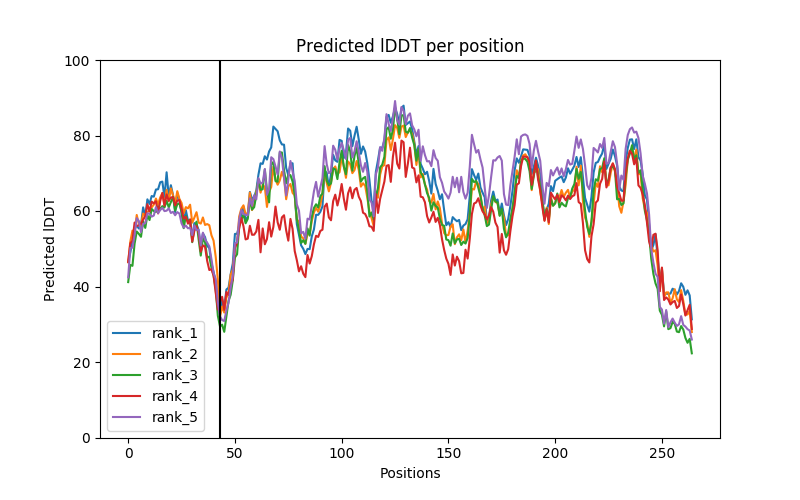

### OSM_ZNF688_coverage.png

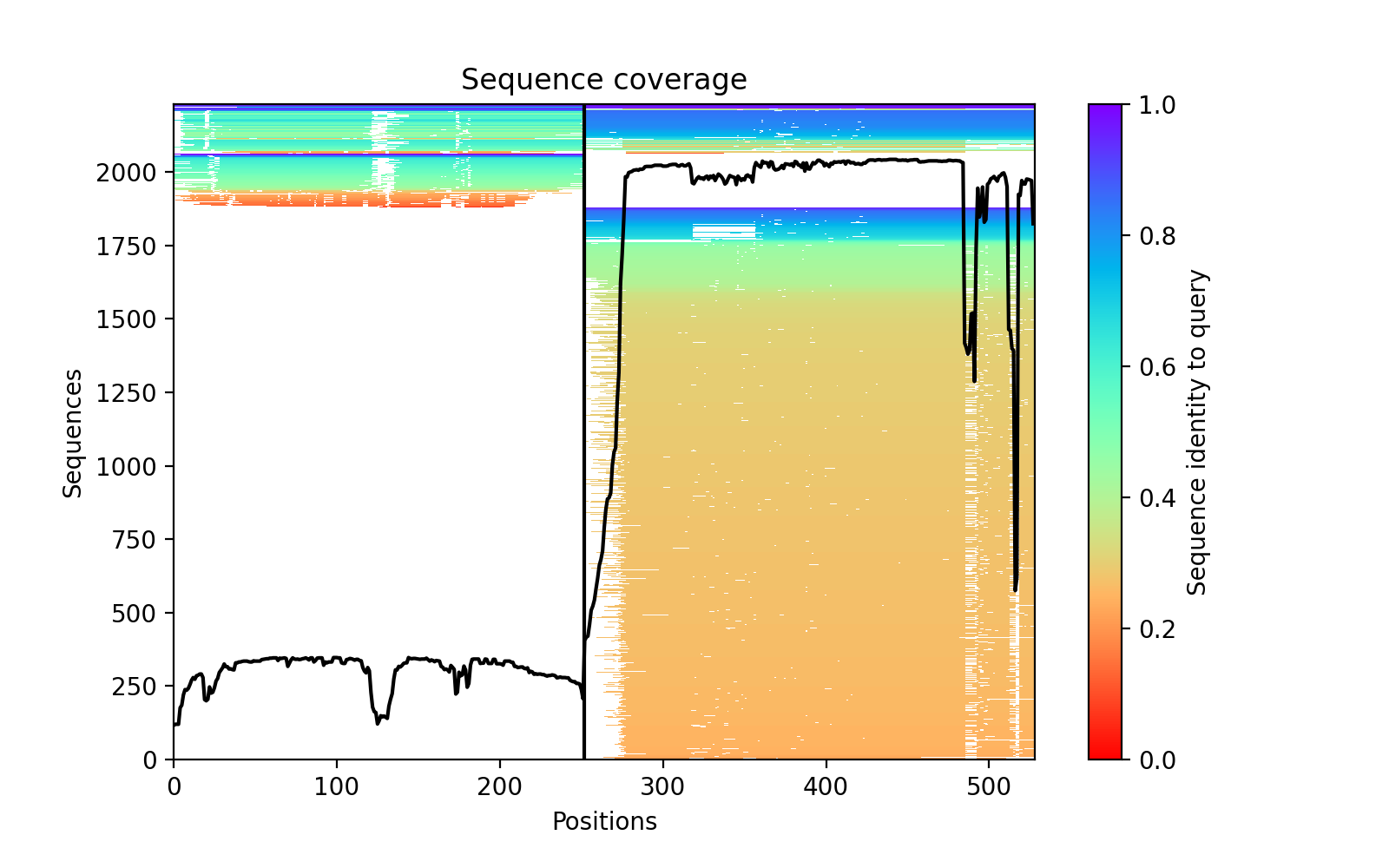

### PDGFRA_NDFIP1_coverage.png

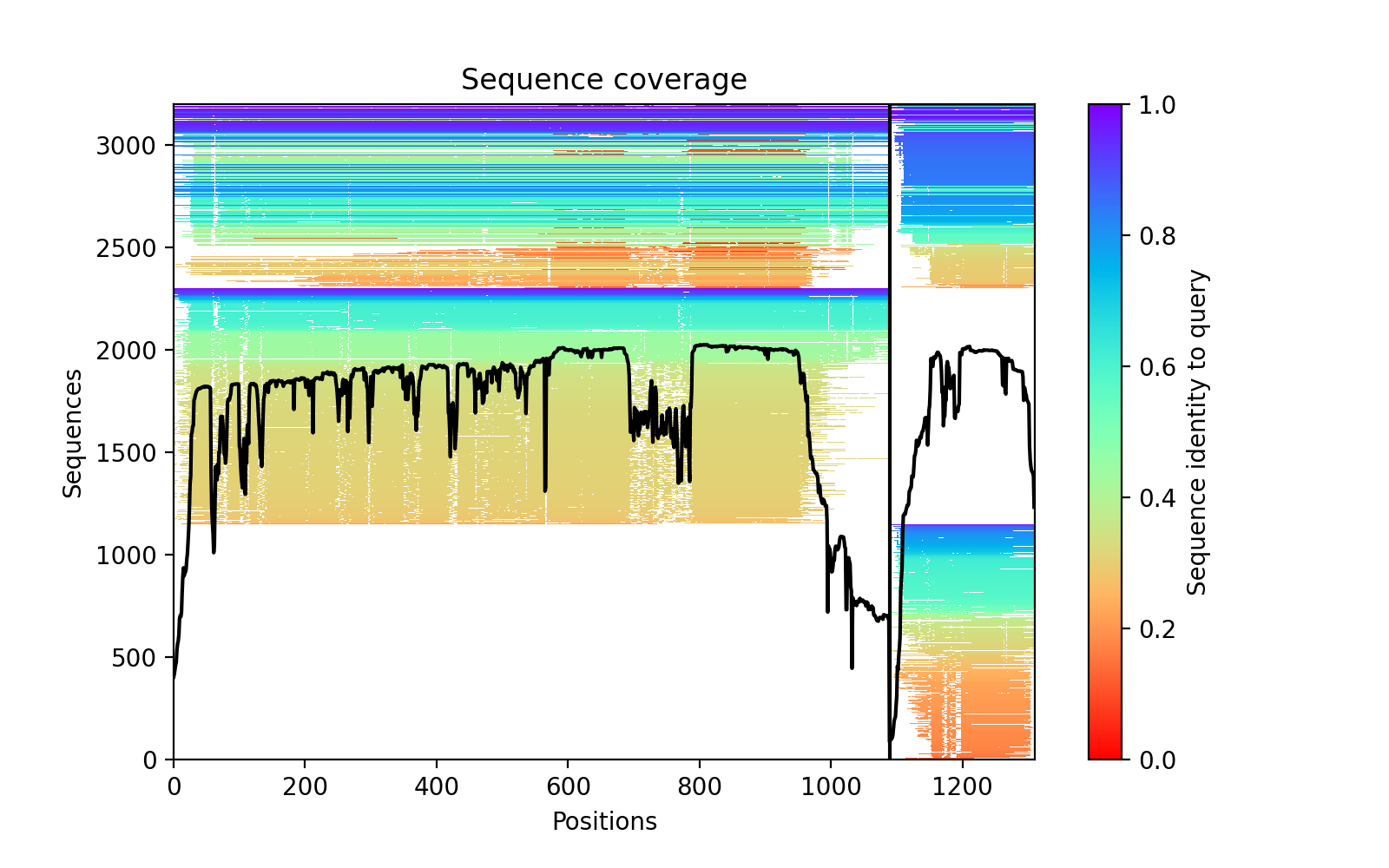

### PDGFRA_NDFIP1_PAE.png

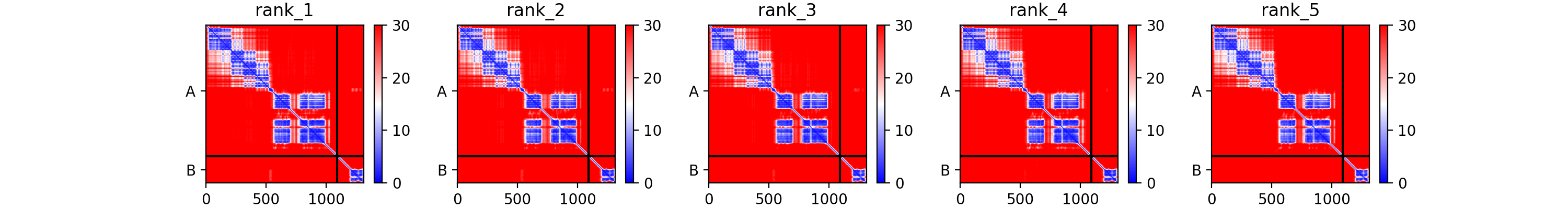

### PEX14_PEX19_PAE.png

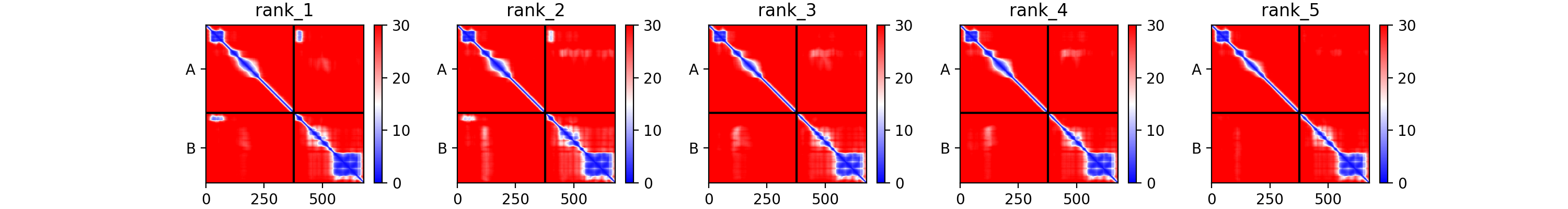

### PEX19_PEX16_plddt.png

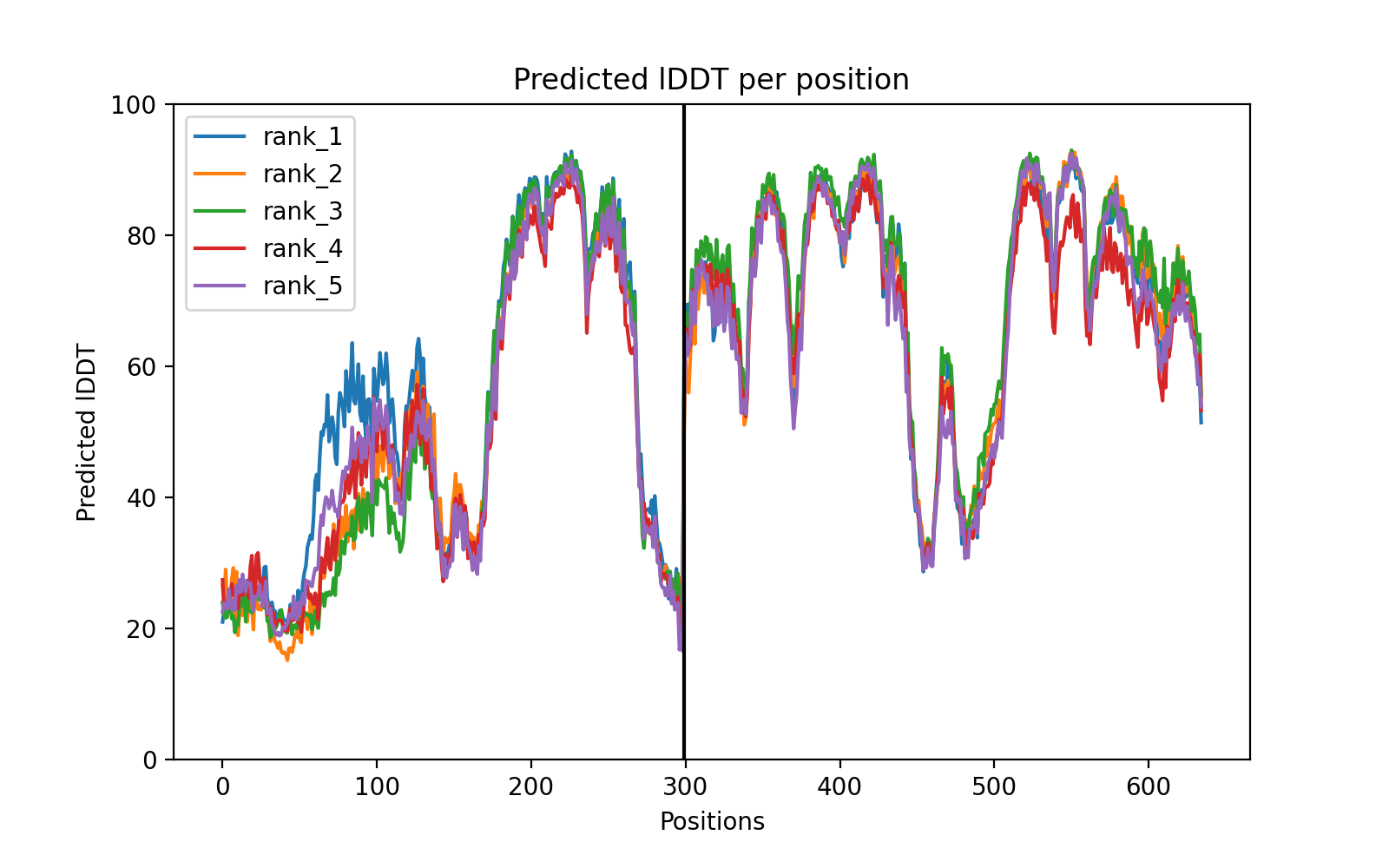

### PMCH_RIC3_plddt.png

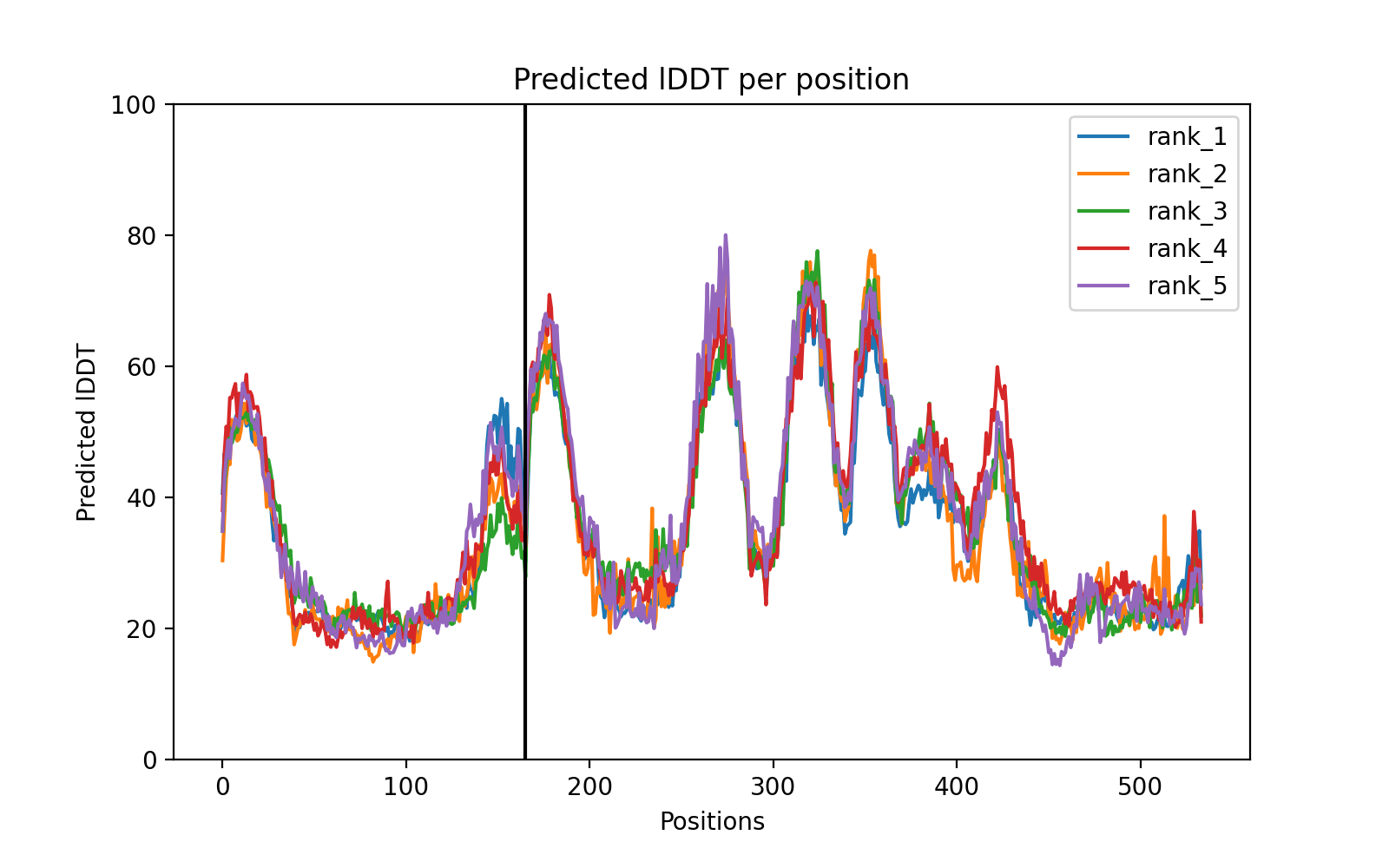

### PPP3CA_PPP3R1_plddt.png

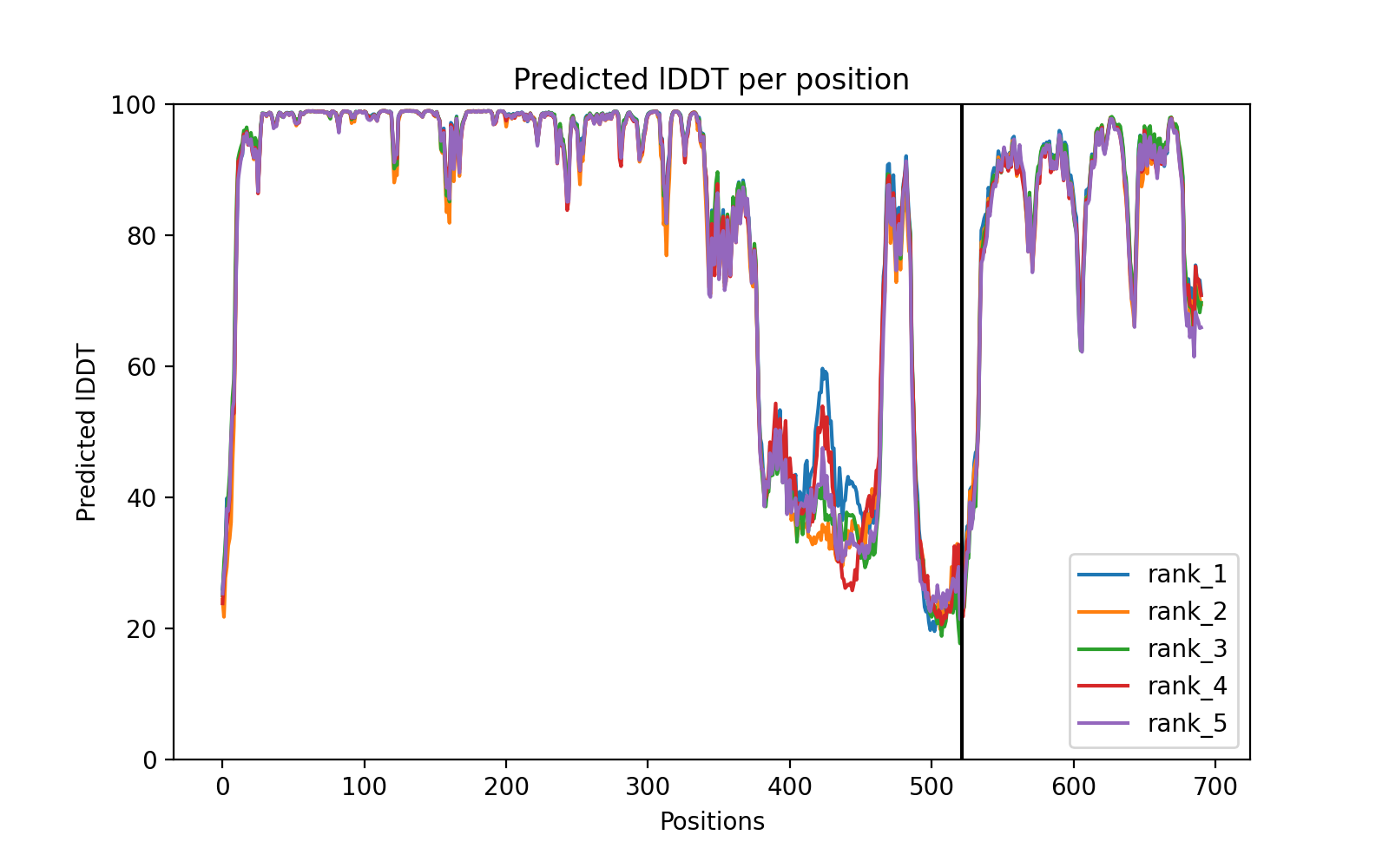

### PSMD5_SLC22A15_PAE.png

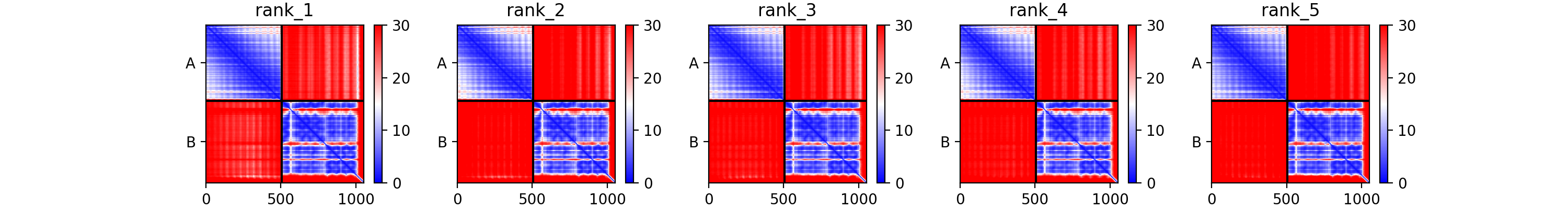

### PSMD12_CRIPT_PAE.png

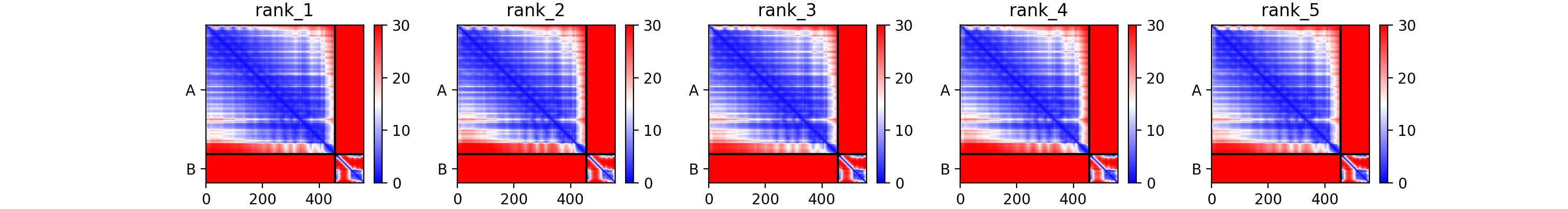

### RCC1_KLHL6_coverage.png

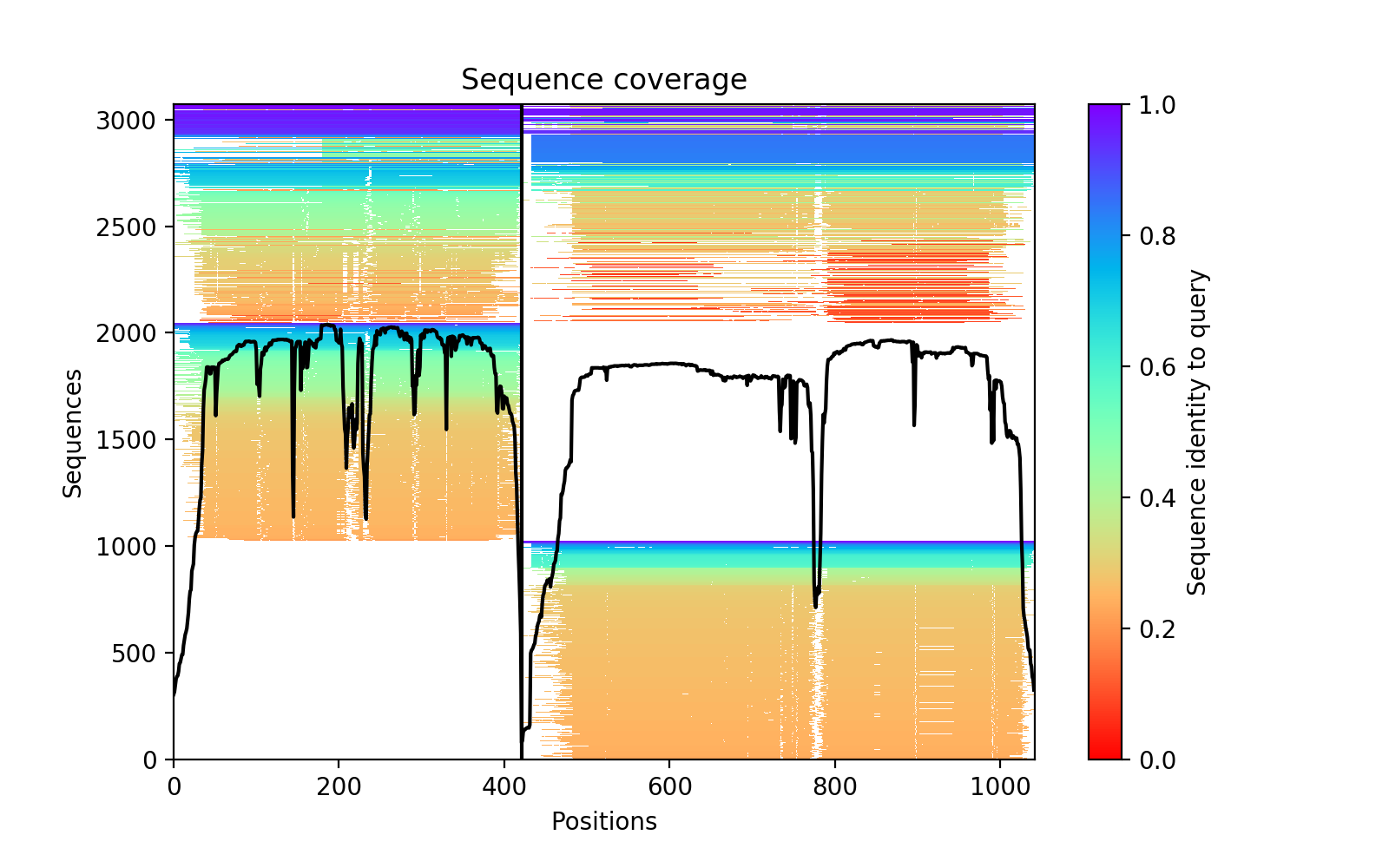

### RCC1_KLHL6_PAE.png

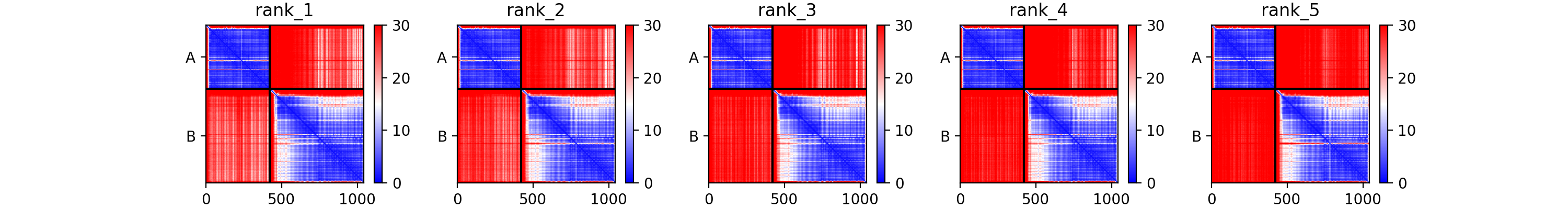

### RCC1_RAN_coverage.png

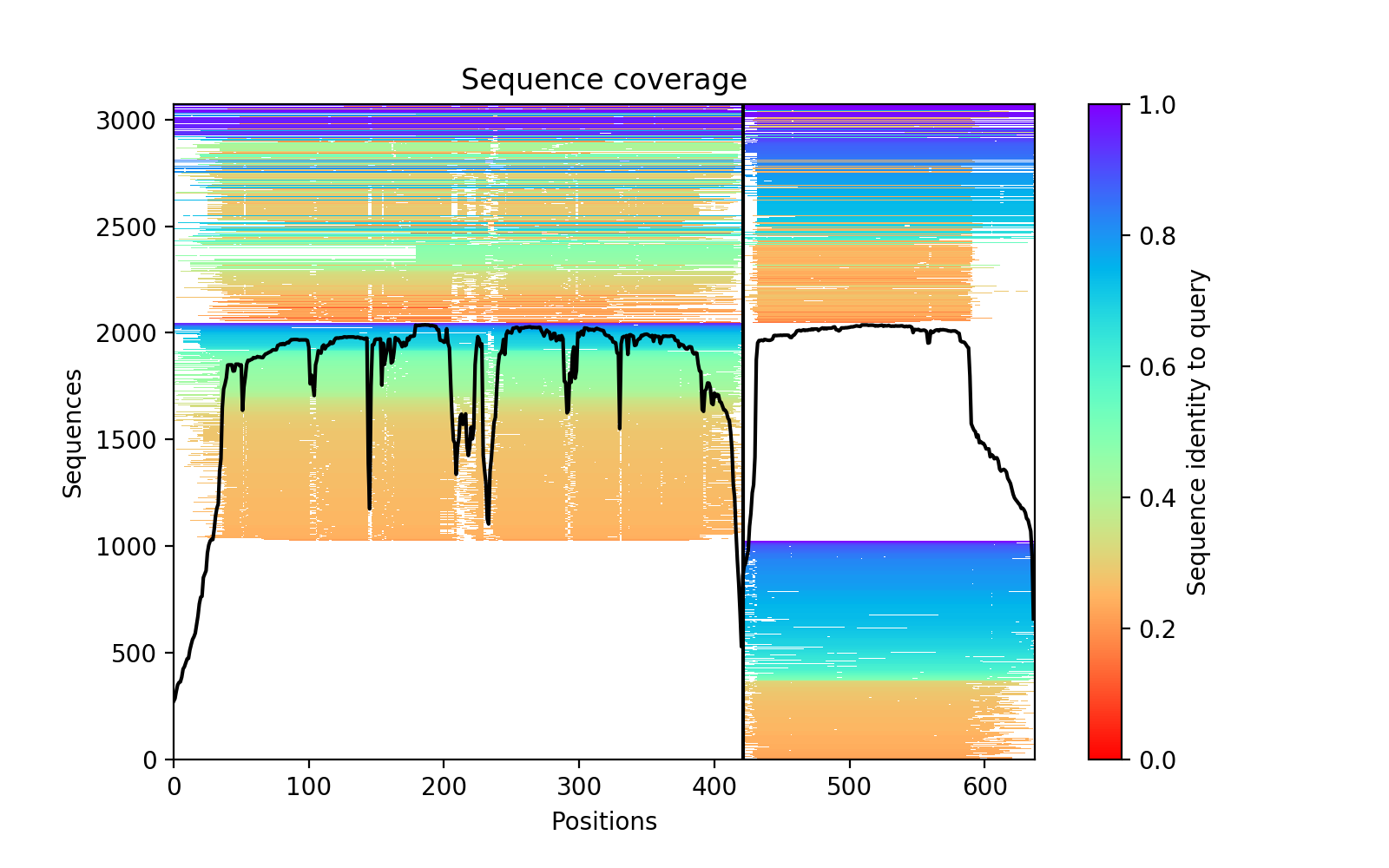

### RGR_ABCF3_coverage.png

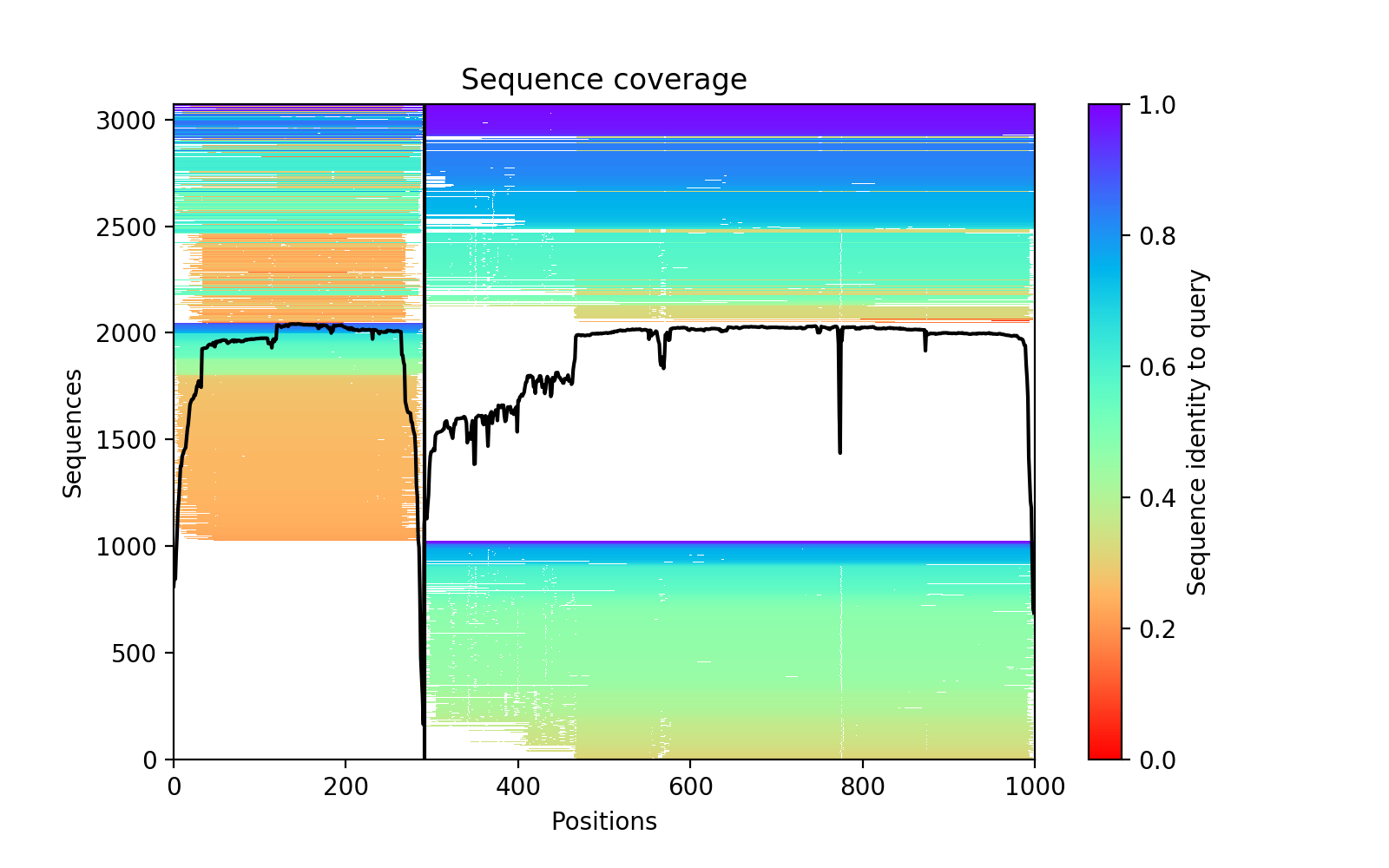

### SHMT2_STAC3_PAE.png

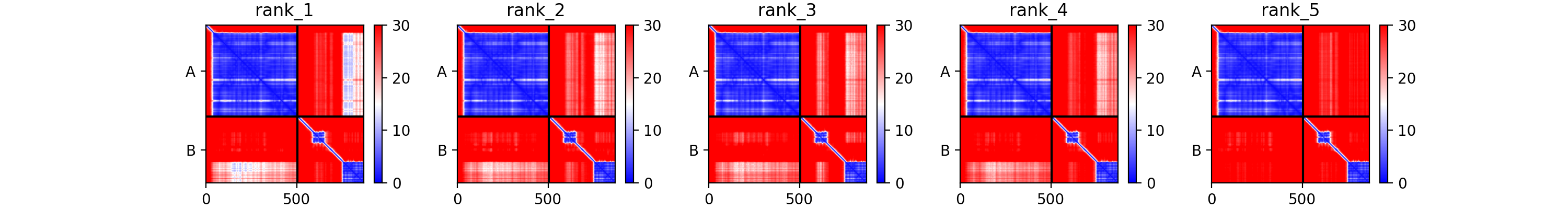

### SKP1_BTRC_PAE.png

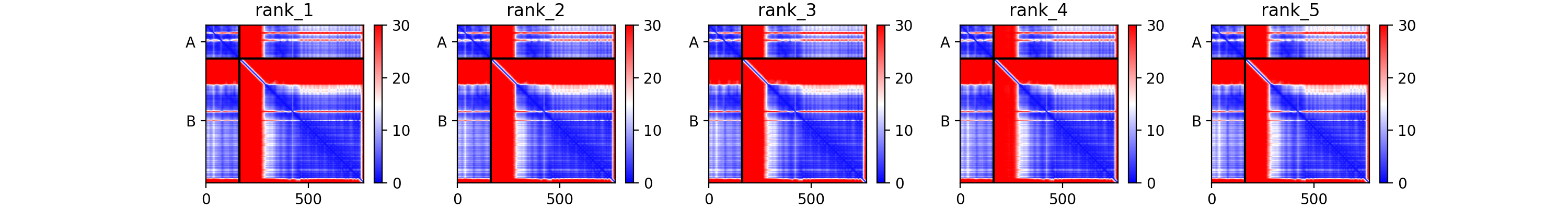

### SLC6A1_TM4SF4_plddt.png

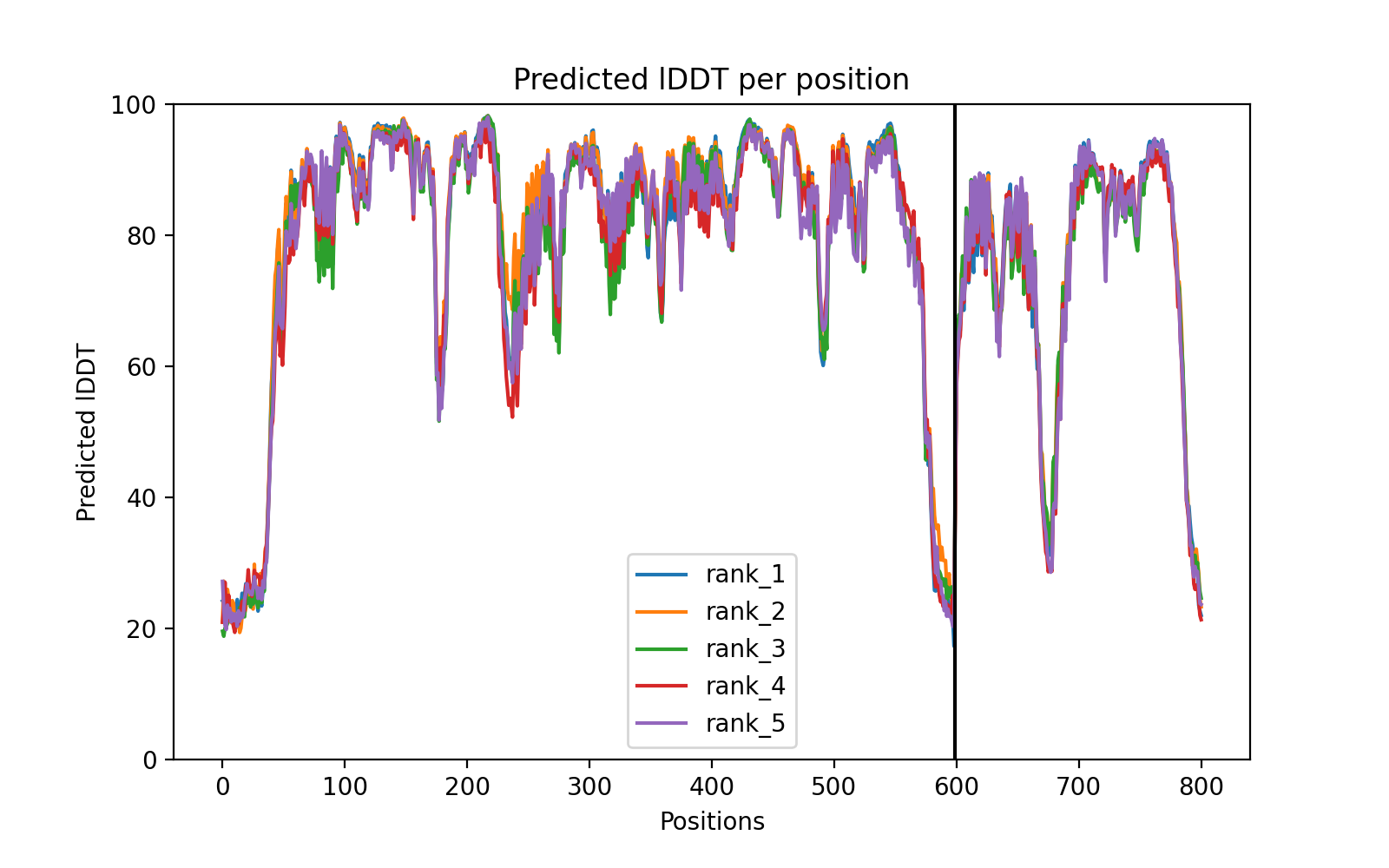

### SMAD1_SMAD4_coverage.png

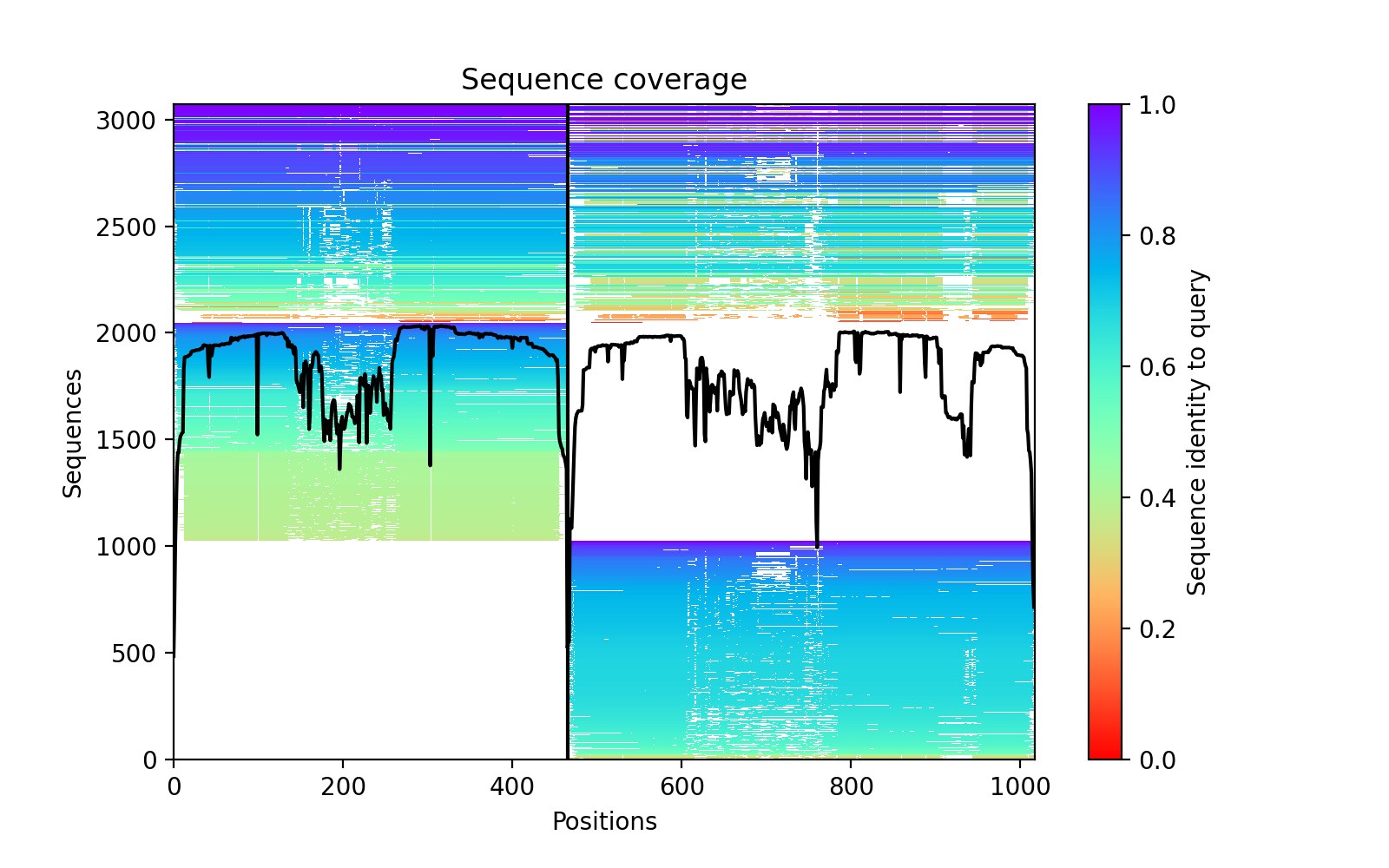

### SMAD4_DCP1A_PAE.png

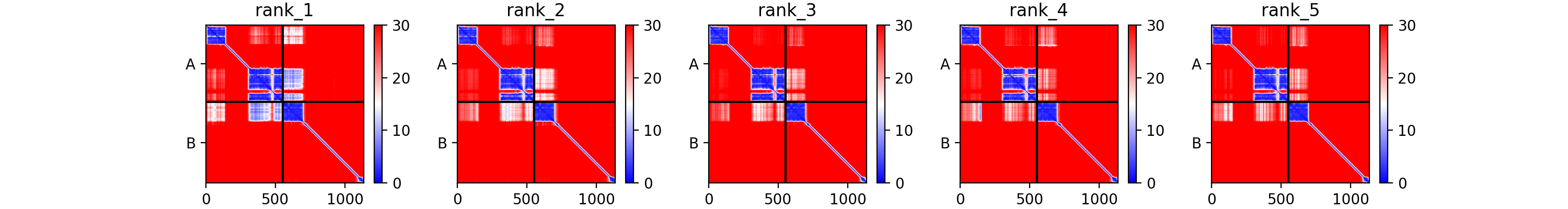

### XIAP_CASP7_plddt.png

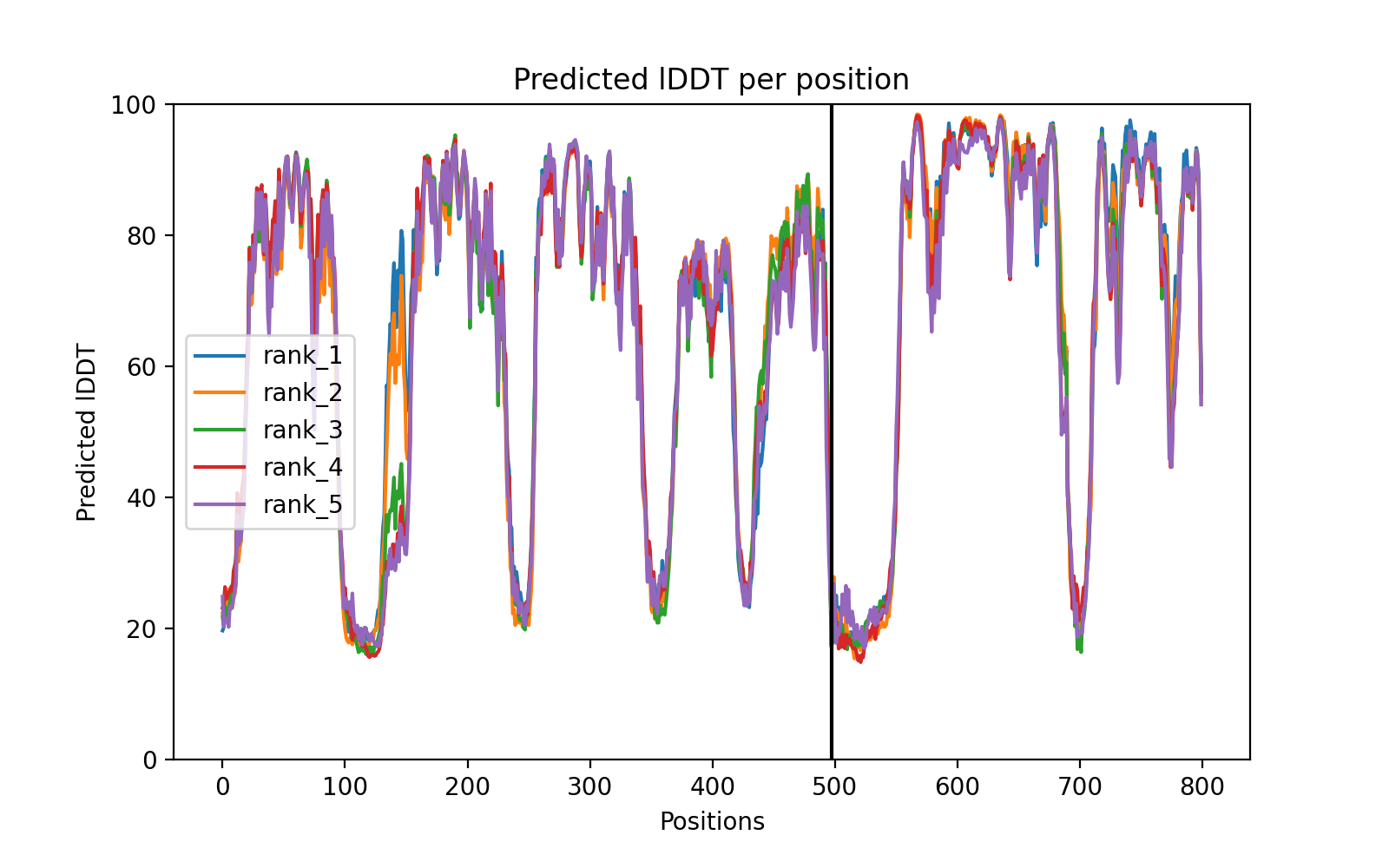

### XIAP_CASP9_PAE.png

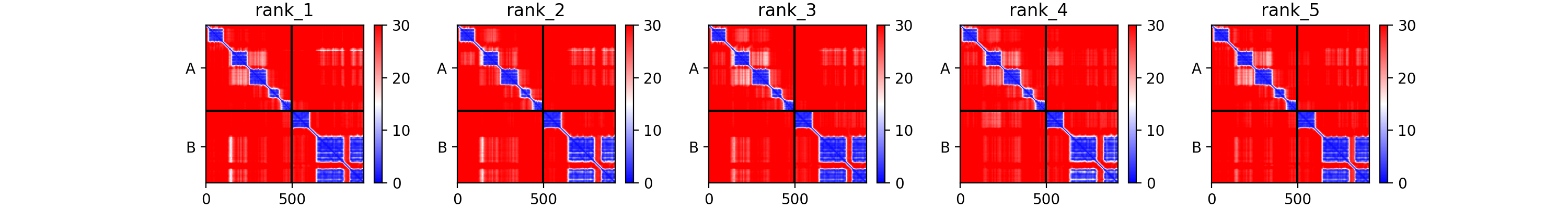
