## Supplementary figures and images for "AI-guided pipeline for protein-protein interaction drug discovery identifies a SARS-CoV-2 inhibitor"

### BAK1_BCL2L1_PAE.png

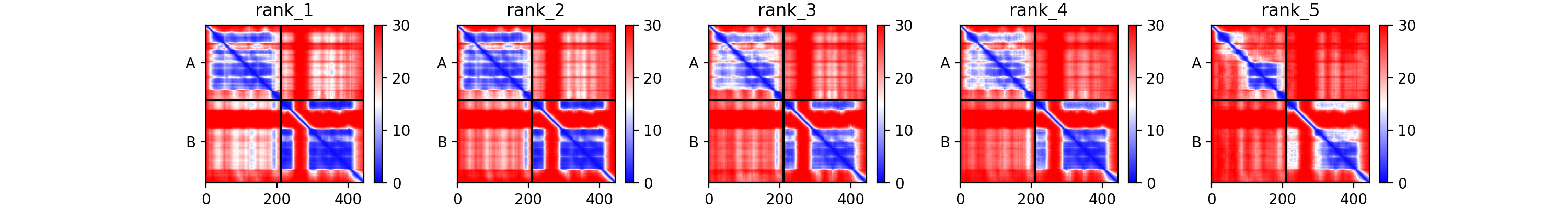
